# Uncovering Functional Distant Mutations by Ultra-High-Throughput Screening of Dehalogenases

**DOI:** 10.64898/2026.03.24.713925

**Authors:** H. Faldynova, D. Kovar, A. Jain, M. Slanska, M. Martinek, A. Jakob, M. Sulova, M. Vasina, J. Planas-Iglesias, S. M. Marques, N. Verma, P. Vanacek, D. Damborsky, C. P. S. Badenhorst, T. Buryska, F. W. Y. Chiu, M. Majerova, T. Kohutekova, P. Kouba, N. Sendlerova, A. J. deMello, J. Damborsky, J. Sivic, U. T. Bornscheuer, D. Bednar, S. Mazurenko, L. Hernychova, M. Marek, P. Klan, S. Stavrakis, Z. Prokop

## Abstract

Conformational dynamics play a central role in enzyme function by controlling substrate access and productive binding. Yet mutations that beneficially modulate these properties are difficult to identify. Here, we used ultrahigh-throughput fluorescence-activated droplet sorting (FADS) with a bulky fluorogenic substrate derived from coumarin (COU-3) to impose steric selection pressure on the haloalkane dehalogenase LinB. Screening a focused library yielded five single substitutions located 11.5–15.5 Å from the catalytic centre. Variant I138N showed a fourfold increase in catalytic efficiency toward COU-3 through reduced *K*_M_ and increased *k*_cat_, associated with increased cap-domain flexibility and facilitated substrate entry. In contrast, variant P208S markedly reduced substrate inhibition and shifted specificity toward bulkier iodinated haloalkanes by reshaping its tunnel environment. Integrated kinetic and structural analyses revealed that screening with bulky substrates directs selection toward distal regions controlling substrate access and unproductive binding. These findings demonstrate that ultrahigh-throughput FADS can reveal dynamic mechanisms of enzyme adaptation that remain difficult to predict by rational design.

**GRAPHICAL ABSTRACT:** 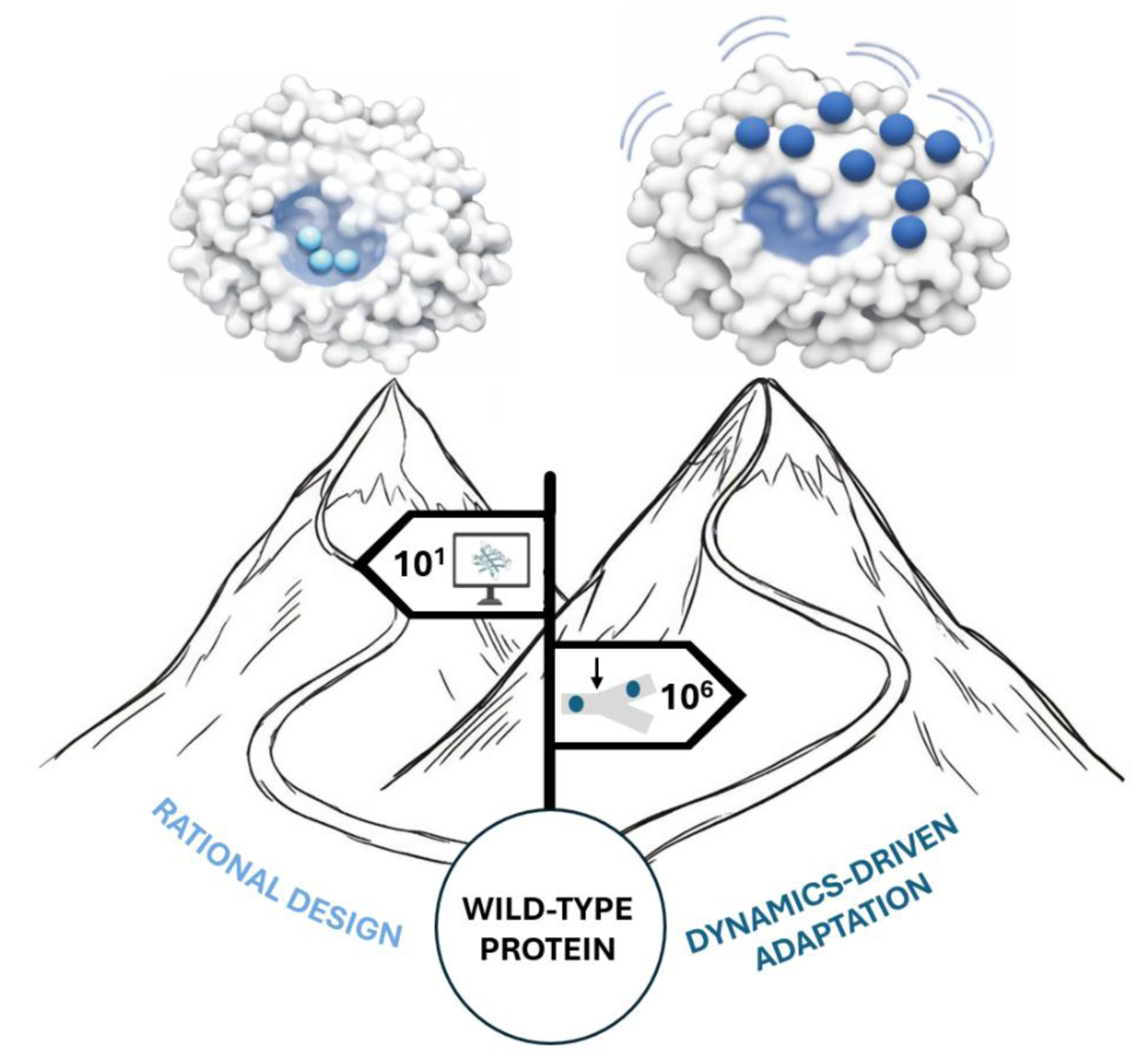

## INTRODUCTION

Enzymes with high specificity and selectivity are essential for industrial, environmental, and biomedical applications. Even so, natural wild-type enzymes often require extensive engineering before achieving the desired performance with non-native substrates. Enzyme activity is often limited not only by the active site but also by conformational dynamics and the architecture of the access tunnel, which govern substrate acquisition and productive complex formation. Such features are difficult to redesign rationally, as mutations affecting flexibility, gating, or substrate transitions are frequently located further from the active site^1^. As a result, they are hard to identify from sequence conservation or using standard computational tools^2^. Consequently, engineering enzyme specificity often demands large, high-diversity libraries that far exceed the screening capacities of conventional assays^3,4^.

Microfluidic systems offer a direct solution to these requirements^5–7^. Their low reagent consumption, precise control of reaction environments, and flexible platform design have enabled a wide range of high-throughput biochemical measurements^8^. Fluorescence-activated droplet sorting (FADS) has proven powerful for selecting promising enzyme variants from large libraries (10^6^–10^8^ variants). By operating at kilohertz frequencies, FADS enables the real-time analysis and selection of thousands of droplets per second based on their fluorescent signatures ^9–14^. Given the massive throughput of such systems, the likelihood of discovering rare beneficial yet remote or allosteric mutations increases dramatically.

In this study, we employed FADS to screen a model enzyme, the haloalkane dehalogenase (HLD) LinB, for which extensive structural and mechanistic data indicate that the dynamics of the evolutionarily variable cap domain are critical for catalytic function and substrate specificity^15–17^. Building on this, a focused LinB cap-domain library was constructed, and a bulky fluorogenic substrate, COU-3, was designed to exceed the size of conventional haloalkane substrates, thereby requiring increased conformational flexibility and adaptation of the cap domain during substrate entry. The selected HLD variant, LinB from the *Sphingobium* genus, has a buried active site accessible via multiple tunnels, making it an ideal system for studying how protein dynamics and distant mutations influence substrate access and specificity. Previous work has shown that mutations in the cap domain or access tunnels can reshape substrate specificity, yet the effects of remote residues remain difficult to predict^18,19^. We therefore hypothesised that screening a cap-domain library under sub-saturating concentrations of a bulky substrate would preferentially recover mutations that improve substrate acquisition or suppress non-productive binding modes.

In this work, we screened a focused LinB cap-domain library against COU-3 using FADS, identifying variants that consistently displayed reduced apparent *K*_M_. These hits featured two distal mutations with distinct functional outcomes: one exhibiting improved catalytic efficiency toward COU-3, and the other showing enhanced specificity for smaller iodinated haloalkanes and reduced substrate inhibition. To rationalise these phenotypes, we combined steady-state kinetics, substrate-specificity profiling, molecular dynamics (MD) simulations, hydrogen–deuterium exchange mass spectrometry (HDX-MS), and machine-learning-based flexibility predictions (**Figure 1**). Together, these approaches revealed mutation-dependent effects on cap-domain dynamics, tunnel hydration, and substrate access that are not evident from static structures, illustrating how ultra-high-throughput droplet-based screening can uncover new dynamic structure–function relationships that remain inaccessible to current rational design strategies.

**Figure 1:**
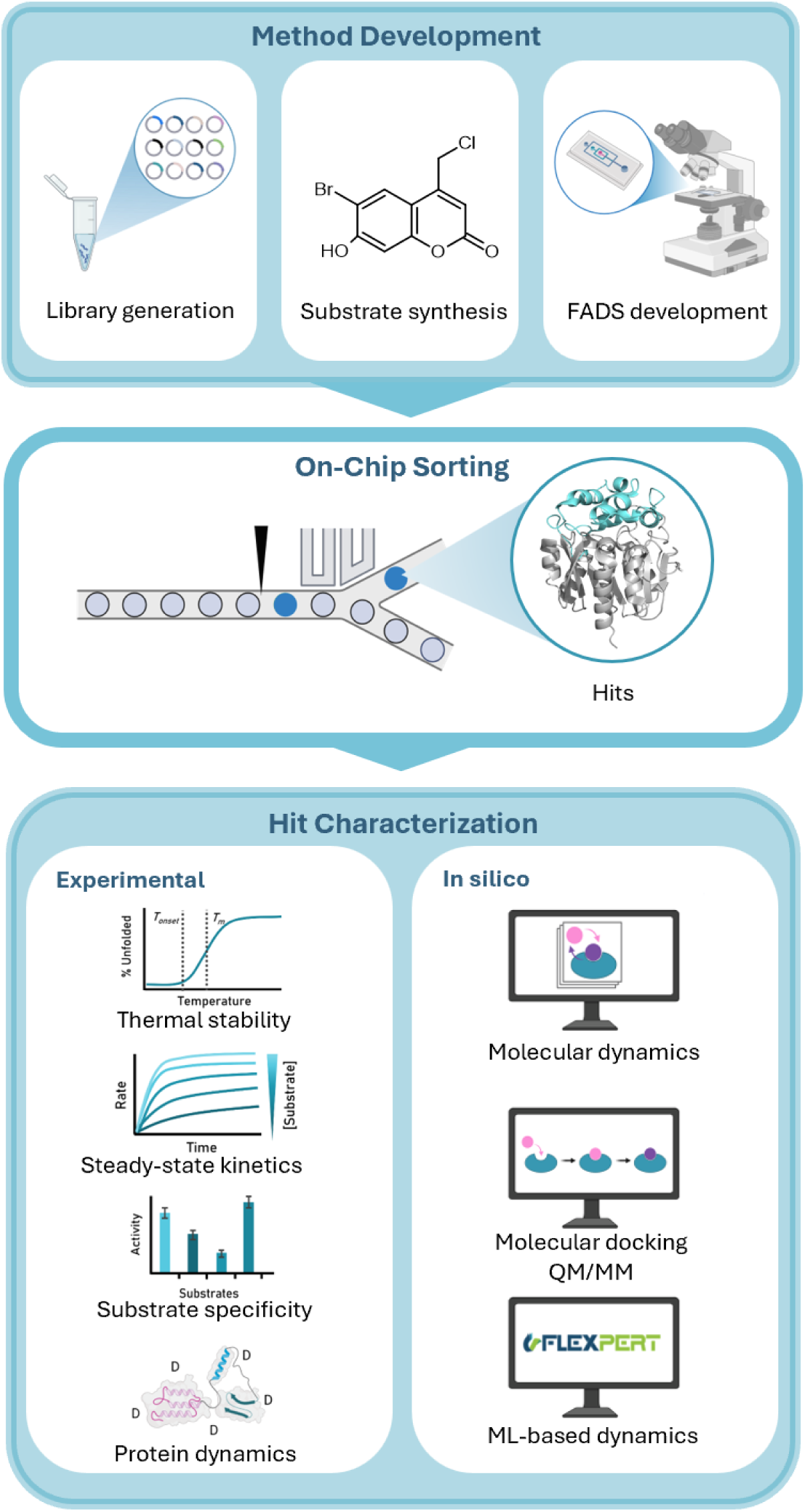
Integrated droplet-based screening and mechanistic analysis of LinB. **Method development** consisted of the construction of a focused substitution library targeting the LinB cap domain, synthesis and photophysical characterisation of the bulky fluorogenic substrate COU-3, and optimisation of a custom-built FADS platform for single-cell, single-droplet enzymatic activity readout. Followingly, **on-chip sorting** of the individual cells was performed, with expressed protein variants co-encapsulated with COU-3 in droplets, incubated, and interrogated by fluorescence. Droplets exceeding a defined activity threshold are dielectrophoretically sorted into the positive channel (dark blue droplet), enriching variants with improved performance under limiting-substrate conditions. Lastly, hit characterization took place as the sorted variants were recovered, sequenced, and subjected to comprehensive biochemical and biophysical analysis, including thermal stability measurements, kinetic profiling, substrate-specificity mapping, and HDX-MS with complementary molecular-dynamics (MD), Docking, Quantum Mechanics/Molecular Mechanics (QM/MM), and ML-based annotations to dissect the mechanistic origins of the observed functional gains.

## RESULTS

### Fluorogenic substrate COU-3 enables selective detection of LinB activity in droplets

To enable fluorescence-based ultrahigh-throughput screening, we synthesised coumarin-based analogues (**Table S2**). The designs consisted of sterically demanding haloalkane bearing a coumarin backbone to shift the selective pressure towards substrate access, tunnel opening, and changes in cap domain dynamics. These bulky substrates were initially evaluated with DmmA, an HLD known to possess a broad substrate range and a large active-site cavity^20,21^ (**SI Section 2.7**), to validate their solubility, sensitivity, and compatibility with efficient dehalogenation. Subsequently, a stable, water-soluble COU-3 was used to detect LinB activity by releasing a strongly fluorescent product upon HLD-catalysed dehalogenation (**Figure 2A** and **2B**). The compound exhibits well-resolved excitation and emission maxima (λ_ex_ 366 nm, λ_em_ 457 nm) and a substantial increase in fluorescence intensity upon enzymatic conversion, providing a high dynamic range suitable for single-droplet readout. Spectroscopic analysis (**Figure 2B**) verified minimal background fluorescence from COU-3 and robust emission of the dehalogenation product under screening conditions with an approximately 10-fold increase in fluorescence signal (product vs. substrate) that ensures reliable discrimination between active and inactive variants within five-picolitre droplets.

**Figure 2:**
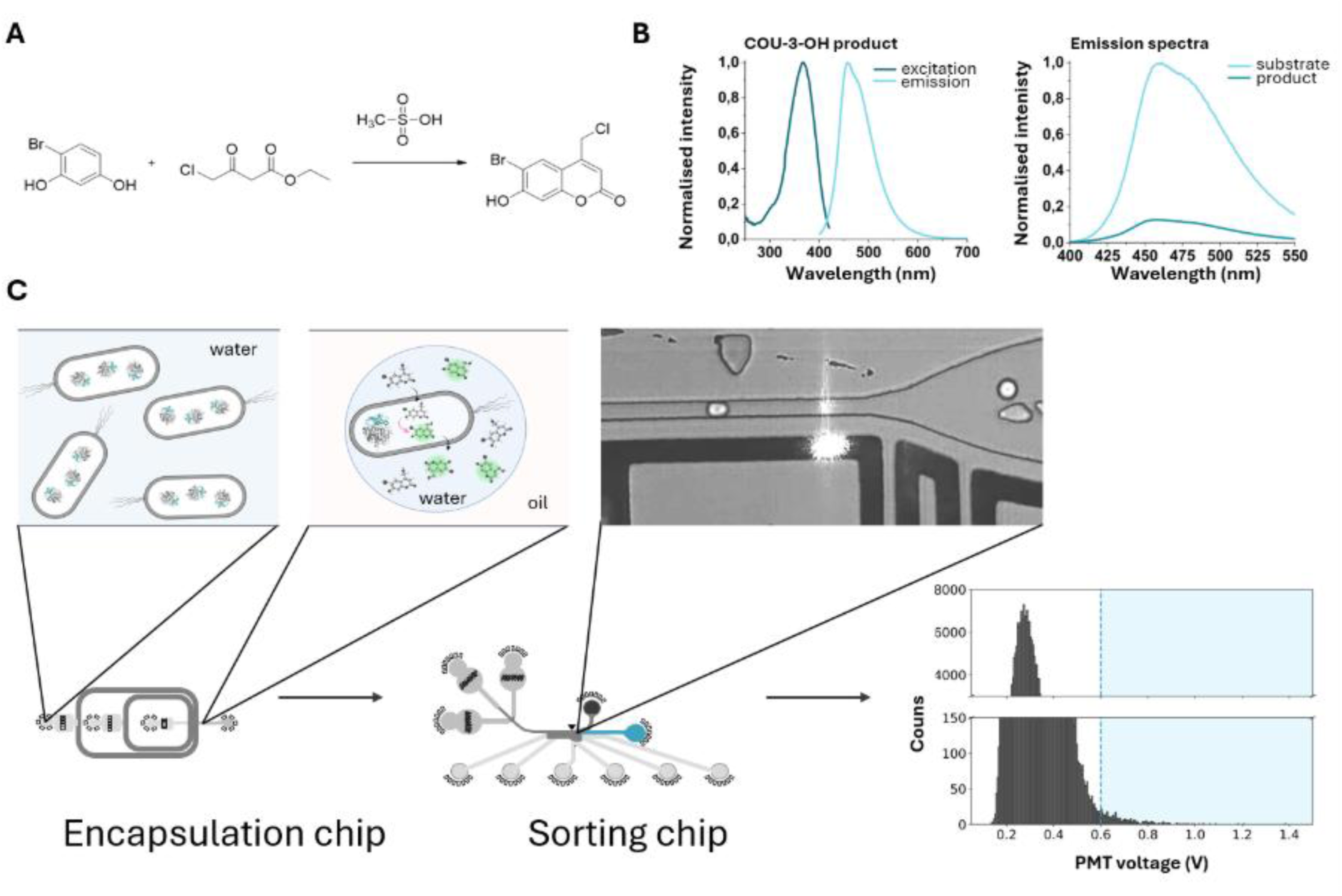
COU-3 Properties and FADS Screening Workflow. **A**) Synthesis of the fluorogenic haloalkane analogue COU-3. **B**) Photophysical characterisation of COU-3 and its product. COU-3 displays excitation and emission maxima at 366 nm and 457 nm, respectively. Enzymatic conversion results in a 10-fold increase in fluorescence. **C**) Schematic of the droplet-based screening workflow. Individual *E. coli* cells expressing LinB variants are co-encapsulated with COU-3 in droplets. After incubation, droplets pass through the detection region of the FADS sorter, where fluorescence intensity is recorded in real-time. Droplets exceeding the activity threshold are deflected into the positive channel, while low-signal droplets are discarded. The representative fluorescence histogram shows clear separation of the most active droplets.

### Droplet encapsulation and sorting enable quantitative selection of active variants

The screening workflow is illustrated in **Figure 2C**. Single *Escherichia coli* (*E. coli*) cells expressing individual LinB variants were co-encapsulated with COU-3 (the final concentration of 50 µM) in water-in-oil droplets, where the substrate readily partitioned across the cell membrane, allowing intracellular conversion and the subsequent accumulation of the fluorescent product. Droplets were then sorted based on real-time fluorescence readouts. The resulting fluorescence histogram demonstrates the threshold used to separate the non- or low-active populations from the active variants, validating the assay’s sensitivity and dynamic resolution.

Droplets exceeding the activity threshold were collected, and the corresponding genetic material was recovered to identify individual LinB variants, which were subsequently produced, purified, and subjected to biochemical and structural characterisation. Crucially, the COU-3 concentration used for screening was well below the apparent *K*_M_ of wild-type LinB toward this substrate (∼174 µM), yet above that of the positive-control DmmA (∼15 µM), which possesses a naturally enlarged access pathway for bulky substrates. Thus, the use of undersaturated substrate concentrations within the droplets imposes strong selection pressure on variants that improve substrate acquisition, tunnel opening, or productive Michaelis complex formation, rather than on catalytic turnover alone. This screening regime is consistent with the systematic recovery of variants displaying reduced apparent *K*_M_ values in the FADS experiments.

### Identified Mutations Exhibited Kinetic and Distance Trends

Screening of the cap-domain focused substitution library using the FADS platform yielded five single-point LinB mutants: P137S, I138N, V173F, P208S, and R209L. Structural mapping (**Figure 3A**) confirmed that all substitutions cluster within the third shell (between 8 and 12Å from the catalytic pentad centre of mass) or reside further away on the cap domain surface and the tunnel-access regions, rather than within a closer range of the catalytic pentad (between 5 and 8Å in the second shell or closer than 5Å in the first shell). This highlights the role of distal residues in modulating activity.

**Figure 3:**
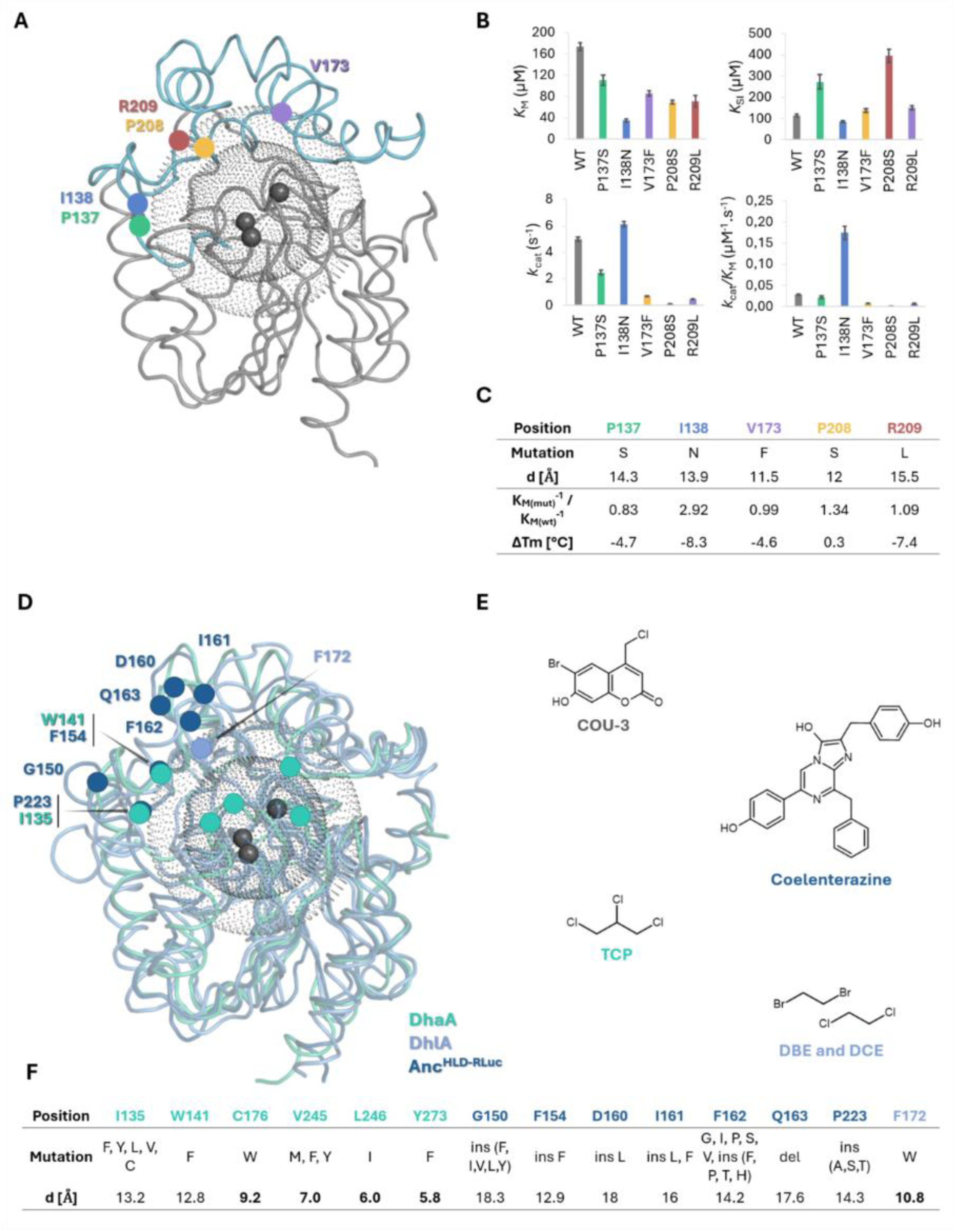
Identification and Kinetic Characterisation of Distant Mutations. **A)** Structural mapping of all five mutations (P137S, I138N, V173F, P208S, R209L), located on the cap domain surface or tunnel-access regions. The residues shaping the active site are shown in black; the shells shown have the radius from the catalytic centre of 8Å (black) and 12Å (grey). **B)** Kinetic analysis of LinB wild-type and identified mutants, based on the Michaelis–Menten fits with COU-3, shows that all variants exhibit reduced *K*_M_. Variant I138N shows a 4-fold improvement in catalytic efficiency, while P208S uniquely exhibits 4-fold greater tolerance to substrate inhibition. **C)** Summary of kinetic parameters, mutation types, distances to the active site, and melting temperature changes. All mutations are located close to the surface of the protein, with 11.5-15.5 Å distance from the active site. **D)** Structural comparison with previously engineered haloalkane dehalogenases (green DhaA, light blue DhlA, and dark blue the reconstructed, bifunctional ancestral protein Anc^HLD-RLuc^). **E)** Substrate panels used in ours and previous studies (grey COU-3 for LinB, dark blue Coelenterazine for Anc^HLD-RLuc^ (luciferase activity), green TCP for DhaA, and light blue DBE with DCE for DhlA). **F)** Compilation of mutation positions from prior engineering campaigns, showing that mutations improving reactivity toward small substrates occur closer to the catalytic centre (5.8–13.2 Å), whereas mutations enabling turnover of bulky substrates arise more distally (12.9–18.3 Å).

Kinetic analysis of the isolated variants (**Figures 3B** and **3C**) revealed a strikingly consistent trend: all mutants exhibited a reduction in *K*_M_ toward the bulky substrate COU-3. This systematic decrease in *K*_M_ across the identified set of distal mutations further suggests that the screening conditions preferentially enriched variants with improved substrate acquisition or altered access-tunnel dynamics, rather than changes in the catalytic turnover (*k*_cat_) alone. Thermal unfolding measurements (**Figure 3C**) further differentiated the identified variants. While some substitutions had only minor effects on protein stability, two variants exhibited a decrease in melting temperatures (*T*_m_) of more than 5 °C relative to the wild-type. These results highlight the stability-activity trade-off frequently observed in directed evolution, suggesting that these mutations achieve enhanced performance through distinct structural and dynamic mechanisms.

### I138N Enhanced Catalytic Efficiency, P208S Increased Tolerance to Substrate Inhibition

Two variants emerged with clearly distinct mechanistic signatures. The variant I138N displayed a pronounced enhancement in catalytic performance. In addition to its reduced *K*_M_, this substitution provided a measurable increase in *k*_cat_, resulting in a fourfold improvement in catalytic efficiency (*k*_cat_ / *K*_M_ increased from 0.04 to 0.17 s⁻¹ μM⁻¹). Despite being located 13.9 Å from the catalytic centre, I138N appears to modulate substrate engagement in a way that enhances overall reaction throughput. In contrast, the P208S variant exhibited an alternative functional gain. While its catalytic efficiency toward COU-3 remained unchanged, P208S exhibited a substantial increase in tolerance to substrate inhibition, with *K*_SI_ rising nearly fourfold from 110 μM to ∼ 400 μM. This phenotype indicates that P208S is less susceptible to unproductive substrate binding at high coumarin concentrations.

### Positions Mirror Evolutionary Patterns Seen for Other HLDs

Importantly, all five substitutions were located exclusively outside the active site, positioned 11.5–15.5 Å from the catalytic centre (**Figure 3C**). To contextualise these findings, we compared the mutational landscape uncovered here with mutations previously reported for four of the most common HLD targets of biotechnological relevance: 1,2,3-Trichloropropane (TCP), 1,2-Dibromoethane (DBE), 1,2-Dichloroethane (DCE), and Coelenterazine on DhaA^22^, DhlA^23^, and an ancestral HLD^24^ (**Figures 3D – 3F**). For small haloalkane substrates (TCP, DBE, and DCE), beneficial mutations systematically cluster closer to the catalytic centre (5.8–13.2 Å), where they fine-tune active-site geometry or transition-state stabilisation. On the other hand, variants optimised for large, bulky substrates (coelenterazine) primarily involve residues on the protein surface or at tunnel entrances, at distances of 12.9–18.3 Å; regions associated with long-range dynamic control rather than catalytic chemistry. The positions identified in this study fall within this latter regime, suggesting that the efficient processing of bulky substrates depends on distal, dynamic regions of the protein rather than on first-shell active-site residues.

### Microfluidics-based Substrate Specificity Profiling

To assess whether the mutations identified in the FADS campaign influenced substrate scope beyond the bulky COU-3, we profiled the activity of all five variants against a panel of 27 non-bulky haloalkanes using the MicroPEX method^25^. Assays were performed at 25 °C, a temperature chosen based on the thermostability profiles of the variants (**Figure S27** and **Table S14**). We categorised the obtained activity data into three groups based on the leaving group of the substrate—chloride, bromide, or iodide—and expressed the specific activities relative to the wild-type enzyme (**Figure 4A**).

**Figure 4:**
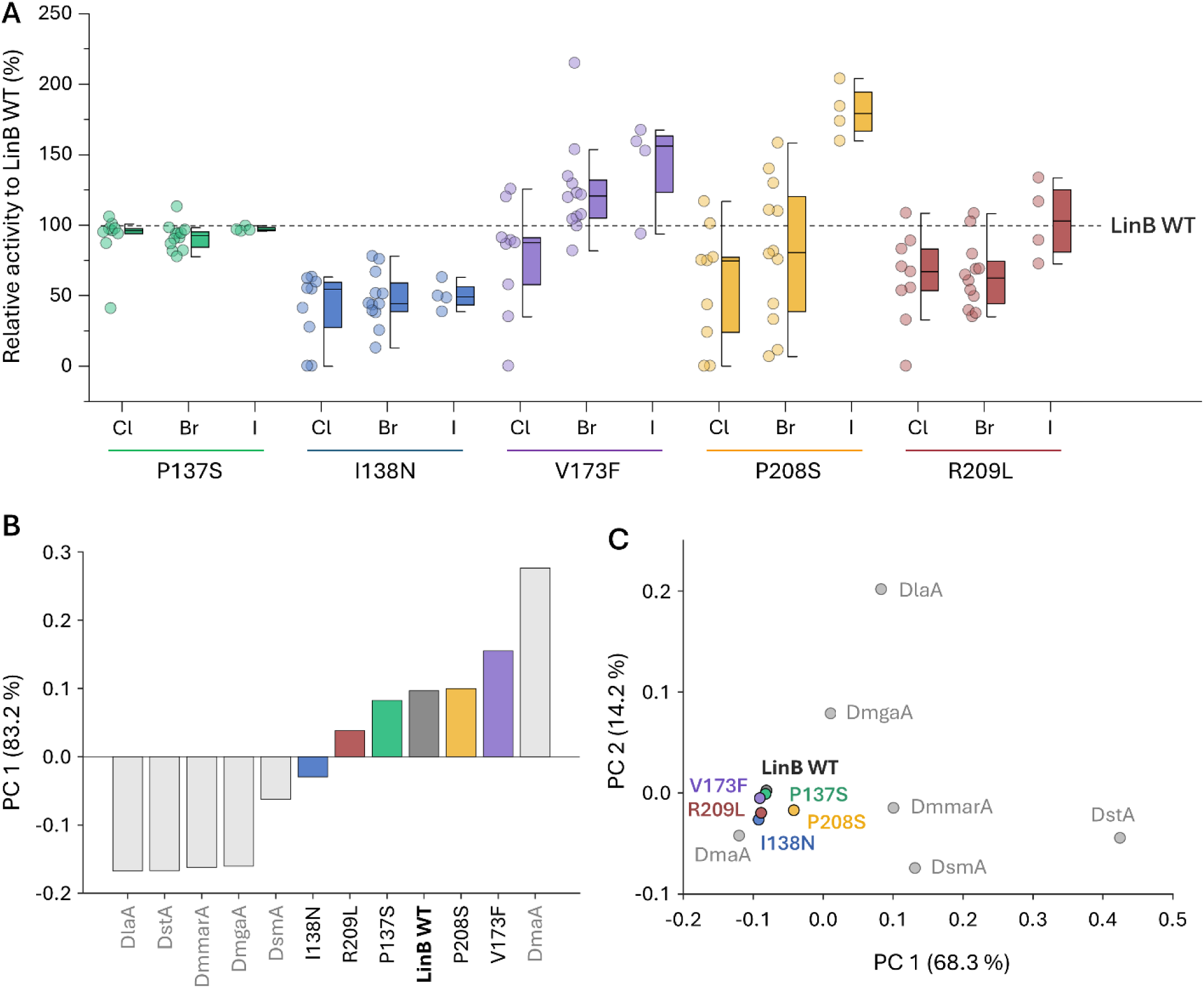
Substrate Specificity Characterisation. **A)** The box chart shows the relative specific activities of individual mutants with respect to LinB wild-type. For each mutant, the datapoints are divided into three subgroups, based on the leaving group (halogen atom) of the particular substrate (Cl: chlorine, Br: bromine, I: iodine). The dashed line represents the activities of LinB wild-type. The boxes show median (line), quartiles, minima, and maxima. **B)** The score plot from principal component analysis (PCA) was performed on absolute activity values. The chart shows the first principal component (PC 1), explaining 83.2% of the data variance terms, and ranks the variants based on their overall activity. LinB hits (colour-coded) are complemented by previously characterised wild-type HLDs^25^ (light grey), selected under identical measurement conditions to the LinB hits dataset. **C)** The score plot from PCA was performed on log-transformed activity values. The chart shows the first two principal components explaining a total of 82.5 % of the data. The graph shows the same variants as in **B)**.

Across the panel, the P137S variant showed activity profiles largely comparable to the wild-type, indicating only minor effects on specificity. In contrast, R209L and I138N exhibited reduced activity toward most small haloalkanes, with R209L showing a modestly increased preference towards iodinated substrates. This inverse relationship between performance on the bulky fluorogenic substrate and the smaller haloalkanes suggests that I138N promotes a functional trade-off: mutations that enhance turnover with large substrates appear to compromise the enzyme’s efficiency toward smaller ones. V173F demonstrated a distinct shift in specificity with selective activity improvements on most iodinated substrates. Notably, its strongest effect was observed with bromomethyl cyclohexane (∼200% of wild-type), indicating that this mutation might enhance reactivity with mid-sized hydrophobic substrates. The preference towards iodinated substrates was even more pronounced in P208S, which achieved 150–200% of wild-type activity across all four iodinated substrates. This selective gain mirrors the functional behaviour of P208S seen in its kinetic characterisation, which exhibited altered substrate interactions and increased tolerance to substrate inhibition without improving turnover toward COU-3 (**Figure 3B**).

To place the observed substrate specificities in the broader context of the haloalkane dehalogenase family with tens of biochemically characterised variants^25^, the principal component analysis (PCA; **Figure 4B** and **4C**) was applied following an established approach^26,27^. To ensure data consistency, the LinB dataset was augmented with six additional variants, for which substrate specificity data were measured under identical conditions, specifically at 25 °C^25^. The PCA performed on absolute activity values ranked the variants by their overall activity (**Figure 4B**). Notably, both V173F and P208S surpassed the LinB wild-type, while the remaining three hits showed reduced overall activities. A second PCA, performed on log-transformed data (**Figure 4C**), clustered all variants close to LinB wild-type and relatively apart from most other wild-type dehalogenases. The only variant that clustered near LinB variants was DmaA, due to its ability to maintain high activity down to 5 °C^25^. This clustering confirms that while the distal mutations fine-tuned the enzyme’s efficiency and access-tunnel dynamics, they preserved the core substrate-preference fingerprint characteristic of the LinB subfamily.

### Prediction of protein dynamics via Flexpert

Flexpert^28^ analysis was applied to assess how each substitution modulates residue-level conformational dynamics. The predicted log₂ (mutant/WT) flexibility profiles (**Figure 5A**) revealed that all five variants induce pronounced changes within the cap domain (residues ∼130-230). 3D mapping of the Flexpert output (**Figure 5C**) confirmed that the cap domain is the dominant dynamic region in wild-type LinB. Among all mutants, I138N exhibited the most pronounced increase in flexibility, with a distinct peak centred at the end of L9, a segment critical for shaping access tunnels p1 and p2. Mutations P137S and R209L were predicted with the most rigidified and dynamic regions within the cap domain, respectively, while flexibility changes on the variants V173F and P208S were estimated to be more moderate.

**Figure 5:**
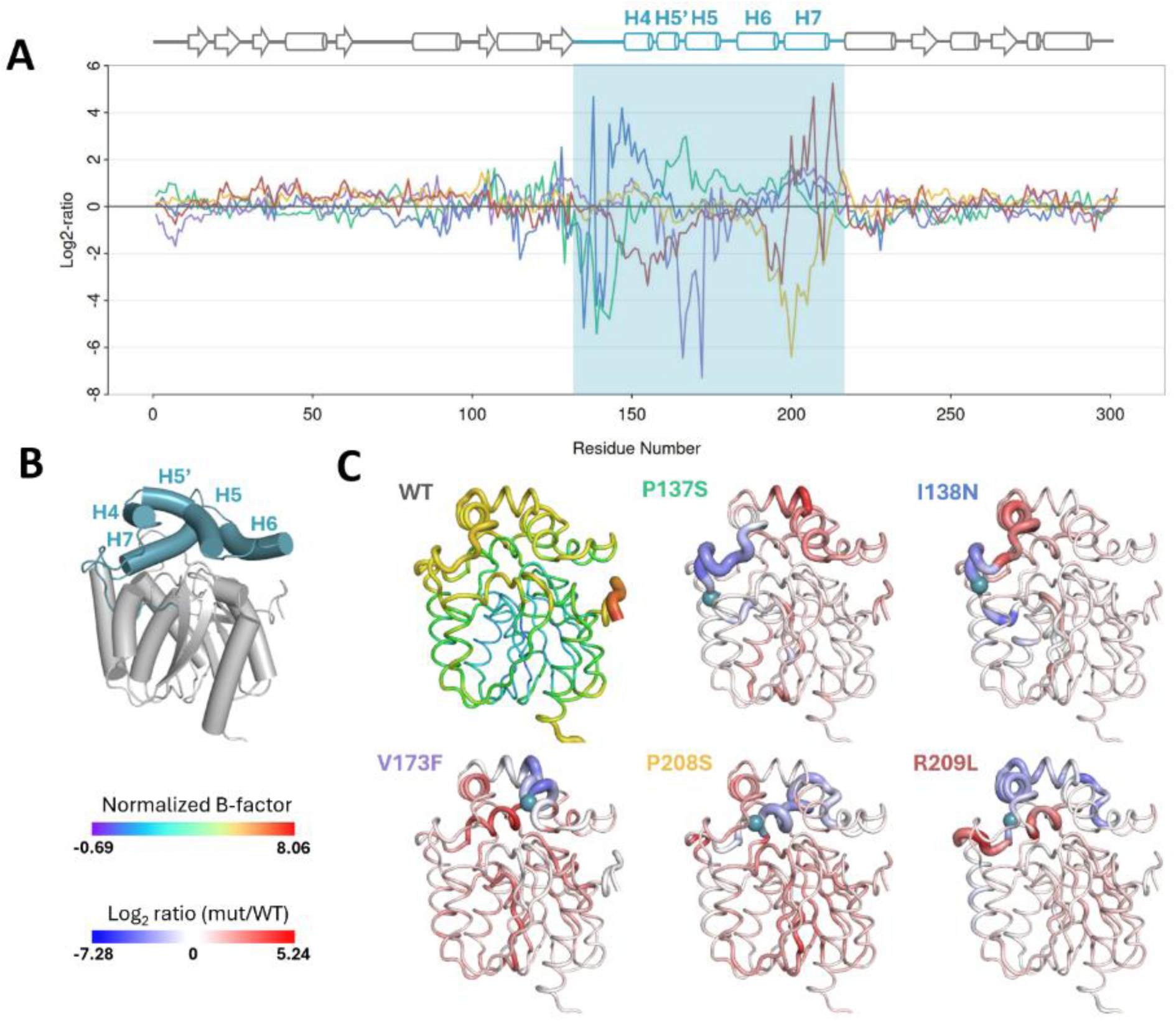
Analysis of ML-based Dynamics Prediction via Flexpert. **A)** log_2_ (mutant/WT) flexibility profiles across all 5 variants, highlighting mutation-induced changes in backbone mobility along the cap domain (turquoise range), which is visualised on the PDB structure 1mj5 in turquoise **B). C)** 3D mapping of normalised B-factors for the wild-type (showing inherently high flexibility in the cap domain) and log_2_ ratio maps for individual mutants with the colour legends on the bottom left. I138N exhibits pronounced flexibility increases in L9-H4 interface, while P208S and the remaining variants result in a minor increase or a notable decrease in that region.

### HDX-MS identifies distinct flexibility patterns in I138N and P208S

To determine how the two functionally distinct variants I138N and P208S differ in backbone dynamics, we performed HDX-MS experiments (**Figure 6A**). At the 10-second timescale, LinB wild-type and P208S showed nearly identical deuterium uptake profiles, indicating comparable global and local flexibility and suggesting that the P208S substitution does not substantially alter the intrinsic structural dynamics of the enzyme in its free state. In contrast, the I138N variant displayed consistently higher deuterium uptake across the cap domain (residues 130–230) and the downstream helices α9–α10, indicating increased solvent exposure and enhanced flexibility throughout these regions. Interestingly, this effect was global across the entire cap rather than being confined to a small region, reflecting a mutation-induced loosening of the cap architecture. Despite these global differences, one region - the L7/β5 segment (residues 100–110) - remained dynamically conserved across all three proteins, thus providing a robust internal reference for the structurally rigid core of the enzyme. Together, these observations align with the functional trade-off observed in the kinetic analysis: the improved activity of I138N toward the bulky COU-3 substrate is coupled with increased conformational flexibility and reduced structural stability, as reflected in its elevated deuterium uptake.

**Figure 6:**
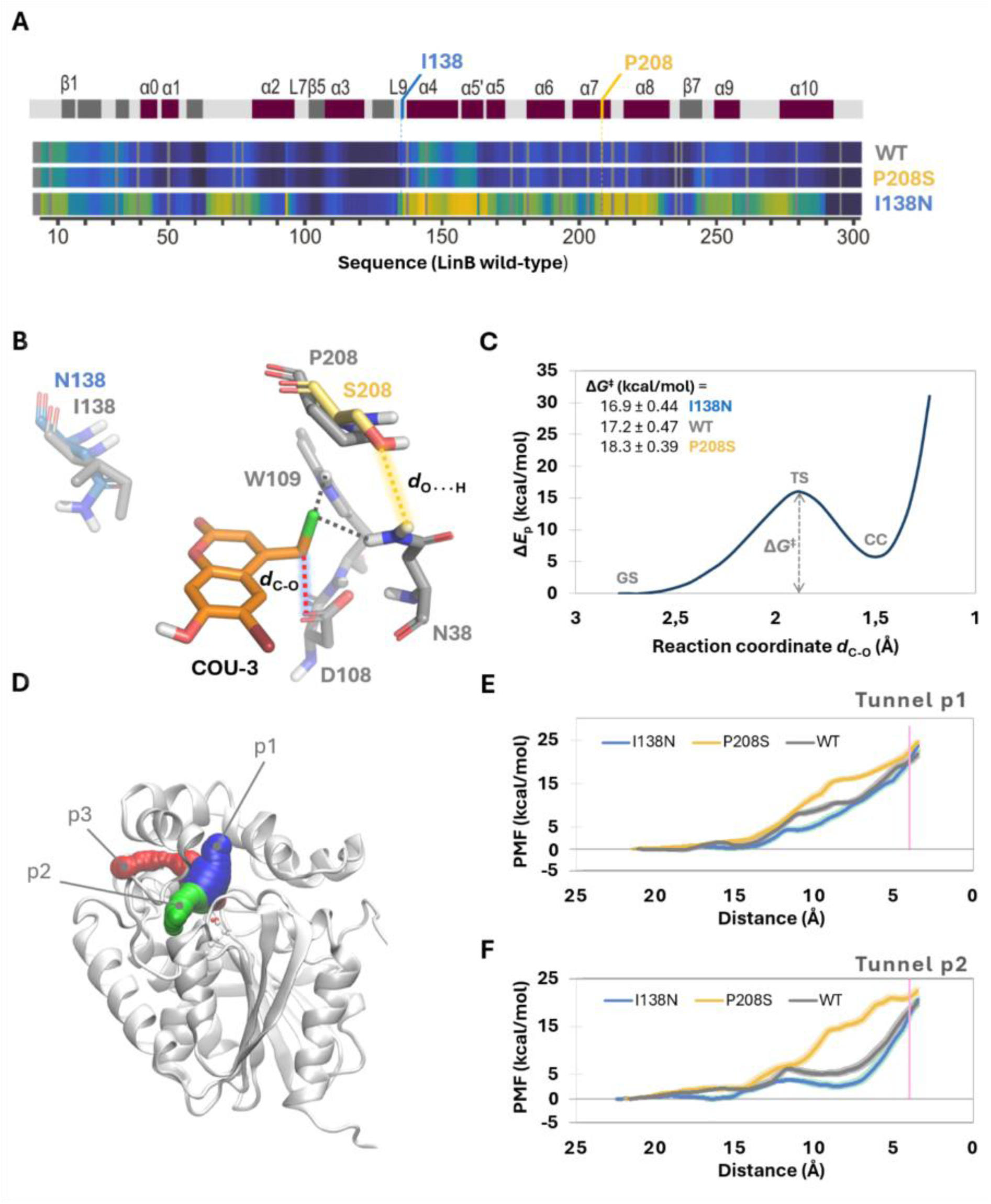
HDX-MS and MD Reveal Structural Basis for I138N and P208S Properties. **A)** HDX-MS deuterium-uptake profiles at 10 s. WT and P208S show similar global dynamics, whereas I138N exhibits increased uptake across residues 130–230 and α9–α10, indicating elevated cap-domain flexibility. **B)** NACs of COU-3 in the active site for wild-type, I138N, and P208S. P208S uniquely forms a hydrogen bond between S208 and N38, competing with the catalytic halide-stabilising interaction. **C)** S_N_2 reaction potential energy profiles showing the lowest activation barrier for I138N, followed by wild-type and P208S, mirroring measured catalytic efficiencies. The profile shows the ground state (GS) of the reactant complex, the transition state (TS), the free Gibbs energy (ΔG^‡^), and the chemical coordinate (CC). **D)** Structural representation of tunnels p1, p2, and p3 calculated using software CAVER. **E)** and **F)** Potential of Mean Force (PMF) profiles for ligand migration through tunnels p1 and p2. COU-3 consistently favours entry through tunnel p2, which provides a lower-energy binding route for all variants. The overall binding free energy follows the order I138N < WT < P208S, aligning with both S_N_2 reaction energetics and kinetic behaviour. The vertical pink line represents the final distance of 4 Å between the substrate reacting carbon and the Asp108-Cγ atom.

### Near-Attack Conformation (*NAC*) geometry reveals mutation-dependent interactions

To investigate the molecular basis for the observed kinetic shifts, COU-3 was docked into all three LinB variants. The docking revealed a conserved binding mode (**Fig. S31**) with nearly identical binding energies across all three proteins (**Table S17**). Therefore, MD simulations were used to examine the pre-reactive complexes, or near-attack conformations (*NAC*s) of COU-3 bound in the active sites of wild-type LinB and the two selected variants: I138N and P208S (**Figure 6B**). In all three enzymes, COU-3 occupies a similar position relative to the catalytic residues, allowing the formation of halide-stabilising interactions between the substrate chlorine atom and residues N38 and W109. However, the P208S substitution introduced a unique interaction, as the serine hydroxyl frequently forms a hydrogen bond with the second hydrogen of N38 (**Fig. S32**), an interaction that is absent in both wild-type and I138N. Because N38 is a key residue for stabilising the S_N_2 transition state through the N38-NH···Cl interaction, competition for its hydrogen-bonding capacity by S208 is predicted to decrease the transition-state stabilisation. This structural competition provides a robust mechanistic explanation for the reduced catalytic performance of P208S.

### S_N_2 reaction energetics correlate with experimental catalytic efficiencies

The *NAC* snapshots observed from the MD simulations were extracted to calculate the activation energy barrier (ΔG^‡^) for the S_N_2 chemical step, using QM/MM adiabatic mapping. This step involves the nucleophilic attack of the Asp108 carboxylate oxygen on the chloromethyl carbon of COU-3. The calculated potential energy profiles for the S_N_2 reaction (**Figure 6C**) revealed different activation energies for the three enzymes. The energy barrier was the lowest for I138N (16.9 ± 0.4 kcal/mol), followed by the wild-type (17.2 ± 0.5 kcal/mol), while P208S showed the highest barrier (18.3 ± 0.4 kcal/mol). This ranking in the energy barriers (I138N < WT < P208S) translates into the reverse order of the kinetic rates of the S_N_2 reaction, and it perfectly mirrors the trends observed experimentally in the *k*_cat_/*K*_M_ values (**Figures 3B** and **3C)**, indicating that changes in the *NAC* geometry translate into measurable differences in the chemical step. The elevated barrier in P208S is mechanistically consistent with the additional hydrogen-bond formed by S208, which destabilises the transition state as described above.

### Ligand-migration energetics reveal preferential COU-3 access via tunnel p2

Mapping of the access tunnels in MD simulations for all three variants identified the three canonical LinB pathways: p1, p2, and p3 (**Figures 6D and S33, and Table S18**). According to the tunnel properties (ratio of detected tunnels (≥0.9 Å), ratio of open tunnels (≥1.4 Å), average bottleneck radius, and priority; **Table S18**), p3 was deemed unlikely to support bulky ligand transport. For that reason, we studied the thermodynamics of entrance of the COU-3 substrate through the p1 and p2 using Adaptive Steered MD (ASMD), obtaining potentials of mean force (PMFs) for both pathways (**Figure 6E** and **6F**). The PMF profiles revealed a surprising trend: despite p1 appearing more "open" in ligand-free simulations (**Table S19**), tunnel p2 provided a significantly lower-energy pathway for COU-3 entry across all variants (**Figures 6E** and **6F** and **Table S19**).

The PMF profiles further indicate that COU-3 can induce local rearrangements that open tunnel p2, particularly within the region spanning L9 and α4 (residues 141–149). This region corresponds partly to the highly dynamic cap segment identified in HDX-MS measurements. When comparing enzyme variants, the total energetic cost for COU-3 to reach a bound state (defined here as *d*_C–O_ = 4 Å) through p2 was the lowest for I138N and the highest for P208S (**Figure 6F** and **Table S19**). These differences are aligned with the experimental catalytic efficiencies, and reinforce the effects of the S_N_2 reaction activation barriers discussed above. Together, these results clearly support an induced-fit mechanism for COU-3 binding into the studied enzymes and suggest that similar conformational gating may underlie the accommodation of other bulky substrates in HLDs.

## DISCUSSION

In this study, we combined droplet-based microfluidic sorting with a custom bulky substrate to investigate how protein dynamics and distal structural features govern catalysis and specificity in the model enzyme haloalkane dehalogenase LinB. Under screening conditions that favour turnover of a bulky substrate, we found that enzyme fitness was predominantly determined by substrate access and tunnel dynamics, rather than by changes in the chemical steps that take place in the active site. All variants isolated from the focused cap-domain library carried substitutions located 11.5–15.5 Å from the catalytic centre, corresponding to the outer-shell or surface positions. This spatial distribution contrasts with adaptations observed for small haloalkanes, where beneficial mutations typically cluster near the active site^29,30^. The fact that only distal mutations were identified under these conditions highlights the importance of allosteric networks in controlling substrate routing and binding energetics. Since such dynamic effects are difficult to predict from static structural models, they are easily missed by rational design strategies. In contrast, ultra-high-throughput screening complemented by machine-learning-based flexibility predictions and HDX-MS measurements enables the systematic exploration of mutational landscapes to identify unexpected adaptive solutions. Our findings align with trends that have been observed in the directed evolution of three *de novo* Kemp eliminases^31^, as well as multiple natural enzymes, including cytochrome p450 monooxygenases^32^, and β-lactamases^33^. These studies collectively demonstrate a fundamental principle: while active-site mutations predominantly affect chemical steps, distal mutations can enhance catalysis by modulating substrate binding and product release^22,34,35^.

Examination of individual LinB variants revealed distinct evolutionary trajectories emerging from the activity screening using a bulky COU-3 substrate. The I138N substitution increases cap-domain flexibility, lowering the energetic barriers for substrate access and S_N_2 activation. While this results in improved catalytic efficiency, the gain comes at the cost of reduced thermal stability and decreased catalytic efficiency on smaller substrates. In contrast, the P208S substitution maintains wild-type–like global dynamics yet reshapes the tunnel environment through localised structural perturbations. More specifically, it exhibits altered substrate specificity and reduced substrate inhibition, suggesting that localised structural perturbations can reshape tunnel environments and suppress unproductive binding without necessitating a global in^22,36,37^. Notably, P208S showed a preference for iodinated haloalkanes, which are larger, more polarizable and possess weaker C–I bonds. These properties lower the intrinsic S_N_2 activation barrier and make catalysis particularly sensitive to subtle changes in halide-stabilising interactions within the access tunnels. In this context, the modified hydrogen-bonding network in P208S likely creates a more favourable electrostatic environment that disproportionately accommodates the high polarizability of iodine. Consequently, the mutation acts as a selective gate, amplifying specificity toward larger halogens even in the absence of an increased catalytic turnover rate^38,39^, where increased conformational flexibility enhances access and catalysis for bulky substrates but compromises structural robustness. In contrast, P208S preserves thermal stability, consistent with a locally confined structural change in the access tunnels rather than a global destabilisation of the protein fold. Together, these observations suggest that local structural tuning of access tunnels can reshape substrate routing and inhibition behaviour independently of catalytic turnover.

In conclusion, our study demonstrates that screening toward large and chemically more complex substrates shifts adaptive pressure away from active-site residues toward distal regions that enhance catalysis by modulating enzyme flexibility and facilitating substrate binding. These findings align with previous work on haloalkane dehalogenases^24^, where engineering conformational behavior^29^ and access tunnels^22,40^ enabled the development of enzymes capable of efficient processing of bulky substrates. Importantly, our work reveals that the adaptation of HLDs toward bulky substrates relies on a complex interplay between distal residues and active-site geometry, where long-range dynamic shifts in the cap domain choreograph substrate access and transition-state stabilisation in ways that static structural models simply cannot predict. More broadly, this study highlights the power of droplet-based ultra-high-throughput evolution to uncover mechanistically diverse, dynamics-driven solutions that are difficult to achieve through rational enzyme design alone.

## METHODOLOGY

### Development of the FADS platform

A custom-built FADS platform, termed *Cinderella*, was developed for ultra-high-throughput screening of enzymatic activity at the single-cell level. The system integrates microfluidic droplet generation, laser-induced fluorescence detection, and dielectrophoretic sorting under FPGA-based real-time control. Monodisperse water-in-oil droplets containing individual *E. coli* cells were generated using PDMS flow-focusing chips and reinjected into a dedicated sorting device equipped with Field’s metal electrodes for fluorescence-triggered deflection.

Fluorescence excitation (405 nm) and PMT-based detection enabled sensitive discrimination of droplet populations. Sorting parameters and flow conditions were optimised to ensure stable droplet handling and precise gating. Custom LabVIEW- and Python-based software was used for real-time signal acquisition, peak filtering, and threshold-based sorting. Benchmarking with fluorescent dyes and mixed enzymatic populations demonstrated reliable separation and enrichment of rare, highly active variants.

### Chemical synthesis of the substrate COU-3

COU-3 was synthesised following a reported procedure^41^. Ethyl-4-chloroacetoacetate and 4-bromoresorcinol were coupled in methanesulfonic acid at 25 °C for 2.5 h. After quenching with ice–water, the precipitate was collected and recrystallised from ethyl acetate to yield COU-3 as a beige solid (39%). The structure was confirmed by ^1^H NMR (300 MHz, DMSO-d6) (**Figure S20**), consistent with the literature^41^. In phosphate buffer (pH 8.0), COU-3 showed λ_abs_ = 366 nm (ε = 6500 M⁻¹ cm⁻¹) and λ_em_ = 457 nm (Φ_fl = 0.32) (**Figure S12**).

### Generation of substitution libraries of the LinB cap domain

Substitution libraries of the LinB cap domain were constructed by error-prone PCR (GeneMorph II kit; 0.7–1.5 mutations/gene) and Gibson Assembly. The ∼340 bp cap region was amplified with GA-compatible primers, gel-purified, and assembled into BsrGI/XmaI-linearized pET21b (**Table S10**). Libraries were transformed into NEB^®^ 10-beta electrocompetent *E*. *coli* by electroporation, recovered in SOC medium, plated, and expanded overnight in Luria Broth (LB) medium with ampicillin for plasmid purification. Library integrity and mutation frequency were assessed by Sanger sequencing of randomly selected clones (**Figure S21 and S22**).

### On-chip sorting of the LinB library based on COU-3 activity

*E. coli* BL21(DE3) cells were transformed with the linB999 library and grown in LB with ampicillin overnight. Protein expression was induced with 0.5 mM IPTG and carried out for 18 h at 20 °C. Cells were harvested, washed twice with PB buffer (20 mM phosphate, pH 8.0), concentrated, and stored at −20 °C until use.

### DNA recovery

The PCR recovery of plasmid DNA from the sorted cells was performed using primers: F: 5′-GAAGGAGATATACATATGAGC-3′ and R: 5′-CGAGTGCGGCCGCAAGCTTTCAG-3′. Sorted droplets were collected directly into a PCR master mix (miliQ water, primers, Verifi buffer). Verifi polymerase was added after the initial 98 °C denaturation step. PCR was run for 35 cycles (Table SX), followed by gel electrophoresis and gel extraction. Reassembled variants were generated by Gibson Assembly of the purified PCR fragment into NdeI/HindIII-linearized and dephosphorylated backbone. Products were transformed into *E. coli* DH5α via heat-shock, recovered in SOC, and plated on LB agar with ampicillin.

### Large-scale overproduction of LinB hits and purification

Recombinant pET21b::linB variants were expressed in *E. coli* BL21(DE3). Two-litre cultures were grown in LB–ampicillin and induced with 50 µM IPTG at OD_600_ ≈ 0.8, followed by overnight expression at 20 °C. Cells were harvested, resuspended in phosphate-imidazole buffer (pH 7.5), lysed by sonication, and clarified by centrifugation. Proteins were purified by Ni²⁺-affinity chromatography and further polished by size-exclusion chromatography (Superdex 200) in PB buffer (pH 8.0). Protein concentration and purity were assessed by UV absorbance, and the final samples (∼5 mg/mL) were flash-frozen or lyophilised.

### Thermal stability

Thermal unfolding was studied using NanoDSF technology on Prometheus Panta (Nano-Temper, Germany) by monitoring tryptophan fluorescence over the temperature range of 20 to 95 °C, at a heating rate of 1 °C/min. The protein samples were dissolved in either 1mM HEPES buffer or 50mM PB buffer. The thermostability parameters (*T*_on_ and *T*_m_^app^) were evaluated by PR.Panta.Analysis software from the fluorescence ratio of 350/330 nm. The results of the thermal stability measurements are shown in **Table S14**.

### Steady-state kinetics

Kinetic measurements were performed in 20 mM potassium phosphate buffer (PB, pH 8.0). Enzyme solutions were filtered (0.2 μm), quantified by UV absorbance, and diluted to final concentrations of 125 nM (WT, I138N, and P137S) or 1.0 μM (V173F, P208, and R209). COU-3 was prepared as a 20 mM DMSO stock and diluted in PB buffer to final concentrations of 25–250 μM. Fluorescence kinetics were recorded using a CLARIOstar Plus plate reader (λ_ex_/λ_em_ = 402/474 nm) with dual injectors. Enzyme solutions (75 μL) were dispensed in triplicate into 96-well plates, followed by automated substrate injection (75 μL), brief mixing, and fluorescence monitoring at 25 °C.

### Substrate specificity profiles

Activity measurements for the determination of substrate specificity were conducted on the capillary-based droplet microfluidic platform MicroPEX^25^, enabling the characterisation of specific haloalkane dehalogenase activity within droplets for multiple enzyme variants in a single run. A set of 27 representative halogenated substrates was used to acquire the substrate specificity profile^42^. A detailed description of the microfluidic method can be found elsewhere^42,43^. The results of substrate specificity screening are provided in **Table S16**.

### HDX-MS

HDX-MS experiments were performed using a LEAP HDX robotic platform. Proteins were diluted in phosphate buffer (pH 8.0), adjusted to 2 μM during labelling, and incubated in D₂O buffer for time points ranging from 10 to 7200 s. Exchange was quenched at low pH, followed by on-line proteolysis and peptide separation at 2 °C prior to MS analysis. Fully deuterated controls were prepared for back-exchange correction. LC-MS measurements were carried out on an Agilent 1290 UHPLC coupled to a Bruker timsTOF mass spectrometer. HDX data were acquired in technical triplicates and processed using DeutEx and pyHDX to obtain residue-level deuterium uptake profiles.

### Flexpert predictions

To make the predicted values comparable among systems (and also to make them comparable to other flexibility measures), all values obtained by Flexpert v.1 (as downloaded from GitHub: https://github.com/KoubaPetr/Flexpert/releases/tag/v1) were internally standardised per system. Thus, each value from a LinB variant prediction series was transformed in the following fashion:

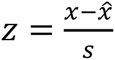

where *z* is the normalised per-residue value, *x* is the predicted value (Flexpert score), *x̂* is the average of predicted values, and *s* is their standard deviation. These standardised values were mapped on the LinB wild-type structure and represented both in colour (rainbow) and thickness (thicker representations indicate larger values). The comparison of the LinB wild-type flexibility values with those of the other variants was achieved by calculating, per residue, the log_2_ ratio of the variant flexibility over the wild-type one.

### Molecular Modelling

Molecular modelling was employed to probe substrate binding, tunnel accessibility, and catalytic reactivity in LinB variants. Docking was used to identify reactive binding poses of the COU-3 substrate, and classical molecular dynamics simulations were performed for free enzymes and enzyme–substrate complexes in explicit solvent. Near-attack conformations for the S_N_2 reaction were analysed, and QM/MM calculations were applied to estimate activation barriers. Tunnel properties and substrate translocation through major access pathways were evaluated using CAVER and adaptive steered molecular dynamics (ASMD).

Further details regarding the methods can be found in the Supplemental Information.

## Author’s contribution

H.F. performed library generation and cultivation, DNA rescue, PCR optimisation, production and purification of hits, protein characterisation, and data visualisation. D.K. assembled the FADS instrument, carried out microfluidic-level assay testing and FADS sorting, performed protein characterisation, substrate specificity analysis, kinetic measurements, and data analysis. A.J., T.B., F.W.Y.C., and T.K. contributed to FADS instrument assembly, microfluidic-level assay testing, and FADS sorting. M.S., M.M., and A.Ja. synthesised and characterised the coumarin substrate. M.Su. performed PCR optimisation and hit production/purification. M.Maj. contributed to library cultivation and PCR optimisation. M.V. performed protein characterisation, substrate specificity testing, data analysis, and data visualisation. J.P.-I. carried out flexibility prediction, data analysis, and visualisation. S.M.M. performed docking, molecular dynamics, and QM/MM simulations. V.N. and P.V. conducted HDX-MS measurements and data analysis; P.V. also contributed to protein characterisation. D.D. performed protein characterisation, substrate specificity testing, and data analysis. C.P.S.B. tested the assays. P.K. performed flexibility prediction. M.Ma. optimized gene amplification. S.M. performed flexibility prediction. H.F., D.K., M.V., J.P.-I., S.M.M., and Z.P. prepared the initial manuscript draft. A.J.dM., J.D., J.S., U.T.B., D.B., S.M., L.H., M.Ma., P.Kl., S.S., and Z.P. conceived the project and devised the research plan. Z.P. additionally performed kinetic measurements, protein characterisation, substrate specificity testing, and data analysis. All co-authors contributed to writing and revisions of the manuscript.

## Acknowledgements

The authors would like to acknowledge funding from the Czech Science Foundation (grant no. 25-15784L and GX25-17329X), by the European Union and Ministry of Education, Youth and Sports of the Czech Republic within ESIF-MEYS Johannes Amos Comenius Programme (OP JAK) under the project CLARA (CZ.02.01.01/00/23_029/0008437). The authors also acknowledge partial financial support from ETH Zurich and the Swiss National ScienceFoundation (Grant number: 205321/176011/1). This work was also supported by the European Union’s Horizon Europe Framework Programme under the grant agreement No. 101136607 (CLARA) and research and Innovation Programme under the grant agreement No. 857560 (CETOCOEN). The authors also thank the RECETOX Research Infrastructure (No. LM2023069), financed by the Ministry of Education, Youth and Sports, for its supportive background. Brno Ph.D. Talent Scholarship holder Hana Faldynova acknowledges funding from the Brno City Municipality.

