## Supplemental Information for "Uncovering Functional Distant Mutations by Ultra-High-Throughput Screening of Dehalogenases"

### List of Content

6.1 Thermal stability ... 51
6.2 Steady-state kinetics ... 52
6.2.1 Analysis and statistics of the steady-state kinetic data ... 53
6.2.2 Steady-state kinetic results ... 54
6.3 Substrate specificity profiles ... 56
6.4 HDX-MS ... 57
7 In Silico Hits Characterization ... 60
7.1 Structure modelling of the proteins ... 60
7.2 Flexibility Prediction ... 60
7.2.1 Flexibility mapping ... 60
7.3 Molecular modelling ... 61
7.3.1 System preparation for docking and simulations ... 61
7.3.2 Molecular docking ... 61
*7.3.3* Classical molecular dynamics ... 63
7.4 Access tunnel calculations ... 64
7.4.1 QM/MM adiabatic mapping ... 68
7.4.2 Adaptive steered molecular dynamics ... 69
8 References ... 71

Figure S18: Schematic reaction mechanism of the hydrolytic SN2 dehalogenation catalysed by a dehalogenase. The enzyme (E) first binds the halogenated substrate to form the enzyme–substrate complex (ES). In the initial $S_N2$ step, the catalytic nucleophile (typically an Asp residue) attacks the electrophilic carbon bearing the halogen, displacing the halide and generating a covalent ester intermediate (EI). A water molecule is subsequently activated by a catalytic base within the active site, enabling nucleophilic attack on the acyl-enzyme intermediate. This hydrolysis step releases the dehalogenated product and regenerates the free enzyme (EP → E), completing the catalytic cycle. Curved arrows represent electron flow during nucleophilic substitution and subsequent hydrolytic cleavage .......... 38

Figure S30: HDX-MS results. (A) Average deuterium uptake of LinB WT (grey), LinB P208S (yellow), and LinB I138N (blue) after 10 s of $D_2O$ labelling. Mutation sites are indicated by dashed lines on the secondary structure. (B) Heatmap representation of D-uptake, with black indicating the lowest and yellow the highest uptake. (C-E) D-uptake values mapped onto the LinB WT structure (PDB: 1MJ5). LinB WT (C) and LinB P208S (D) display highly similar uptake profiles across the structure, whereas LinB I138N (E) shows markedly higher D-uptake in the cap-domain (residues 130-230), indicating increased flexibility due to the mutation. .. 59

Figure S31: Docking of COU3, the three LinB variants. The reactive binding conformations of COU3 docked to WT (magenta), I138N (yellow) and P208S (yellow) are superimposed almost perfectly. Only the structure of WT is represented as white cartoons, and the catalytic residues as sticks; the distance between the reacting carbon in the substrate and carboxylic oxygen is represented by the red dashed line, and the hydrogen bonds between the chlorine and the halide-stabilising residues N38 and W109 are represented by the black dotted lines........................ 62

Figure S32: Structure of the NAC complexes in the three variants. A) Representative NAC complex of COU3 (in orange) in the active site of LinB WT (in grey), displaying the catalytic and the mutated residues, superimposed with the NAC complexes of mutants I138N and P208S. For simplicity, only the mutated residues are represented for the mutant variants (residues N138, in blue, and S208 in yellow); the distance between the reacting carbon in the substrate and the carboxylic oxygen ($d_{C\text{-}O}$) is represented by the red dotted line, and the hydrogen bonds between the chlorine and the halide-stabilizing residues N38 and W109 are represented by the black dotted lines; the distance between the hydroxyl oxygen atom of S208 and the second N38 hydrogen atom ($d_{O\cdots H}$) is represented by the yellow dotted line. B) Histogram distribution of the $d_{O\cdots H}$ distance illustrated in (A) over all the NAC complexes detected in the MD simulations with the P208S variant. The generally small values of the $d_{O\cdots H}$ distance ($d_{O\cdots H} \leq 3.5$ Å in 74% of the NACs) suggests a strong interaction between the serine hydroxyl oxygen with one of the N38 hydrogen atoms. This may lead to a decrease in the stabilization provided by the other N38 hydrogen atom to the transition state, thus explaining the higher activation barrier calculated for of the $S_N2$ reaction in P208S. ................................ 65

Figure S33: The three main tunnels found in the MD simulations with the free enzymes (LinB WT is shown here). A) “timeless” tunnels, and B) tunnels in the snapshot at 410 ns. The timeless tunnels show the centreline of all the tunnels detected in all the snapshots of the MD, superimposed and coloured by the respective cluster colour (p1 blue, p2 green, and p3 red). The tunnels are named according to Chovancová et al.[37,39] ............................................................ 66

Table S6: Calculated steady-state parameters of DmmA with COU-1, COU-2 and COU-3 ($k_{cat}$ = k+2, Km = k-1/k+1, Ki = k+3/k-3, KSI = k-4/k+4). Standard errors are calculated by dividing standard deviations (sigma – sum of the residuals squared divided by the number of measurements minus one – noise quantification) by the square root of the number of measurements (which means that the estimated accuracy improves with the square root of the number of measurements, which may not always be accurate). $\chi$2 is defined as the sum of squared residuals, normalised by the known sigma values. The $\chi$2 threshold for calculation of lower and upper limits was set to 0.95. ........................................................................ 37

Table S17: Docking binding affinities ($\Delta G_{bind}$, in kcal/mol) of the productive binding conformations. 63
Table S18: Properties of the main tunnels during the MD simulations with the free LinB variants. * 67
Table S19: PMF values at the final distance of 4 Å, obtained by ASMD for tunnels p1 and p2, and respective statistical significance. * 70

### 1 Fluorescence-activated droplet sorting system – Cinderella

An assembled fluorescence-activated droplet sorting (FADS) system, optimised for high-throughput screening of enzymatic activity, is presented and hereafter referred to as Cinderella. The system integrates an inverted fluorescence microscope with customised microfluidic components, enabling precise droplet generation, single-cell encapsulation, and fluorescence-based sorting. Through extensive optimisation of optical, microfluidic, and computational parameters, we achieved reliable detection and sorting of droplets based on enzymatic activity. Benchmark experiments with both fluorescent dyes and enzymatic variants demonstrated the system's ability to discriminate between different levels of activity with high precision.

Additionally, we developed a custom analysis software (Cinderella$^{A}$) that enhances data processing capabilities, facilitating on-time threshold determination and post-experiment analysis. This FADS platform represents a significant advancement for applications in enzyme engineering, directed evolution, and single-cell analysis.

#### 1.1 System Overview and Components

The fluorescence-activated droplet sorting system represents a sophisticated integration of optical instrumentation with microfluidic technology, creating a powerful platform for high-throughput screening applications. At its core, the FADS system is built around a standard inverted fluorescence microscope enhanced with specialised components that enable precise droplet manipulation and analysis (*Figure S1*). The microfluidic system employs high-precision pumps, either syringe-based or pressure-driven, providing the fine control necessary for consistent droplet generation and manipulation. Fluorescence emission signals are captured using a photomultiplier tube (PMT) and processed in real-time through a Field Programmable Gate Array (FPGA) card, which generates sorting pulses when signals meet predefined threshold parameters. These sorting pulses are subsequently amplified using a high-voltage amplifier, deflecting droplets containing cells or molecules of interest into designated collection channels through dielectrophoretic forces.

Our implementation utilises an IX73 double-deck inverted microscope (Olympus, Japan) with a CELESTA Light Engine (Lumencor, USA) as the primary optical platform. This foundation is complemented by low-pressure syringe pumps (neMesys, Cetoni, Germany) for droplet generation on the microfluidic chip and a four-channel pressure-driven system (OB1 MkII,

Elveflow, France) for droplet reinjection and sorting. The entire assembly is mounted on a Nexus optical table with active isolator legs (T1220CK, Thorlabs).

A violet laser with a wavelength of 405 nm (CELESTA Light Engine, Lumencor) serves as the excitation source for the described sorting assay (see section 2). Unless otherwise specified, all subsequent components are supplied by Thorlabs.

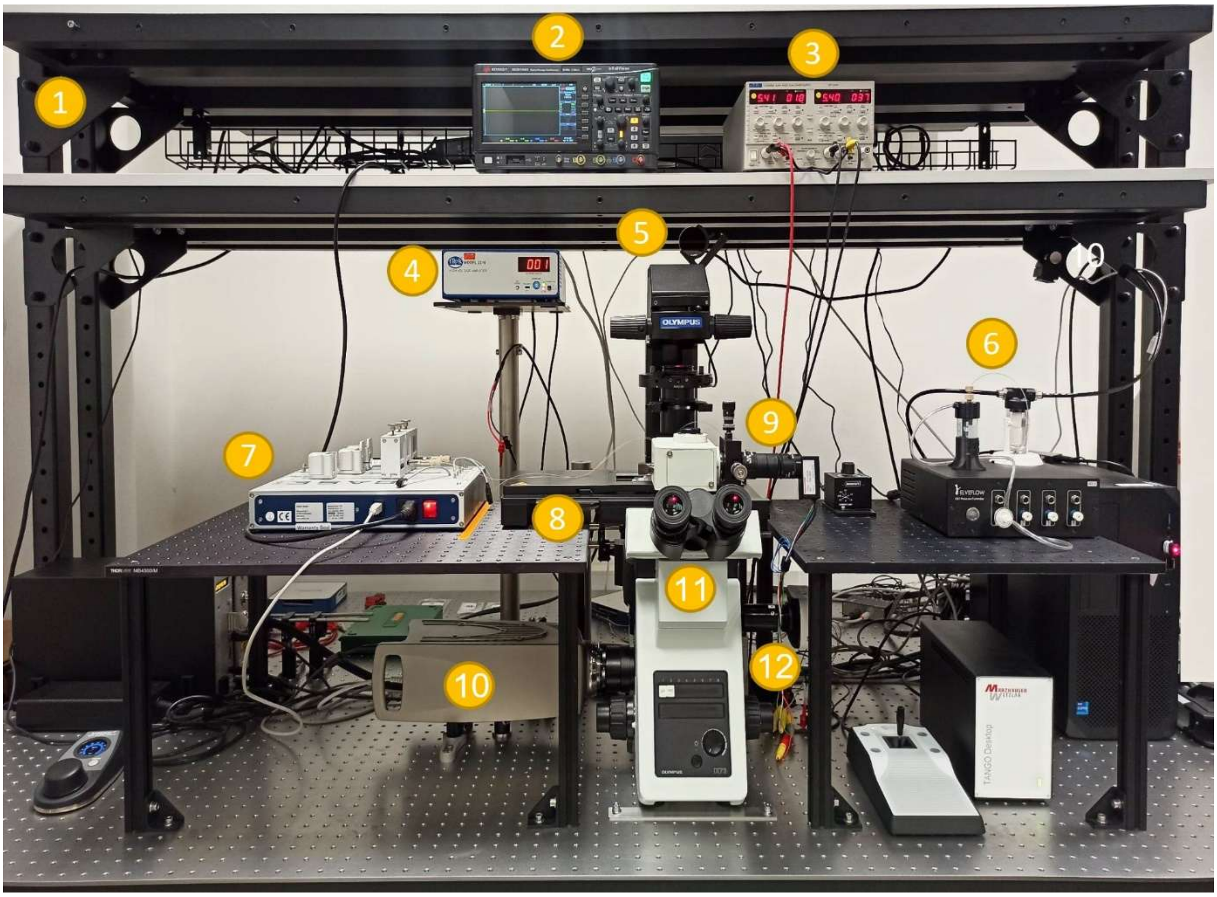

*Figure S1: An assembled setup used for droplet sorting - front view; free-standing optical table shelf (1), oscilloscope DSOX1204G (2), power supplier PL303QMD (3), high-voltage amplifier Trek 2210 (4), CollLED pE-300white illuminator (5), pressure pumps OB1 (6), syringe pumps neMesys (7), motorised stage SCAN IM 120 x 80 (8), PMT detector H10720-20 (9), high-speed camera iSpeed (10), inverted microscope IXT3 (11), fluorescence camera Infinity 2 (12).*

The laser beam is first directed through a plan achromatic 10× objective (PLN10X/0.25, Olympus, Japan), which is mounted in an adapter with external SM1 threads and internal RMS threads (SM1A3). This assembly is attached to a 30 mm cage XY translation stage (CXY1A) to enable precise alignment of the objective in the optical path. After the objective, a graduated ring-actuated iris diaphragm (SM1D12C, mounted in a CP33T/M holder) is positioned to shape the beam profile. The iris is adjusted to block the beam's peripheral (corona) regions, allowing only the central, most uniform portion (the core) to pass through. This configuration ensures a well-defined, homogeneous beam for further focusing and alignment.

The refined beam is then reflected by a visible-light mirror (BB1-E02) mounted in a kinematic mirror mount (KM100) on a pedestal post assembly with a right-angle clamp (RA90/M), directing it into a telescope assembly for precise focusing onto the specimen. The telescope consists of a second visible-light mirror (BB1-E02) in a kinematic cage mount (KCB1/M), which reflects the beam onto the curved surface of a plano-convex NBK-7 lens (LA1805-A, f = 30 mm, mounted in CP33T/M). The beam then passes through a second plano-convex lens (LA1805-A), oriented in the opposite direction and mounted in a Z-axis translation mount (SM1ZA), exiting again through the curved surface. This lens pair functions as a beam expander and focus adjuster, enabling precise control of beam diameter and convergence. Next, the beam passes through a longer focal length plano-convex lens (LJ1558RM, f = 300 mm) placed in a rotation mount (CRM1T/M), which focuses the beam onto the back focal plane of the microscope objective. More specifically, inside the microscope, the beam is reflected by a right-angle dichroic mirror (DMLP425R) and directed through a 20× microscope objective (LUCPLFLN20X/0.45, Olympus, Japan), which focuses it into the microfluidic channel. This arrangement ensures efficient excitation of the chip and supports sensitive fluorescence detection for droplet sorting.

Fluorescence emission from droplets is collected by the objective, transmitted through the dichroic mirror, and guided by the microscope body to the camera port, where the PMT detector is coupled. The overall detector assembly consists of camera adapters (U-TV1X-2 and U-CMAD3, both Olympus, Japan), an adapter with external SM1 threads and internal C-Mount threads (SM1A10), a 40mm pinhole (P40K) mounted on a XY translator (ST1XY-D/M), followed by a lens tube filter holder (SM1QB with SM1QT) equipped with an MF-469-35 GFP bandpass filter. A focal length of 50 mm (LA1131-A) focuses the light onto the surface of the photomultiplier tube (PMT, H10723-20; Hamamatsu) through a C-mount ring (A9865, Hamamatsu, Japan).

The PMT output signal is routed to a shielded I/O connector terminal block (SCB-68A, NI, USA), which is connected to a field-programmable gate array (FPGA) card (PCIe-7842R, NI, USA) that executes a program written in the LabVIEW FPGA module (NI, USA). The program was adapted from Abate's lab [https://github.com/AbateLab/sorter-code] and modified for our purposes and is hereafter referred to as Cinderella[C]. The program performs real-time droplet identification based on peak height and width, applies user-defined intensity thresholds, and triggers dielectrophoretic (DEP) sorting pulses. The DEP pulses are amplified 100-fold using a high-voltage amplifier (Model 2210, Trek, USA) to drive the sorting electrodes.

##### 1.2 Fabrication of microfluidic devices

Microfluidic designs for droplet generation and sorting chips were created using AutoCAD (Autodesk, San Francisco, USA), and photolithography film masks printed by Micro Lithography Services (Chelmsford, UK). Briefly, a layer of GM 1070 SU-8 photoresist (Gersteltec Engineering Solutions, Puly, Switzerland) was spin-coated at 3500 rpm onto a 100 mm diameter silicon wafer (yielding a final resist thickness of 20 µm) and soft-baked at 95 °C for 20 minutes. Microfluidic designs were patterned onto the photoresist layer by UV exposure (500 mJ/cm2) through a photolithographic mask. This was followed by baking at 95 °C for 10 minutes, then development using 1-methoxy-2-propanol acetate (Acros Organics, Geel, Belgium) for 150 seconds. The generated master moulds were exposed to chlorotrimethylsilane (ABCR, Karlsruhe, Germany) vapour in a desiccator at 150 mbar pressure to facilitate subsequent PDMS demoulding.

##### 1.3 Microfluidic Chip Fabrication

We fabricated two distinct chip designs: a co-encapsulation chip for droplet generation and a sorting chip for fluorescence-based droplet separation. Both chips were manufactured using Polydimethylsiloxane (PDMS) elastomer (Sylgard 184, Dow, USA). This material choice aligns with established protocols in the field and ensures compatibility with biological samples and fluorescence detection methods.

The fabrication process begins with the careful preparation of the PDMS mixture. A silicone elastomer base is combined with a crosslinker at a precise 10:1 weight ratio, then thoroughly mixed for 3 minutes to ensure homogeneity. The resulting mixture must be completely degassed in a vacuum desiccator to eliminate any bubbles that could potentially compromise the integrity of the microfluidic channels. Once degassed, the PDMS mixture is poured onto a wafer attached to a plastic or aluminium dish, creating the foundation for the microfluidic channels. The assembly is then baked at 65 °C for a minimum of 3 hours in a universal oven (UF30, Memmert, Germany), fully curing the PDMS. Subsequently, the PDMS slab was peeled off the mould and fluid inlets, and outlets were created using Ø1 mm UniCore Punches (QIAGEN, Germany).

The plasma treatments were conducted using a capacitively coupled discharge system (Zepto ONE, Diener Electronics) with a 1.7 L gas chamber equipped with a glass shelf. The system is equipped with a 13.56-MHz RF generator with variable power (0–30 W). The cut PDMS slabs and glass microscope slides were placed into the plasma chamber, which was then pumped down for 3 minutes. The pressure of oxygen in the sealed gas-flushed chamber was adjusted to 2 Nl·h$^{-1}$ and held for 3 minutes, to allow it to stabilise, at which point the RF generator was activated for 45 seconds. Following chamber ventilation, the activated surfaces of PDMS slabs and glass slides were immediately brought into contact to achieve bonding. The assembled chips were then placed on a hot plate and cured at 120 °C for at least 20 min to strengthen the bonding. Surface modification represents an essential step in preparing the chips for droplet-based applications. Despite PDMS being naturally hydrophobic, the plasma treatment used for bonding reduces this hydrophobicity, necessitating additional surface modification. For single cell encapsulation devices, we perform hydrophobic modification by filling the channels with 1% TCPS in HFE-7500 oil, incubating for 5 minutes at room temperature, and thoroughly washing with pure HFE-7500 oil. This hydrophobic coating ensures the stable formation of water-in-oil droplets and prevents unwanted wetting of the channel walls by the aqueous phase.

##### 1.4 Encapsulation Chip Design and Function

The encapsulation chip is designed according to a flow-focusing microdroplet generator, featuring three inlets and one outlet (*Figure S2*). This configuration enables the precise formation of monodisperse droplets through the controlled interaction of immiscible phases. One inlet accommodates the oil phase, specifically a solution of 1.25% (w/w) 008-FluoroSurfactant RAN in HFE-7500 oil. The surfactant plays a crucial role in reducing surface tension and preventing droplet coalescence, while the HFE-7500 oil provides a biocompatible continuous phase for the water-in-oil emulsion. The second inlet is dedicated to the Coumarin substrate (COU-3). The third inlet is dedicated to the sample, which contains either purified enzyme or a bacterial suspension in PB buffer with 25% OptiPrep (Merck, Germany), depending on the specific experimental requirements.

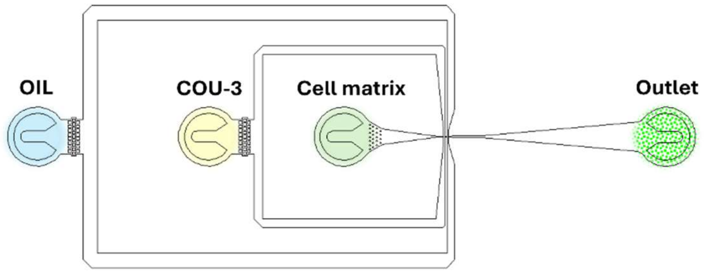

*Figure S2: Schematic of the encapsulation chip with three inlets. The aqueous phase (COU-3 and cell matrix) flow*

*rates were set equal to adjust the cell density (OD600) for optimal Poisson distribution and to control the substrate concentration.*

#### 1.5 Sorting Chip Design and Electrode Fabrication

The sorting chip incorporates specialised features for dielectrophoretic droplet sorting. A distinctive aspect of this chip is the incorporation of electrodes fabricated from Field's metal, an eutectic alloy comprising 32.5% bismuth, 51% indium, and 16.5% tin. This material offers several advantages, including electrical conductivity, lower toxicity compared to alternatives such as Wood’s metal, and a low melting point of approximately 62 °C, which facilitates easier processing.

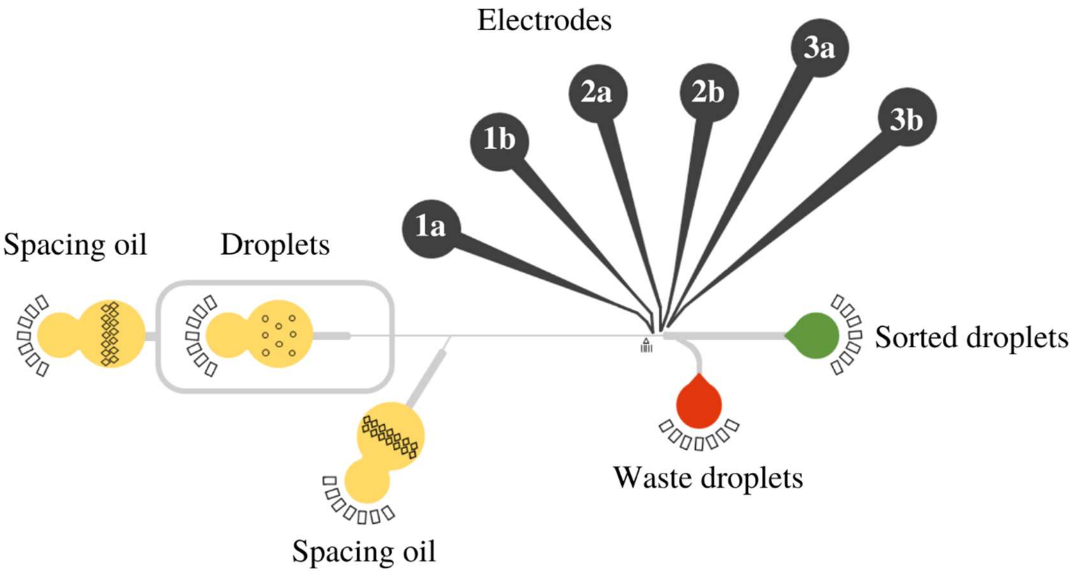

*Figure S3: Schematic of sorting chip. The reinjected droplets are separated by a pair of spacing oils. Each electrode channel has one inlet and one outlet, where the pieces of Field's metal or the metal pin are inserted during fabrication. On the side of the electrodes, there is a positive outlet channel through which the active hits are deflected. The droplets below the fluorescence threshold spontaneously flow to the negative (waste) channel.*

The electrode fabrication process involves melting Field's metal granules into an aluminium dish on a hot plate set at 100 °C. The molten metal is drawn into a Teflon tube (Ø 1 mm) with an oblique cut at one end, allowed to cool, and then carefully removed from the tubing. The resulting metal wires were cut into appropriate lengths for insertion into the PDMS sorting chip. Pieces of Field’s metal wires were inserted into inlets marked with 1a, 2a, and 3a in *Figure S3*, while metal pins with a gold layer were inserted into outlets marked with 1b, 2b, and 3b, respectively. The entire assembly was heated on a hot plate at 100 °C until the electrode channels were filled with metal, and another set of metal pins was inserted into the inlets “a”, then allowed to cool to room temperature. A microscope and a multimeter were used to verify proper channel filling and to check electrical insulation between neighbouring electrodes by measuring resistance.

#### 1.6 Handling Droplets

Hydrophobised vials were used for the collection, incubation and reinjection of the droplets throughout the whole process. The inner surface of conical vials was hydrophobised according to the silanisation protocol published in Angewandte Chemie[1]. Briefly, the vials were plasma treated using same conditions as for the chip bonding. The conical vials (MACHEREY-NAGEL, 702860) in the Junior UniRack (Simport, S500-25) were placed into the plasma chamber, and the chamber was pumped down for 3 minutes. The gas pressure in the sealed, gas-flushed chamber was reduced to 2 $Nl \cdot h^{-1}$ of oxygen flow and maintained for 3 minutes, allowing the inner atmosphere to stabilise. The RF generator was then turned on after this time. A 1% solution of trichloro(1H,1H,2H,2H-perfluorooctyl)silane (Sigma-Aldrich, 448931) in toluene was prepared during the etching period. After 45 seconds of plasma treatment, the chamber was ventilated, and the vials were immediately filled with the freshly prepared silanisation solution, capped with a silicone/PTFE seal (Machery-Nagel, 70245), and stored in a hood for 3 hours. After incubation, the silanisation solution was poured from the vials, washed twice with toluene, then filled with acetone and sonicated for 20 minutes in a Sonorex Digitec ultrasonic bath (Bandelin, Germany). The process was finished by washing the vials using absolute ethanol and drying under compressed air. The vials were closed with new caps and stored for later usage. The hydrophobic coating is stable for at least 6 months.

A droplet reservoir was required for pressure-driven droplet reinjection into sorting chips. Unfortunately, there are no commercially available adapters for such small vials; therefore, the insert vial holder for Falcon tube was designed using Autodesk Inventor. Bottom tube holder was purchased from Essi Biotech (Slovenia), and the M-size cap adapter was purchased from Elveflow (France). An alternative Falcon tube holder was downloaded from (https://www.printables.com/model/79347-flared-falcon-tube-holder-hex-holes). The insert and tube holder were printed on a Prusa MK4 3D printer (Czech Republic). The assembled reservoir, equipped with Tygon tubing for the compressed air inlet and PTFE tubing for the droplet outlet, is depicted in *Figure S4*.

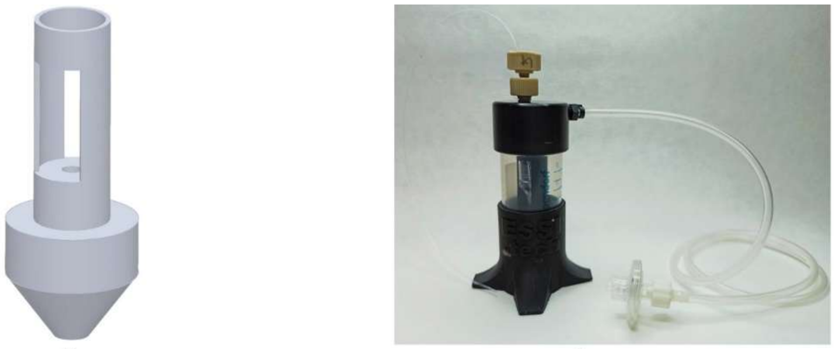

*Figure S4: In-house-made vial holder insert for Falcon tube-based pressure chamber for reinjection of the droplets into sorting chips. A 3D model of the insert (a) and assembled pressure chamber used in the laboratory (b).*

#### 1.7 System Optimisation and Operation

The successful implementation of Cinderella required careful optimisation across multiple parameters to ensure reliable performance. As a newly assembled system, initial optimisation focused on aligning the laser into the optical path and achieving precise beam focusing into the microfluidic channel. This critical adjustment directly affects the sensitivity and accuracy of fluorescence detection, which underpin subsequent sorting decisions.

A general alignment of the laser beam was performed by precisely positioning each optical element along the beam path, from the laser source to the microscope specimen stage. During this procedure, the laser output was continuously monitored to ensure stable and reproducible illumination. Measurements of the beam intensity were obtained using an optical power meter equipped with a compact photodiode sensor (PM100D with S121C detector, Thorlabs). This allowed fine adjustments of the optical components to achieve optimal beam focusing and uniform power delivery at the sample plane.

Final laser alignment was performed using a fluorescence camera, a photomultiplier tube (PMT), and the sorting software Cinderella[C] to determine the optical configuration that yielded the highest, most stable signal during sorting. Initial focusing was performed at the inlet of the encapsulation chip, where fluorescence intensity is maximal and continuous because droplet formation has not yet begun. Aligning the beam at this point allowed for more accurate control of the laser profile and its spatial distribution. Once the optimal signal was identified, the focus position was shifted downstream to track droplet trajectories after the T-junction, with minor adjustments made as needed to maintain signal quality.

Optimal parameters for droplet generation were determined to ensure consistent droplet formation with suitable dimensions for single-cell trapping. Systematic optimisation identified flow rates of 335 $\mu L \cdot h^{-1}$ for the oil phase and 30 + 30 $\mu L \cdot h^{-1}$ for the two aqueous phases as the most effective conditions. Under these settings, droplets exhibited uniform size distribution with minimal variability, a key factor for reproducible downstream analyses and sorting. The encapsulation workflow employed hydrophobically treated microfluidic chips, glass syringes and syringe pumps for accurate flow control, as well as hydrophobically modified conical glass vials for efficient droplet collection.

##### 1.8 Data Acquisition and Analysis Software

Recognising the limitations of the existing[2] "sort-code", we customised the software for our FADS system to improve data processing capabilities. The software development addressed specific needs in data acquisition, storage, and analysis that were not adequately met by the original sorting software available in the LabVIEW environment. While the original software, developed at the University of California, and available on GitHub, provided basic functionality, it lacked comprehensive analysis, data processing, and visualisation capabilities necessary for our application.

A key improvement involved changing the data saving format from the original *.acq format to the binary *.tdms format, which is native to LabVIEW. This change enabled the inclusion of essential metadata alongside measurement data, facilitating more comprehensive analysis. Each TDMS file was configured to contain 2,000 data cycles, along with metadata such as burst parameters, threshold values, date, pressure, and flowrate settings, laser properties, PMT voltage settings, pulse characteristics, and reagent information. This structured approach to data storage ensures that all relevant experimental parameters are preserved for subsequent analysis and reporting. During data acquisition, the software generates a series of TDMS files, each sequentially numbered with an incremental numerical suffix. This naming convention facilitates organised storage and retrieval of time-series measurement data in a hierarchical format. Such incremental indexing ensures chronological ordering, supporting efficient post-processing.

*Table S1: List of metadata stored in TDMS file from the experiment*

| **Parameter** | **Unit** |
|---|---|
| **Burst cycles** | - |
| **Cut-off value** | % |
| **Date** | - |
| **Droplet reinjection pressure** | mbar |
| **Laser intensity** | % |
| **Laser wavelength** | nm |
| **Name** | - |
| **Oil / Aqueous phase flowrate** | $\mu L \cdot h^{-1}$ |
| **PMT1 Gain Control Voltage** | V |
| **PMT2 Gain Control Voltage** | V |
| **PMT1 Voltage** | V |
| **PMT2 Voltage** | V |
| **Pulse amplitude** | V |
| **Pulse delay** | µs |
| **Pulse frequency** | Hz |
| **Reagents** | - |
| **Sampling frequency** | Hz |
| **Sorting pulses** | V |
| **Sorting threshold** | V |

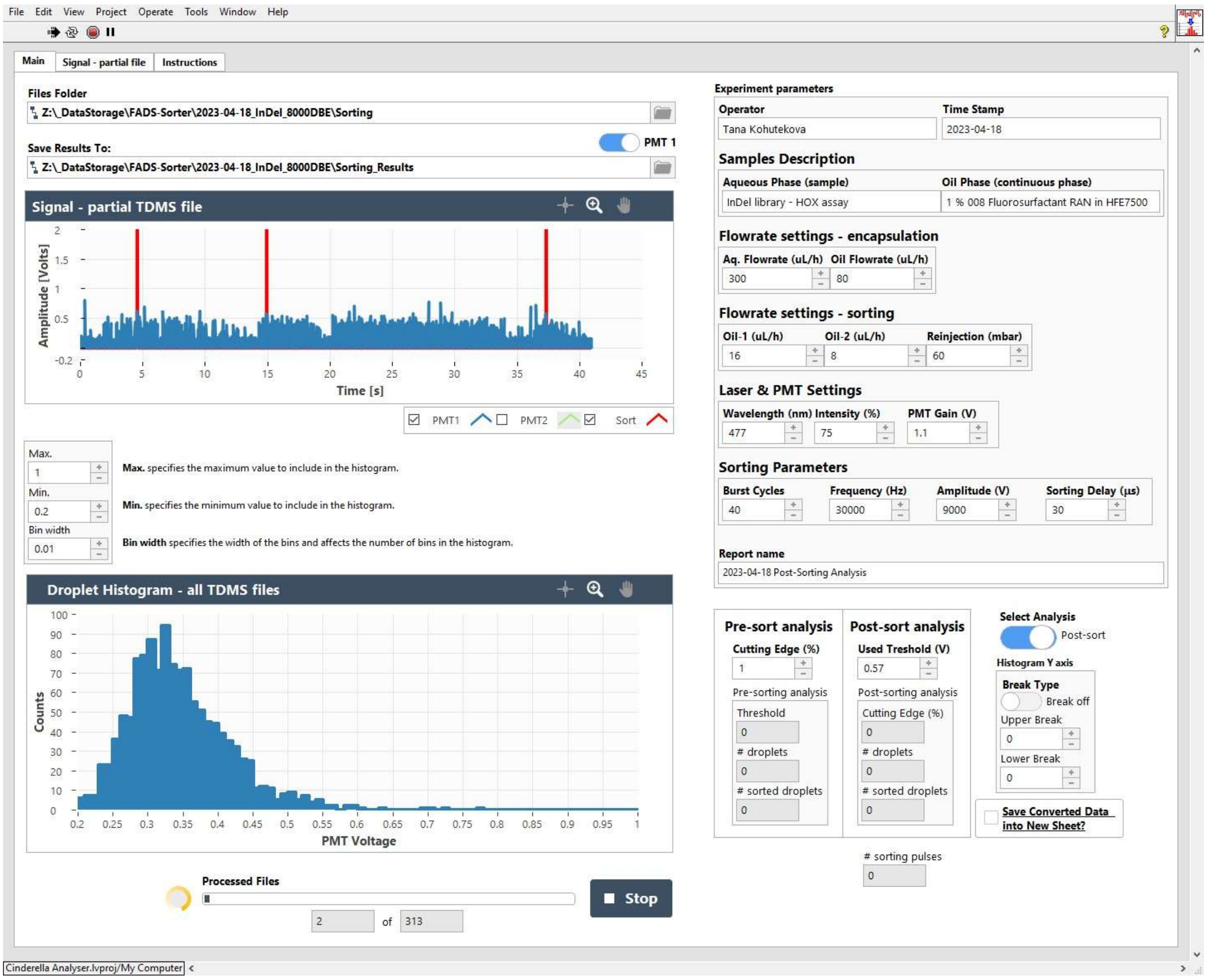

*Figure S5: Graphical user interface of Cinderella$^A$ analysis software for FADS data analysis programmed in LabView and integrated with Python.*

The graphical user interface (GUI) of our Cinderella$^A$ analysis software, implemented in LabVIEW bundled with Python, provides an intuitive interface for data analysis. Upon loading TDMS files from a selected directory, the software displays a graph with time on the x-axis and measured voltage from the chosen PMT channel (blue) and sorted pulses (red) on the y-axis. Users can adjust histogram parameters, such as bin width and voltage range, and the resulting histogram displays voltage on the x-axis and counts in the corresponding bins on the y-axis. Given the time-intensive nature of processing large datasets—with a single experiment potentially generating over 100 files—a progress bar visually and numerically indicates the processing status.

The analysis workflow (*Figure S6*) follows a structured process, beginning with the user selection of input parameters, including the folder containing the experimental data, the destination folder for results, and histogram parameters (minimum and maximum analysis voltage and bin width). The software loads all TDMS files from the specified folder and recalculates the raw signal values to voltage using constants specific to the FPGA card. The resulting PMT voltage values are stored in an array and processed by the CountHistogram function in Python, which creates a NumPy array and integrates the signal using a FilterPeakDetection function with a window length of 150 samples. This processing smooths the raw signal, suppresses noise and random high peaks, and highlights peaks of interest for easier detection.

In the developed software, the user begins by selecting a directory containing the TDMS files to be analysed and specifying the primary PMT (photomultiplier tube) channel of interest. The software then processes the selected files and automatically generates multiple output formats, including a histogram in .xls format, a histogram image in .png format, and a detailed report in .pdf format. All resulting data are stored in a dedicated output directory chosen by the user.

For each loaded TDMS file, the software displays a corresponding graph in the main window. The time course is plotted along the x-axis, while the measured voltage signals from the selected PMT channel and the sorting pulses are displayed on the y-axis, in blue and red, respectively. Users can interactively adjust the histogram parameters—specifically, the bin width and voltage range—used by the underlying Python function for data computation. As the PMT operates within a sensitivity range up to 4 V, this voltage defines the default upper boundary for analysis. After each iteration, the histogram for the currently processed TDMS file is rendered, with the x-axis representing voltage and the y-axis indicating the event count per bin.

Given that a single experiment typically consists of over 100 TDMS files, real-time processing feedback was implemented. A progress bar, accompanied by a moving progress ring, provides both visual and numerical indicators of the number of files already processed.

The right panel of the graphical user interface (GUI) is divided into two main sections. The upper section displays metadata retrieved from the TDMS files where available. For legacy datasets lacking embedded metadata, the user is prompted to manually enter the corresponding information. The lower section contains controls for selecting the analysis mode. Specifically, users can choose between two processing pathways: (i) a preliminary analysis performed prior to cell sorting to determine the optimal sorting threshold according to defined criteria, or (ii) a comprehensive post-sort analysis encompassing the entire experiment.

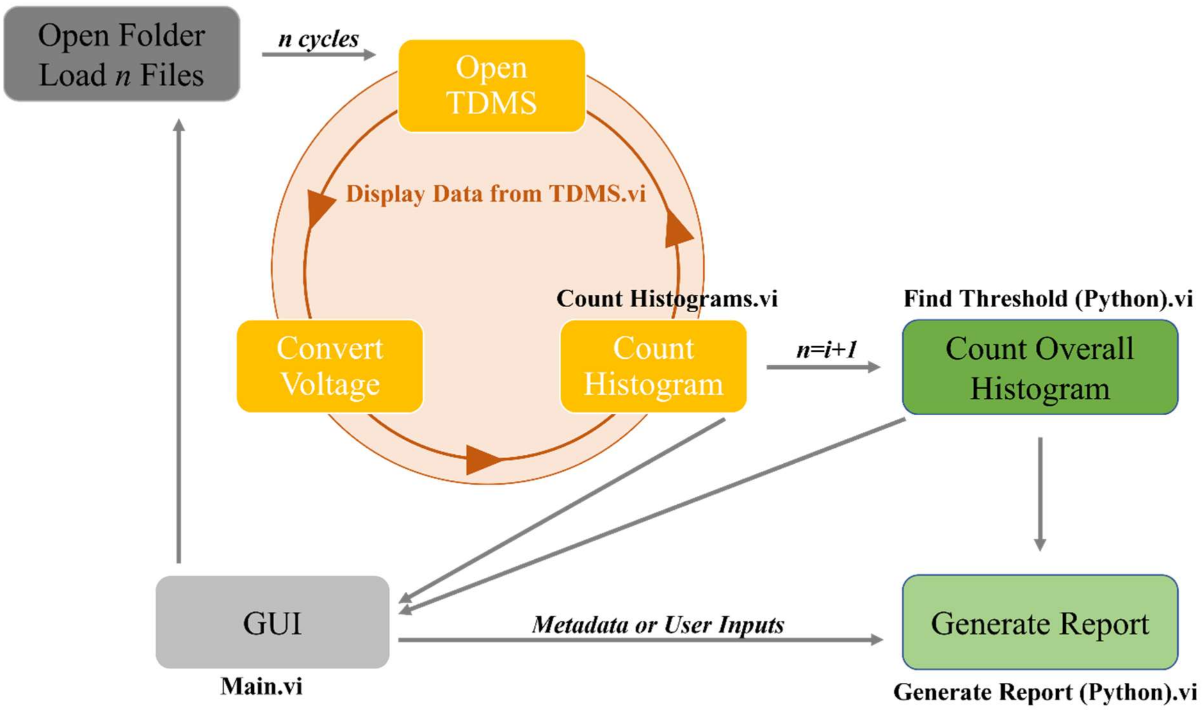

*Figure S6: A flowchart of the processes under Cinderella[A] software.*

In large-scale sorting experiments—often exceeding tens or even hundreds of thousands of droplets—typically only the most active 1 % or fewer are of primary interest. To facilitate more informative visualisation under these conditions, an option enabling histograms with broken axes was implemented. This feature allows users to define axis limits, thereby emphasising relevant data regions while maintaining overall data integrity. In the event of erroneous input or unexpected runtime issues, the software includes an emergency stop function. Activating the STOP button halts all ongoing operations and prevents any files from being written to disk.

There are two selectable analyses in the SW.

##### 1.8.1 Pre-sort analysis

The first analysis is used to estimate the threshold. During FADS, we expect the fluorescence signals from individual protein variants to follow approximately a normal (Gaussian) distribution. In such a distribution, most variants cluster around the average fluorescence intensity, while fewer variants show much higher or lower values. The goal is to capture only a small fraction of the top-performing variants—usually 1% or less of the total population. To do this, the Cinderella[C] collects data of 10-50 thousand droplets, and the analysis software examines the fluorescence histogram and fits it with a Gaussian curve. From this model, it determines the photomultiplier tube (PMT) intensity that marks the upper percentile corresponding to the desired Cutting Edge, such as the top 1 %. Variants with fluorescence above this threshold fall within the extreme high-fluorescence region of the distribution and are selectively collected for further characterisation.

##### 1.8.2 Post-sort analysis

This analytical module processes all fluorescence and sorting data acquired during the experiment, constructs a histogram of fluorescence intensities using predefined parameters, and determines the number of sorted droplets based on the selected intensity threshold. Furthermore, it enables quantitative comparison between the calculated droplet counts, the corresponding dielectrophoretic (DEP) pulse events recorded during sorting, and the droplet statistics reported by the Cinderella$^C$ software.

Future improvements to the software architecture are currently under consideration. In particular, the histogram generation process could be optimised by generating both standard and broken-axis histograms from aggregated data after all files have been processed, thereby improving computational efficiency and overall performance.

#### 1.9 Experimental Validation

To validate the functionality and performance of Cinderella, we conducted a benchmark experiment using a fluorescent dye at different concentrations. This experiment was designed to assess the system's ability to distinguish between different fluorescence intensities and to accurately sort droplets based on predefined threshold criteria.

The initial benchmark experiment utilised the HPTS (8-Hydroxypyrene-1,3,6-trisulfonic acid trisodium salt, Merck) fluorescent dye at three different concentrations: 10 μM, 40 μM, and 70 μM in 20 mM phosphate buffer with pH 8.2. HPTS was selected as a fluorescent pH indicator with excitation and emission properties (excitation peak at 460 nm, emission maximum at 510-515 nm). Each concentration was encapsulated separately using the optimised flow rates (300 $\mu L \cdot h^{-1}$ for the oil phase, 80 $\mu L \cdot h^{-1}$ for the aqueous phase), generating droplets that were collected for 1 hour each to ensure equal representation of all fluorescence intensities.

The generated droplets were reinjected into the sorting chip at 65 mbar, and the flow and frequency of the droplets were adjusted using two spacing oils at flow rates of 10 and 15 $\mu L \cdot min^{-1}$, respectively.

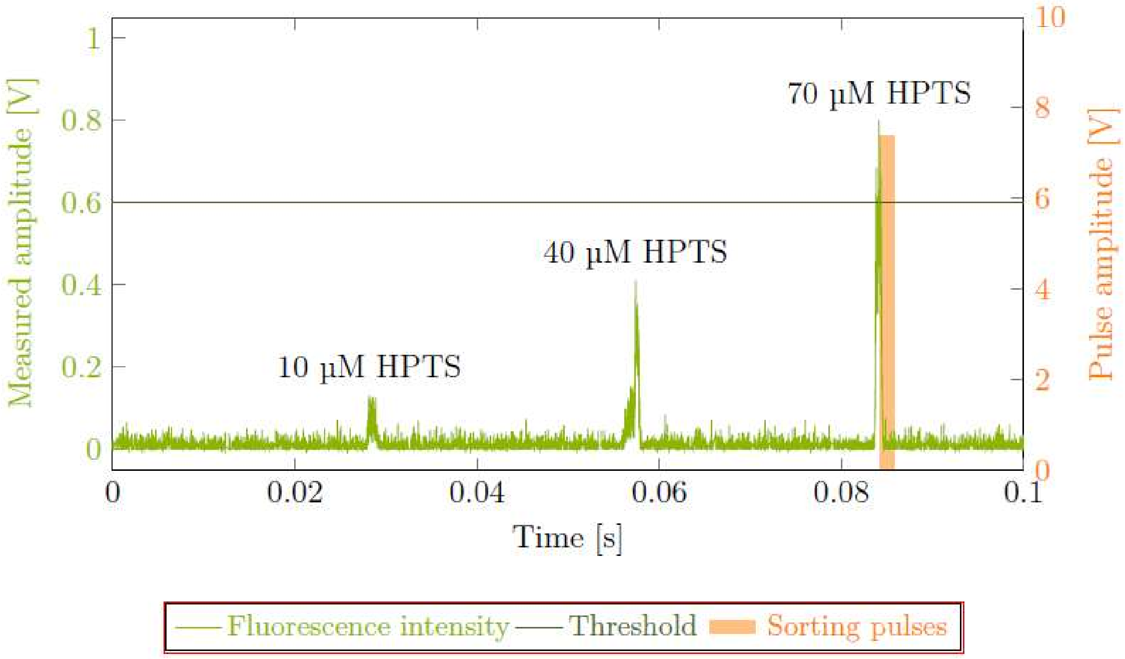

*Figure S7: A 10-ms segment of the signal obtained from a benchmark experiment with different concentrations of HPTS. A higher concentration produces a higher signal amplitude measured by the PMT in volts. The threshold level is visualised. The signal over the threshold triggers the generation of the sorting pulses and deflection of the droplet to the positive collection channel on the chip.*

Analysis of the resulting signal clearly demonstrated Cinderella's ability to distinguish peaks corresponding to different HPTS concentrations (*Figure S7*), with higher concentrations producing higher amplitude signals. This relationship between concentration and signal intensity confirms the quantitative detection capabilities of our system, which is essential for discriminating between enzymatic variants with different activity levels.

### 2 Fluorogenic Substrate

Two fluorogenic substrates for haloalkane dehalogenases (HLDs) have been described previously[3]:
BDP-Cl (8-chloromethyl-4,4′-difluoro-3,5-dimethyl-4-bora-3a,4a-diaza-s-indacene) and
COU-Br (4-(bromomethyl)-6,7-dimethoxycoumarin). Both compounds exhibit very low aqueous solubility, limiting the usable concentration range in enzymatic assays and necessitating the inclusion of organic cosolvents. The development of fluorogenic probes with improved water solubility would expand their applicability, enabling on-chip screening for HLDs and potentially facilitating live-cell activity in droplet-based assays such as fluorescence-activated droplet sorting (FADS).

Three novel fluorogenic substrates for HLDs with a coumarin core structure were synthesised to improve solubility. They were characterised by physicochemical properties, stability, steady-state kinetics with DmmA[4] and solubility in water. The results were then compared to the properties of BDP-Cl and COU-Br to deduce potential advantages of these probes.

*Table S2: Structures of the tested compounds based on the three core scaffolds COU-1, COU-2, and COU-3. The shortened names refer to the parent molecular scaffolds and denote the substrate forms bearing the –Cl group, whereas the –OH suffix indicates the corresponding hydrolysis products.*

| substrate | COU-1 | COU-2 | COU-3 |
|---|---|---|---|
| structure | 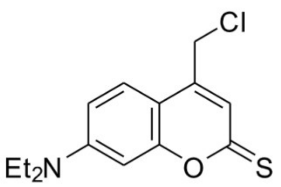 | 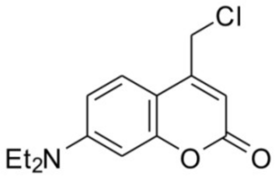 | 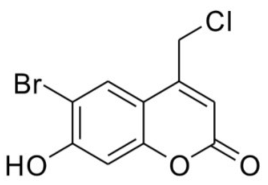 |
| Molecular weight | COU-1-Cl: 281.80<br>COU-1-OH: 263.36 | COU-2-Cl: 265.74<br>COU-2-OH: 247.29 | COU-3-Cl: 289.51 |

#### 2.1 Description of COU-2 synthesis

The reported procedure was used to prepare the COU-2[5].
7-(N,N-Diethylamino)-4-methylcoumarin (4.63 g, 20.0 mmol) was dissolved in p-xylene (120 mL). Then the selenium dioxide (3.6 g, 32.4 mmol, 1.5 eq) was added in one portion, and the mixture was heated to 140 °C for 20 h. The mixture was then filtered through a silica pad while hot to remove the black powder. The solvent was then removed in vacuo. The resulting brown oil was dissolved in ethanol (60 mL), and sodium borohydride (0.409 g, 10.8 mmol, 0.5 eq) was added in one portion. The reaction mixture was then stirred at RT for 18 h. Aqueous

HCl (5 %, 20 mL) was then added cautiously, and the mixture was extracted with dichloromethane (DCM; 3x 50 mL) and brine (50 mL). Organic layers were combined, dried with $MgSO_4$, filtered, and the solvent was removed in vacuo. Crude product was purified using flash column chromatography (silica gel, hexane/EtOAc 5/1 to1/1, then EtOAc to EtOAC/DCM 1/1, then DCM/acetone 5/1) to yield pure product as a yellow solid (385 mg, 7 %). $^1H$ NMR (400 MHz, DMSO-$d_6$) δ 7.43 (d, J = 9.0 Hz, 1H), 6.65 (dd, J = 9.0, 2.6 Hz, 1H), 6.51 (d, J = 2.6 Hz, 1H), 6.07 (t, J = 1.3 Hz, 1H), 5.48 (s, 1H), 4.66 (s, 2H), 3.42 (q, J = 7.0 Hz, 4H), 1.11 (t, J = 7.0 Hz, 6H). The $^1H$ NMR data were in agreement with the literature[6]. Spectroscopy data in phosphate buffer (pH = 8.0): absorption $\lambda_{max}$ = 387 nm (ε = 3900 L·mol$^{-1}$·cm$^{-1}$), emission $\lambda_{max}$ = 497 nm.

7-(N,N-Diethylamino)-4-(hydroxymethyl)-coumarin (0.15 g, 0.6 mmol) was dissolved in dry DCM (10 mL), and pyridine (0.064 mL, 0.79 mmol, 1.3 eq) was added. Tosyl chloride (0.347 g, 1.82 mmol, 3 eq) was added, and the solution was stirred at RT under nitrogen in the dark for 48 hours. Water (20 mL) was then added, and the reaction mixture was extracted with DCM (3x 20 mL). Organic layers were combined, dried with $MgSO_4$, filtered, and the solvent was removed in vacuo. The crude product was purified by flash column chromatography (silica gel, hexane/EtOAc 5/1) to yield pure product COU-2 as an orange solid (25 mg, 15 %). $^1H$ NMR (300 MHz, DMSO-$d_6$) δ 7.58 (d, J = 9.1 Hz, 1H), 6.74 (dd, J = 9.1, 2.5 Hz, 1H), 6.55 (d, J = 2.5 Hz, 1H), 6.20 (s, 1H), 4.90 (s, 2H), 3.44 (q, J = 7.0 Hz, 4H), 1.12 (t, J = 7.0 Hz, 6H). The $^1H$ NMR data were in agreement with the literature[6]. Spectroscopy data in phosphate buffer (pH = 8.0): absorption $\lambda_{max}$ = 389 nm (ε = 2500 L·mol$^{-1}$·cm$^{-1}$), emission $\lambda_{max}$ = 496 nm.

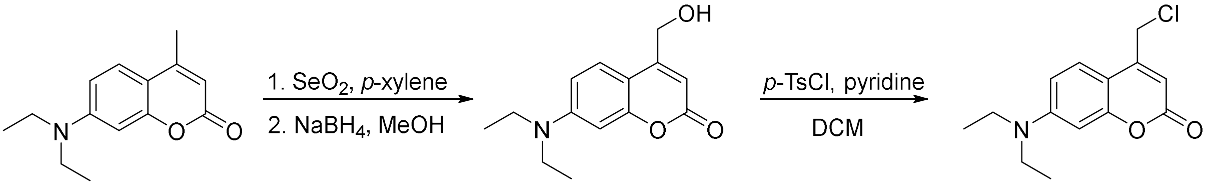

*Figure S8: Synthesis of COU-2 derivative (Molecular weight of 265.74 g·mol$^{-1}$).*

#### 2.2 Description of COU-1 synthesis

The reported procedure was used to prepare the COU-1[7].

7-(*N*,*N*-Diethylamino)-4-(chloromethyl)-coumarin (0.1 g, 0.38 mmol) and Lawesson's reagent (0.099 g, 0.24 mmol, 0.65 eq) were dissolved in dry toluene (40 mL) and stirred at 95 °C for 12 h. The solvent was removed in vacuo, and the crude product was purified by flash column chromatography (silica gel, hexane/EtOAc 6/1) to yield pure product COU-1 as a solid (80 mg,

75 %). $^{1}$H NMR (300 MHz, DMSO-$d_6$) $\delta$ 7.68 (d, $J$ = 9.1 Hz, 1H), 7.05 (s, 1H), 6.88 (dd, $J$ = 9.2, 2.5 Hz, 1H), 6.73 (d, $J$ = 2.4 Hz, 1H), 4.90 (s, 2H), 3.48 (q, $J$ = 7.0 Hz, 4H), 1.14 (t, $J$ = 7.0 Hz, 6H).

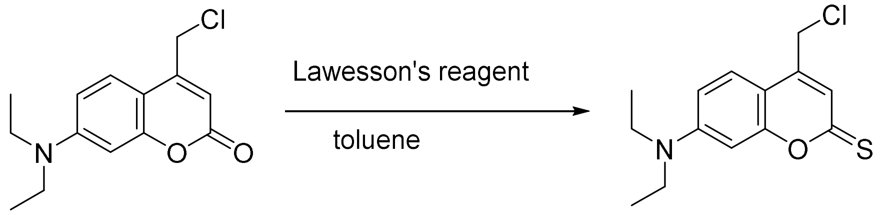

*Figure S9: Synthesis of COU-1 derivative (Molecular weight of 281.80 g·mol$^{-1}$).*

#### 2.3 Description of COU-3 synthesis

The reported procedure was used to prepare the COU-3[8]. Ethyl-4-chloroacetoacetate (1.6 mL, 11.9 mmol, 1.5 eq.) was dissolved in methanesulfonic acid (12 mL), and 4-bromoresorcinol (1.5 g, 7.94 mmol, 1.0 eq.) was added to the solution and stirred at 25 °C for 2.5 h. Then ice and water (50 mL) were added, and the suspension was filtered and washed several times with water. The solid was dried under reduced pressure to obtain crude product (2.05 g) as a beige solid. The crude product was purified using ethyl acetate (EtOAc) crystallisation (approximately 60 mL). The pure product was obtained as a beige solid (900 mg, 39 %). $^{1}$H-NMR (300 MHz, DMSO-$d_6$) $\delta$ 7.98 (s, 1H), 6.91 (s, 1H), 6.46 (s, 1H), 4.97 (s, 2H). The $^{1}$H-NMR data were in agreement with the literature[8]. Spectroscopy data in phosphate buffer (pH = 8.0): absorption $\lambda_{max}$ = 366 nm ($\varepsilon$ = 4600 L·mol$^{-1}$·cm$^{-1}$), emission $\lambda_{max}$ = 457 nm ($\Phi_{fl}$ = 0.49 ± 0.04 ($\lambda_{ex}$ = 372 nm, A($\lambda_{ex}$) < 0.1)) (*Figure S19* and *Figure S20*).

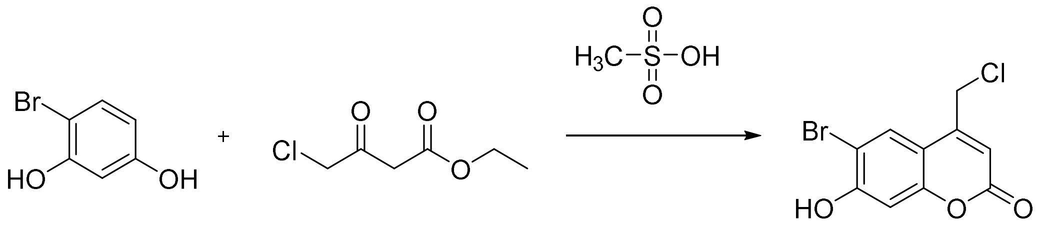

*Figure S10: Synthesis of COU-3 derivative (Molecular weight of 289.51 g·mol$^{-1}$).*

#### 2.4 Estimation of solubility and molar absorption coefficient

All substrates were dissolved to their maximum achievable concentration, which was then determined using UV/VIS spectroscopy. The results from solubility measurements are listed in *Table S3*. Only two compounds were successfully dissolved to a detectable concentration (COU-3, COU-2).

*Table S3: Solubility of tested compounds (detectable only for COU-2 and COU-3, n.d. – not determined).*

| substrate | COU-1 | COU-2 | COU-3 |
|---|---|---|---|
| **ε (l mol cm$^{-1}$)** | n.d. | 2500 | 4600 |
| **solubility (μM)** | n.d. | ~20 | ~290 |

Solubility tests ranked these substrates in the following order: COU-1 < COU-2 < COU-3, which shows a negative correlation with catalytic efficiency (COU-1 > COU-2 > COU-3).

A set of three stock solutions of COU-3 (0.70 mg, 0.88 mg, and 0.69 mg) in 2 mL of phosphate buffer (pH = 8.0) was prepared. A series of diluted samples (8x, 15x, 30x) was prepared, and absorption spectra were recorded. The molar absorption coefficient ($\varepsilon^{376}$ = 4600 L·mol$^{-1}$·cm$^{-1}$) was calculated as an average from the linear fits of absorbances at corresponding concentrations of the samples.

#### 2.5 Spectral properties

The excitation and emission spectra of all three coumarin substrates and products were measured in microtiter plates using a spectrofluorometer Synergy H4 (Biotek, USA) equipped with a xenon flash lamp and excitation and emission monochromators at 30 °C. The reaction mixture consisted of 90% phosphate buffer (pH 8.0) and 10% dimethyl sulfoxide (DMSO) with a total concentration of the substrates 80 μM (COU-1, COU-2) or 40 μM (COU-3).

*Figure S12* shows the excitation and emission spectra of COU-1-OH, COU-2-OH and COU-3-OH products. *Figure S13* shows the increase in quantum yields after conversion of substrates to products COU-1-OH, COU-2-OH and COU-3-OH. Product COU-3-OH was not synthesised, so the spectra were measured from the reaction mixture after total conversion. In *Table S4*, wavelengths corresponding to maximal excitation and emission are listed.

*Table S4: Maximal excitation and emission wavelengths of tested substrates.*

| substrate | $\lambda_{max,ex}$ (nm) | $\lambda_{max,em}$ (nm) |
|---|---|---|
| COU-1 | 475 | 559 |
| COU-2 | 389 | 496 |
| COU-3 | 366 | 457 |

A diluted ($A(\lambda_{ex}) < 0.1$) sample of COU-3 in phosphate buffer (pH = 8.0) in a fluorescence cuvette (1 cm) was prepared, and coupled emission spectra ($\lambda_{ex}$ = 372 nm) of solvent/sample were measured using an integration sphere in Edinburgh Instruments FLS920. The quantum yield was estimated to $\Phi_{fl}^{468}$ = (0.49 ± 0.04) as the average from three replicates (0.45; 0.49; 0.53).

#### 2.6 Stability

Stability of the probes was examined by measuring the increase in fluorescence intensity following spontaneous hydrolysis of 80 µM substrate solutions in 90% phosphate buffer (pH 8.0) with 10% DMSO after 2 h and 24 h. In between measurements, the substrates were kept in the dark (covered with aluminium foil) at room temperature.

*Figure S11* shows the increase in fluorescence intensity of stock solutions of the substrates 2 hours and 24 hours after dissolving them in DMSO. Especially in the case of COU-1 and COU-3, spontaneous hydrolysis leading to the corresponding product is evident. COU-2 is the most stable of the three, with less than 1% of product formed after 24 hours. All subsequent kinetic measurements were initiated 10 minutes after dissolving the compounds to ensure consistent results with the same percentage of product formed spontaneously.

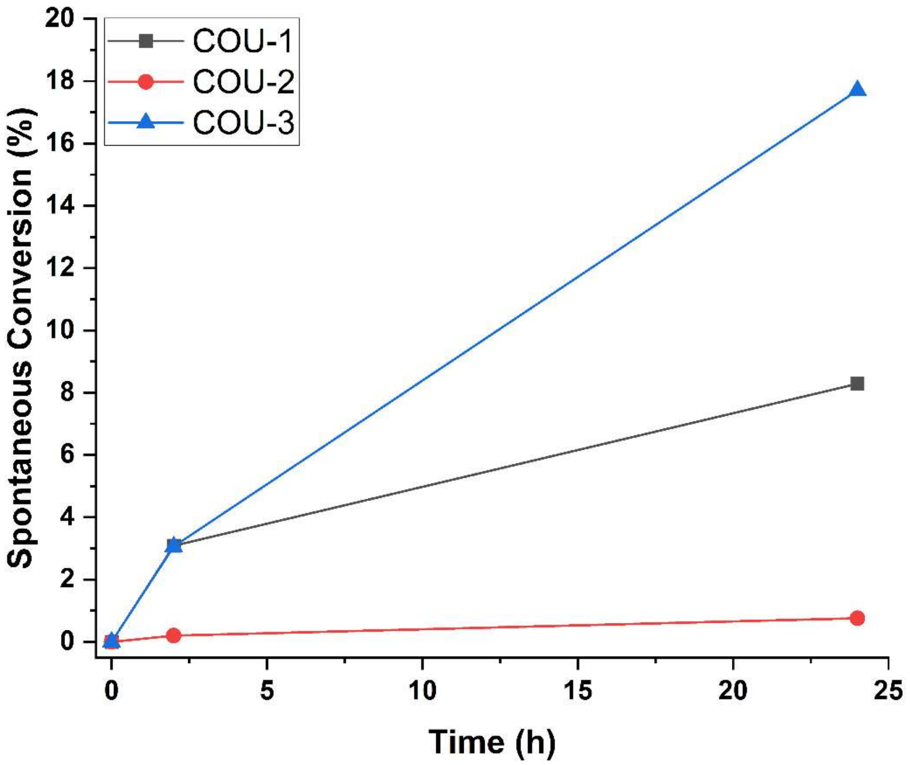

*Figure S11: Stability of COU-1, COU-2 and COU-3 in stock solutions over time plotted as % of conversion into product.*

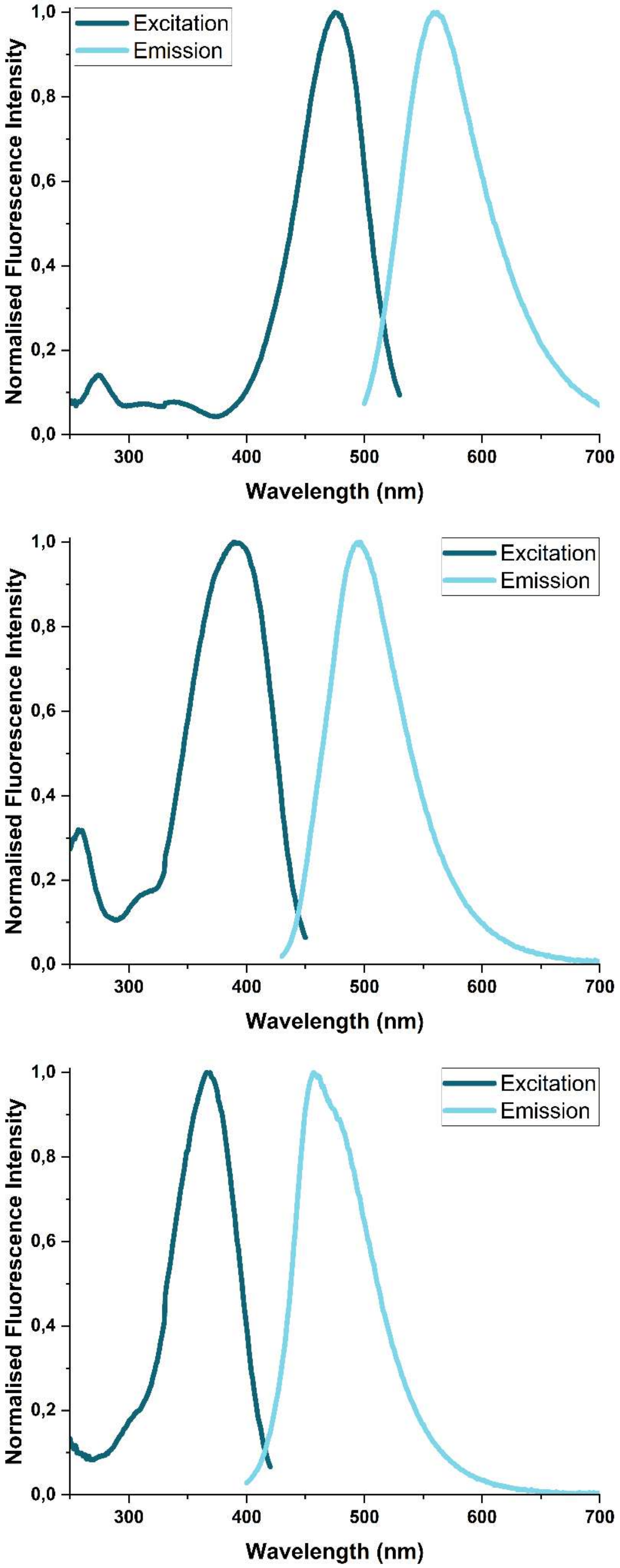

*Figure S12: Excitation (dark teal) and emission (light cyan) spectra of the coumarin-scaffold products (in the order from top). All fluorescence intensities are normalised to facilitate direct comparison of the spectral characteristics.*

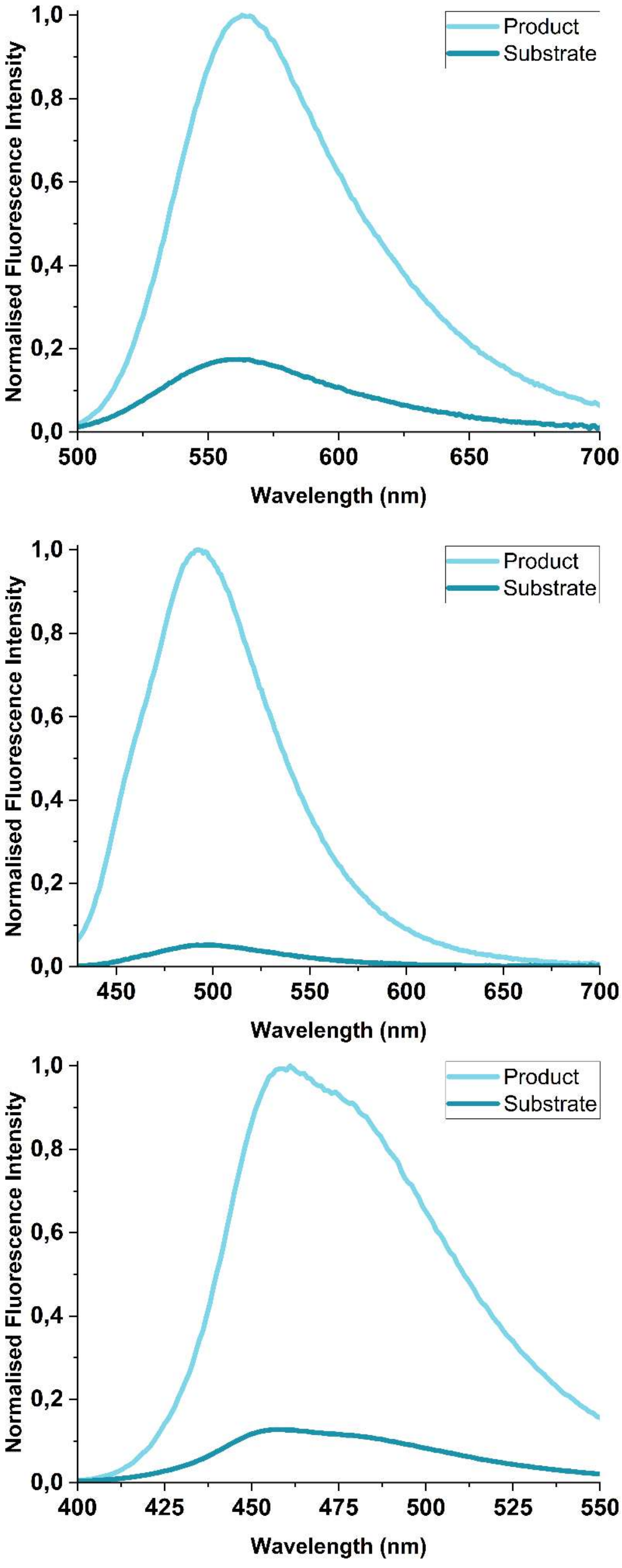

*Figure S13: Emission spectra of substrates COU-1, COU-2 and COU-3 (dark teal) and corresponding products COU-1-OH, COU-2-OH and COU-3-OH (light cyan). Spectra are sorted from top to bottom.*

#### 2.7 Steady-state kinetics

Coumarin probes were dissolved in DMSO to the appropriate concentration (tenfold higher than the concentration in the reaction mixture) 10 minutes before starting the measurement. The reactions were carried out in microtiter plates at 30 °C using a spectrofluorometer Synergy H4 (Biotek, USA) equipped with a xenon flash lamp. 20 µL of the substrate solution was added to the reaction mixture by an automatic syringe pump, bringing the final volume to 200 µL, with 90 % of 50 mM phosphate buffer (pH 8.0) containing the enzyme and 10 % of the substrate solution. Changes in the fluorescence intensity were measured from the top using excitation/emission monochromators set to the appropriate wavelengths (COU-1: 475/559, COU-2: 389/496, COU-3: 366/457) with a bandwidth of 9 nm. Before the first measurement, the microtiter plate was shaken for 2 s, and then the increase in fluorescence intensity was monitored at regular time intervals. The reaction was performed in 8 wells, along with 8 abiotic control wells, to monitor the spontaneous hydrolysis of the substrate.

The observed fluorescence signal was converted to product concentration using quadratic calibration curves derived from fluorescence intensities after total conversion. The fluorescence intensity increase caused by spontaneous hydrolysis of the substrates was subtracted from the converted data (**Equation 1**).

$$\mathrm{f}(\mathrm{c_p}) = \mathrm{FI} - \mathrm{f}(\mathrm{c_s})$$

*Equation 1: FI – observed fluorescence intensity; $f(c_p)$ – formula of a calibration curve for the product; $f(c_s)$ – formula of a calibration curve for the substrate.*

Global Kinetic Explorer[9,10] was used to analyse the steady-state data. The converted kinetic data for COU-2 and COU-3 were fitted to the reaction in Scheme 1 corresponding to the Michaelis-Menten model; for COU-1, a model accounting for substrate inhibition (Scheme 2) was used.

$$\mathrm{E} + \mathrm{S} \overset{K_\mathrm{m}}{\rightleftharpoons} \mathrm{ES} \xrightarrow{k_\mathrm{cat}} \mathrm{EP} \overset{K_\mathrm{i}}{\rightleftharpoons} \mathrm{E} + \mathrm{P}$$

*Scheme 1: $K_m$ – Michaelis constant, $k_{cat}$ – catalytic rate constant, $K_i$ – equilibrium dissociation constant of enzyme-product complex EP (measure of product inhibition).*

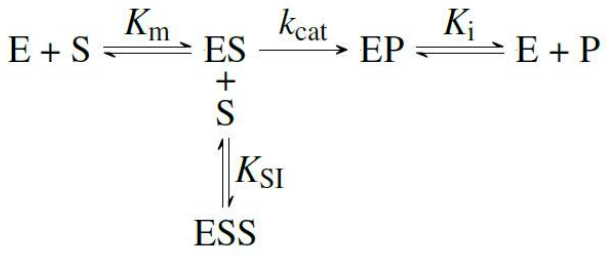

*Scheme 2: $K_m$ – Michaelis constant, $k_{cat}$ – catalytic rate constant, $K_i$ – equilibrium dissociation constant of enzyme-product complex EP (measure of product inhibition), $K_{SI}$ – equilibrium dissociation constant of non-productive complex ESS (measure of substrate inhibition).*

The reaction was carried out for several substrate concentrations, and the concentration of DmmA was fixed (low enough compared to substrate to maintain steady-state conditions). Calibration curves of concentration dependence of fluorescence intensity were constructed from the endpoints of each reaction by assuming 100% conversion of substrate into product (*Figure S14* - red, *Table S5*). For COU-1-OH and COU-2-OH, for which pure product samples were available, calibration curves were also constructed from these standards (*Figure S14* – purple).

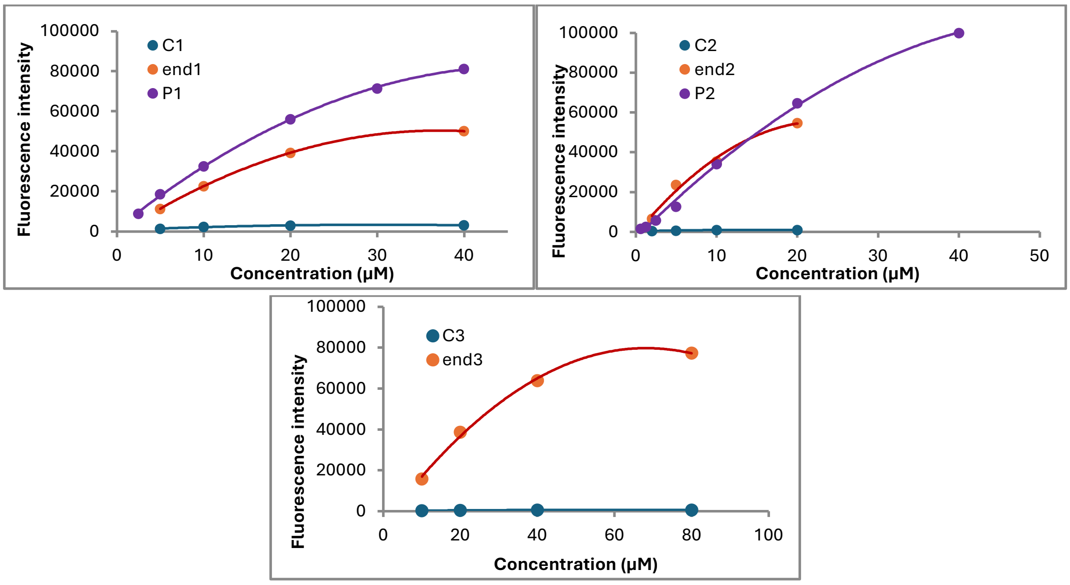

*Figure S14: Calibration curves of COU-1, COU-2 and COU-3 (in the reaction with DmmA constructed by quadratic fitting of fluorescence intensities corresponding to the endpoint of the reactions (red - end1, end2, end3) and spontaneous hydrolysis (blue – COU-1, COU-2, COU-3). Calibration curves of fluorescence intensity dependence on product concentration from pure product samples (purple – COU-1-OH, COU-2-OH, product sample of COU-3-OH was unavailable).*

*Table S5: Formulas for the calibration curves constructed by quadratic fitting of fluorescence intensities corresponding to the endpoint of each reaction.*

| **compound** | | **equation** | **$R^2$** |
|---|---|---|---|
| **COU-1** | substrate | $-2.9802x^2 + 179.46x + 642.98$ | 0.9469 |
| | product | $-37.736x^2 + 2803.2x - 1686.4$ | 1 |
| **COU-2** | substrate | $56.104x + 355.35$ | 0.9864 |
| | product | $-114.01x^2 + 5155.7x - 3054.8$ | 0.9865 |
| **COU-3** | substrate | $-0.1362x^2 + 16.822x + 100.92$ | 0.982 |
| | product | $-18.538x^2 + 2530.1x - 6563.2$ | 0.9971 |

The converted data were analysed in Global Kinetic Explorer[9,10]. The results of the best fits, including the FitSpace 1D analysis, are presented in *Figure S15*-*Figure* S17, and the derived kinetic constants and calculated steady-state parameters are in *Table S6*.

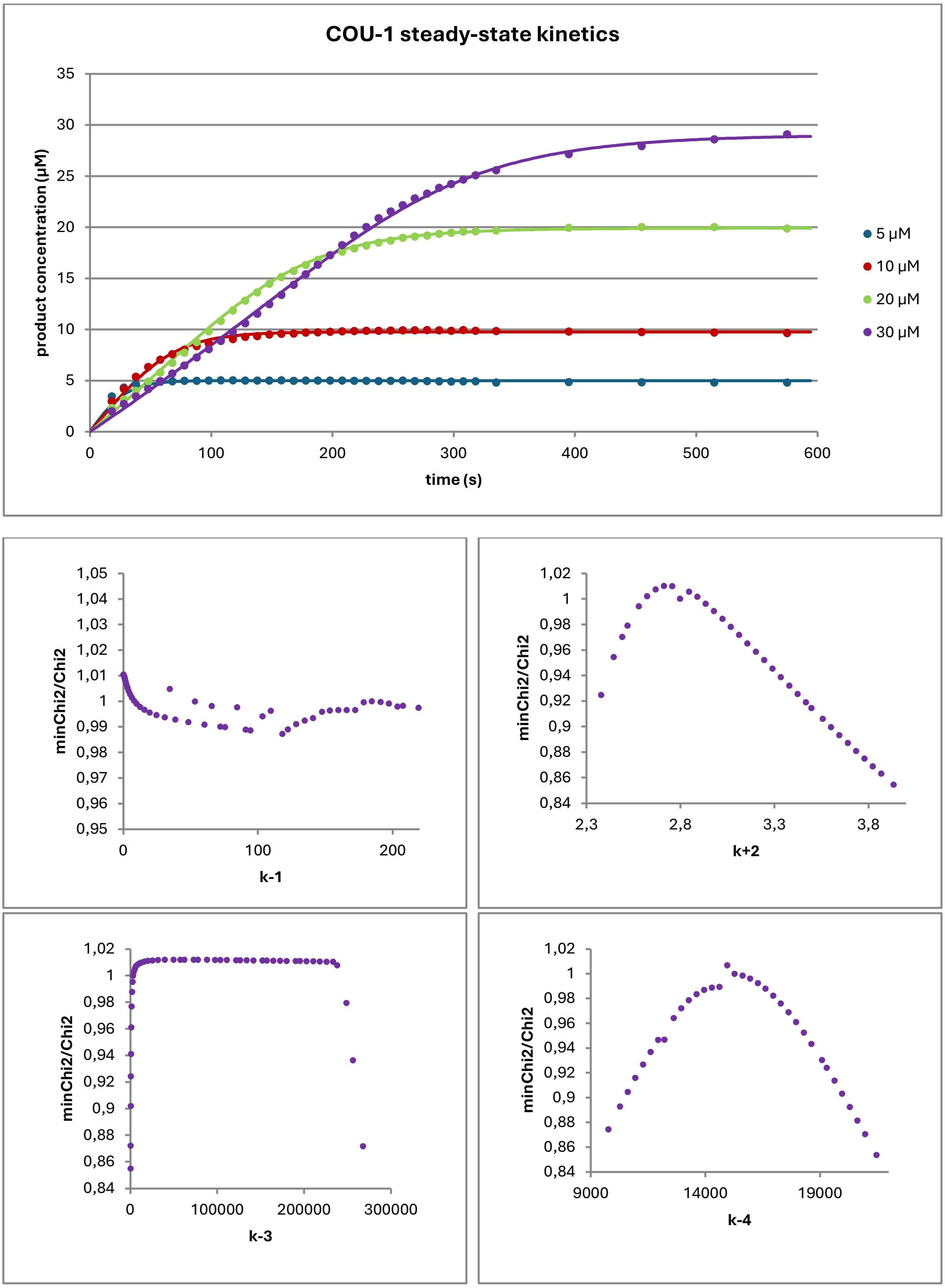

*Figure S15: Steady-state kinetic dataset for DmmA (0.083 µM) with different concentrations of COU-1. The lines represent the best fit to the proposed kinetic model (Scheme 2). Results of the FitSpace 1D analysis of the best fit. The graphs were obtained by fixing one parameter to different values, allowing the other parameters to float, and measuring the normalised χ2. χ2 is plotted as a function of the fixed parameter.*

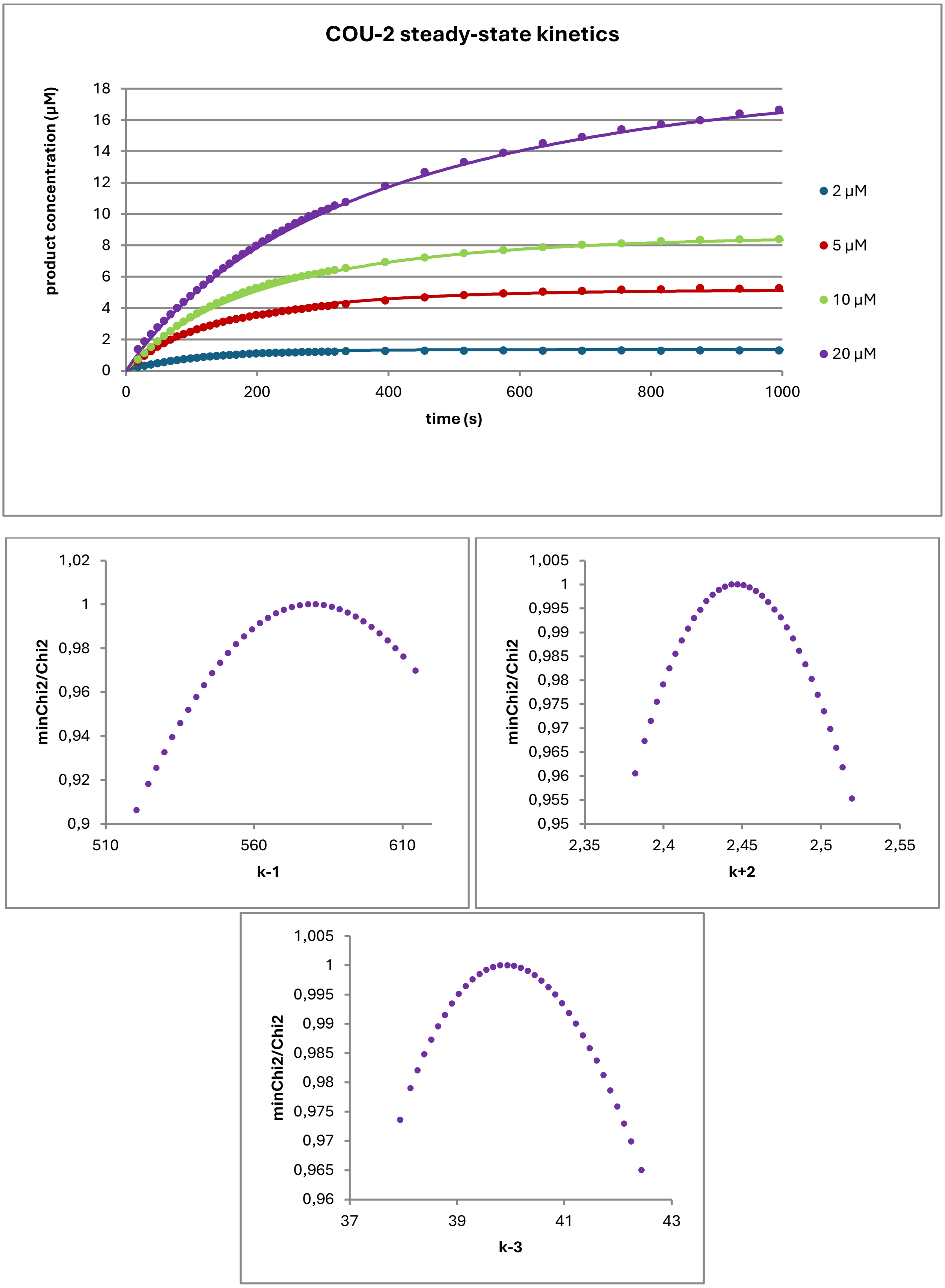

*Figure S16: Steady-state kinetic dataset for DmmA (0.034 µM) with different concentrations of COU-2. The lines represent the best fit to the proposed kinetic model (Scheme 1). Results of the FitSpace 1D analysis of the best fit. The graphs were obtained by fixing one parameter to different values, allowing the other parameters to float, and measuring the normalised χ2. χ2 is plotted as a function of the fixed parameter.*

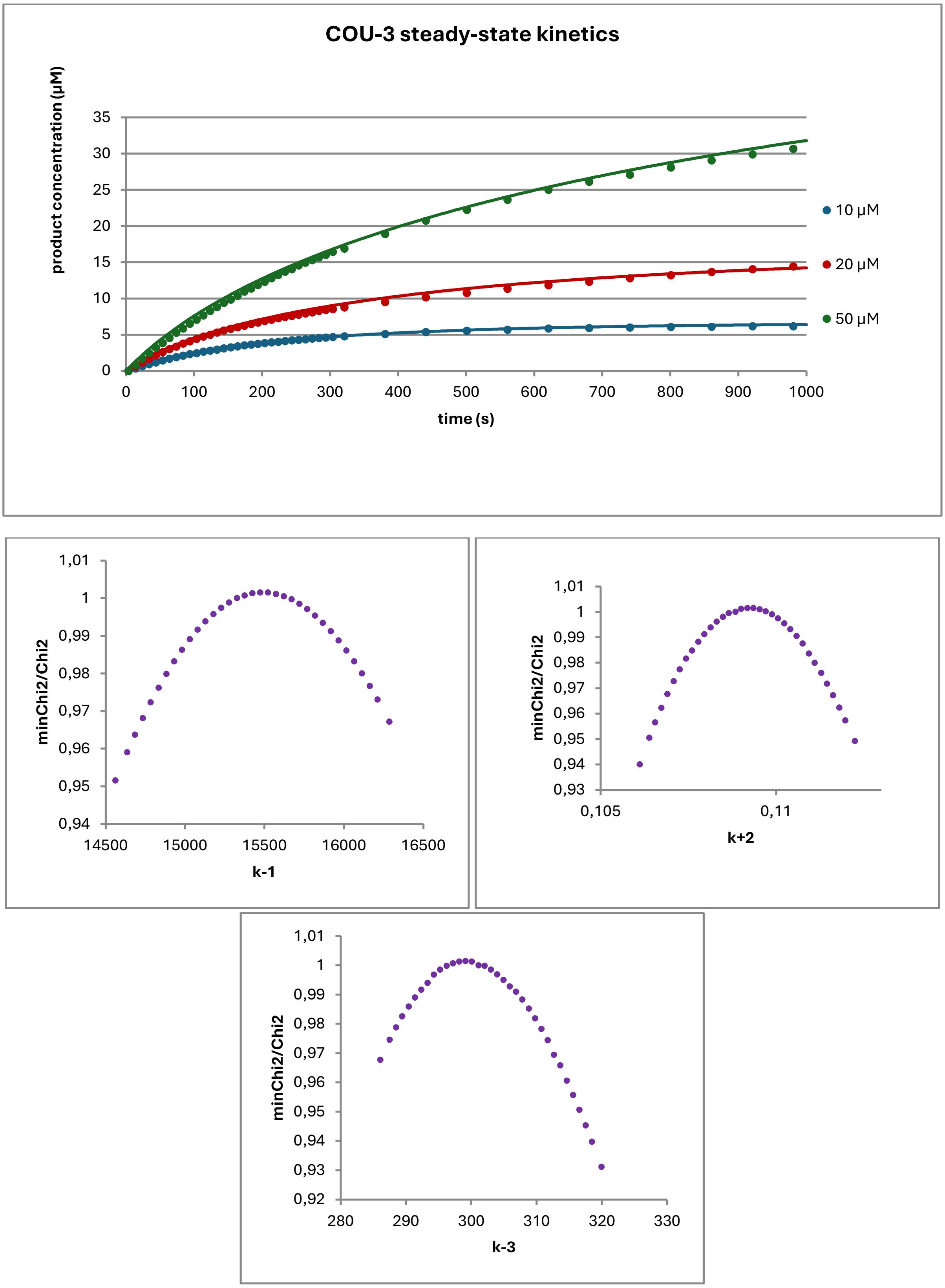

*Figure S17: Steady-state kinetic dataset for DmmA (1.18 µM) with different concentrations of COU-3. The lines represent the best fit to the proposed kinetic model (Scheme 1). Results of the FitSpace 1D analysis of the best fit. The graphs were obtained by fixing one parameter to different values, allowing the other parameters to float, and measuring the normalised χ2. χ2 is plotted as a function of the fixed parameter.*

*Table S6: Calculated steady-state parameters of DmmA with COU-1, COU-2 and COU-3 ($k_{cat}$ = k+2, Km = k-1/k+1, Ki = k+3/k-3, KSI = k-4/k+4). Standard errors are calculated by dividing standard deviations (sigma – sum of the residuals squared divided by the number of measurements minus one – noise quantification) by the square root of the number of measurements (which means that the estimated accuracy improves with the square root of the number of measurements, which may not always be accurate). $\chi 2$ is defined as the sum of squared residuals, normalised by the known sigma values. The $\chi 2$ threshold for calculation of lower and upper limits was set to 0.95.*

| substrate | parameter | value | StdErr | Lower limit | Upper limit |
|---|---|---|---|---|---|
| **COU-1** | $k_{cat}$ ($s^{-1}$) | 2.8 | 1.3 | 2.44 | 3.25 |
| | $K_m$ (μM) | 0.2 | - | - | 0.5 |
| | $K_i$ (μM) | 0.29 | - | 0.004 | 0.7 |
| | $K_{SI}$ (μM) | 15 | 10 | 12 | 19 |
| | **$k_{cat}$/ $K_m$ ($\mu M^{-1}$ $s^{-1}$)** | **14** | | | |
| **COU-2** | $k_{cat}$ ($s^{-1}$) | 2.4 | 0.1 | 2.38 | 2.52 |
| | $K_m$ (μM) | 5.8 | 0.6 | 5.4 | 6.1 |
| | $K_i$ (μM) | 2.50 | 0.272 | 2.4 | 2.6 |
| | **$k_{cat}$/ $K_m$ ($\mu M^{-1}$ $s^{-1}$)** | **0.41** | | | |
| **COU-3** | $k_{cat}$ ($s^{-1}$) | 0.109 | 0.005 | 0.106 | 0.112 |
| | $K_m$ (μM) | 15.3 | 1.53 | 14.6 | 16.3 |
| | $K_i$ (μM) | 3.32 | 0.32 | 3.15 | 3.50 |
| | **$k_{cat}$/ $K_m$ ($\mu M^{-1}$ $s^{-1}$)** | **0.0071** | | | |

*Table S7: Steady-state kinetic parameters of DmmA with COU-Br and BDP-Cl*[3]

| substrate | $k_{cat}$ ($s^{-1}$) | $K_m$ (μM) | $K_i$ (μM) | $K_{SI}$ (μM) | $k_{cat}$/ $K_m$ ($\mu M^{-1}$ $s^{-1}$) |
|---|---|---|---|---|---|
| COU-Br | 1.07 ± 0.08 | 0.77 ± 0.11 | 2.17 ± 0.14 | - | **1.4** |
| BDP-Cl | 3.5 ± 2.1 | 1.2 ± 0.8 | 0.40 ± 0.08 | 2.1 ± 1.5 | **2.9** |

In the case of COU-2 and COU-3, the specificity constant $k_{cat}/K_m$ is lower than that of COU-Br and BDP-Cl. For COU-3, substrate inhibition was observed. COU-1 shows a significant improvement in binding affinity, resulting in a higher specificity constant than COU-Br and BDP-Cl.

The enzymatic conversion of COU-2 and COU-3 was evaluated in a quartz cuvette containing a two-phase system composed of HFE-7500 oil and an aqueous buffer, designed to approximate the physicochemical conditions present in microfluidic droplets. Fluorescence measurements performed in the oil phase revealed substantial leakage of COU-2 into the HFE-7500 layer, indicating inadequate retention of this scaffold within the aqueous compartment. Owing to this pronounced partitioning behaviour, COU-2 was deemed unsuitable for droplet-based assays. In contrast, COU-3 exhibited markedly reduced leakage under identical conditions and was therefore selected as the preferred substrate for subsequent microfluidic experiments.

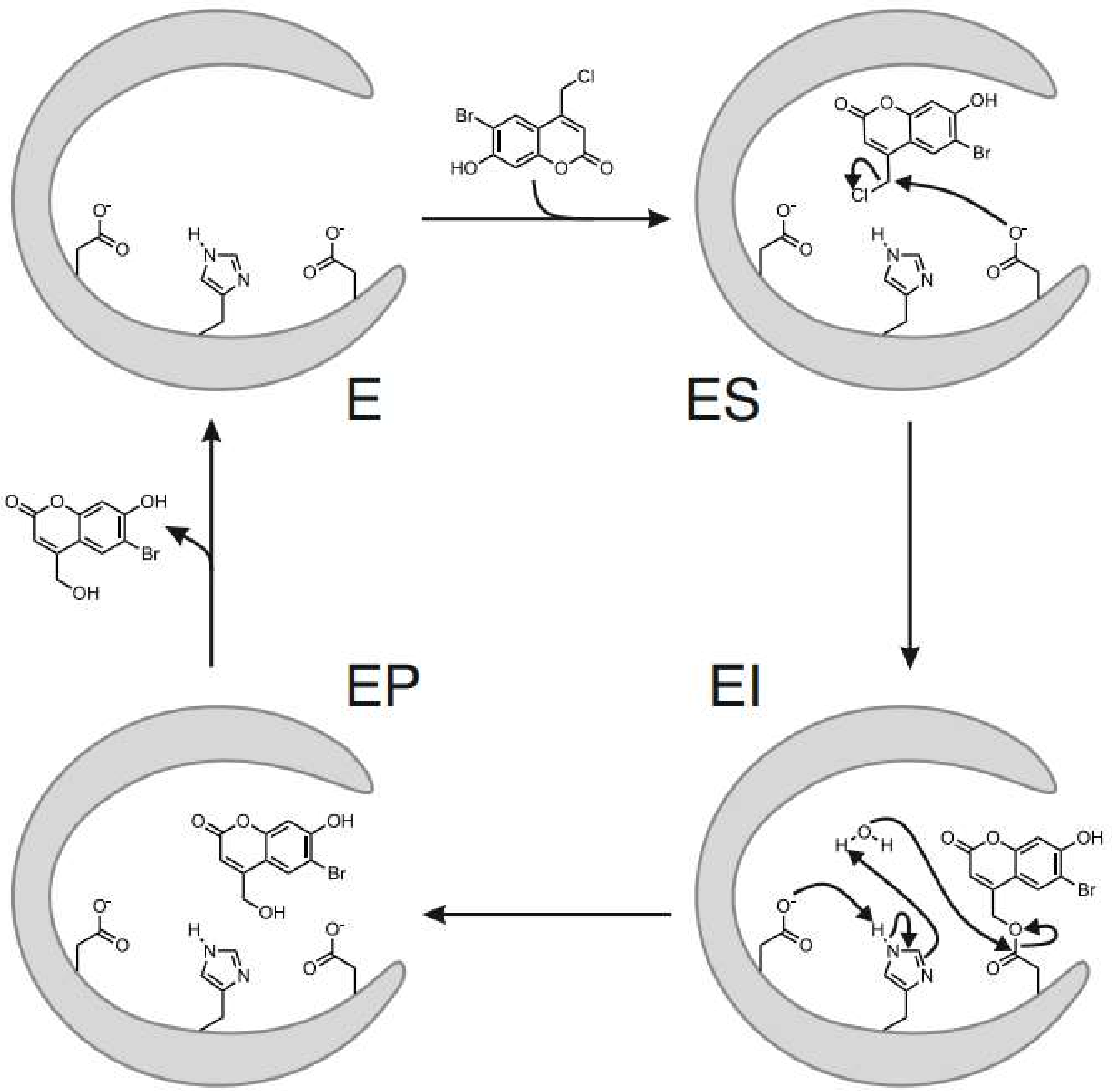

*Figure S18: Schematic reaction mechanism of the hydrolytic SN2 dehalogenation catalysed by a dehalogenase. The enzyme (E) first binds the halogenated substrate to form the enzyme–substrate complex (ES). In the initial $S_N2$ step, the catalytic nucleophile (typically an Asp residue) attacks the electrophilic carbon bearing the halogen, displacing the halide and generating a covalent ester intermediate (EI). A water molecule is subsequently activated by a catalytic base within the active site, enabling nucleophilic attack on the acyl-enzyme intermediate. This hydrolysis step releases the dehalogenated product and regenerates the free enzyme (EP → E), completing the catalytic cycle. Curved arrows represent electron flow during nucleophilic substitution and subsequent hydrolytic cleavage.*

#### 2.8 Stability of the COU-3 substrate

The stability of the COU-3 substrate was monitored over time to assess whether any degradation products might affect cellular DNA or cell viability. The initially synthesised batch of COU-3 was analysed by $^1$H-NMR at defined time points, using samples stored either at room temperature or at 4 °C. The initial $^1$H-NMR spectrum confirmed high purity, and spectra recorded up to 14 months later revealed no additional peaks indicative of decomposition, aside from a minor OH signal attributed to differences between DMSO batches. The strong agreement between the original and later spectra demonstrates that COU-3 remains chemically stable under the tested storage conditions, with no detectable degradation over the examined period.

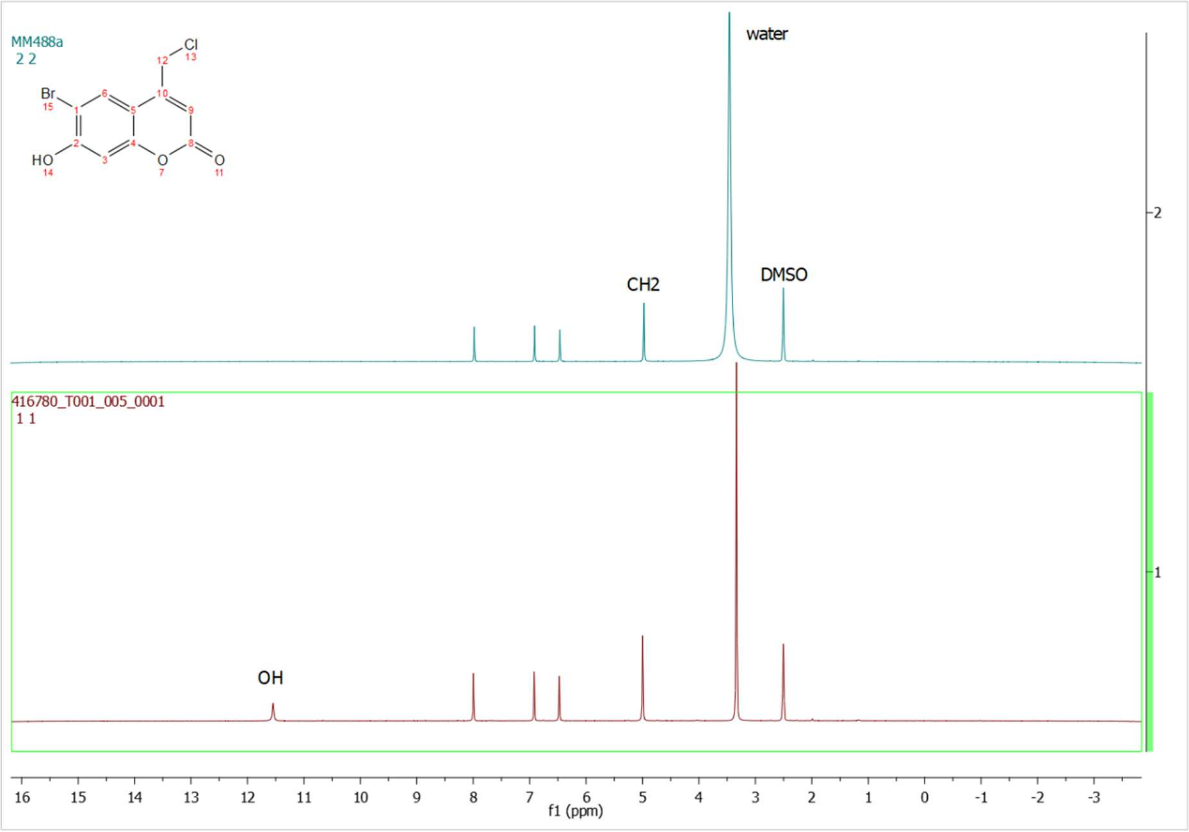

*Figure S19: $^1$H-NMR (300 MHz, DMSO-$d_6$) spectra of a freshly synthesised COU-3 (MM488a), and the same batch kept at room temperature for 14 months (sample 0001). The -OH group is probably given by a different batch of DMSO.*

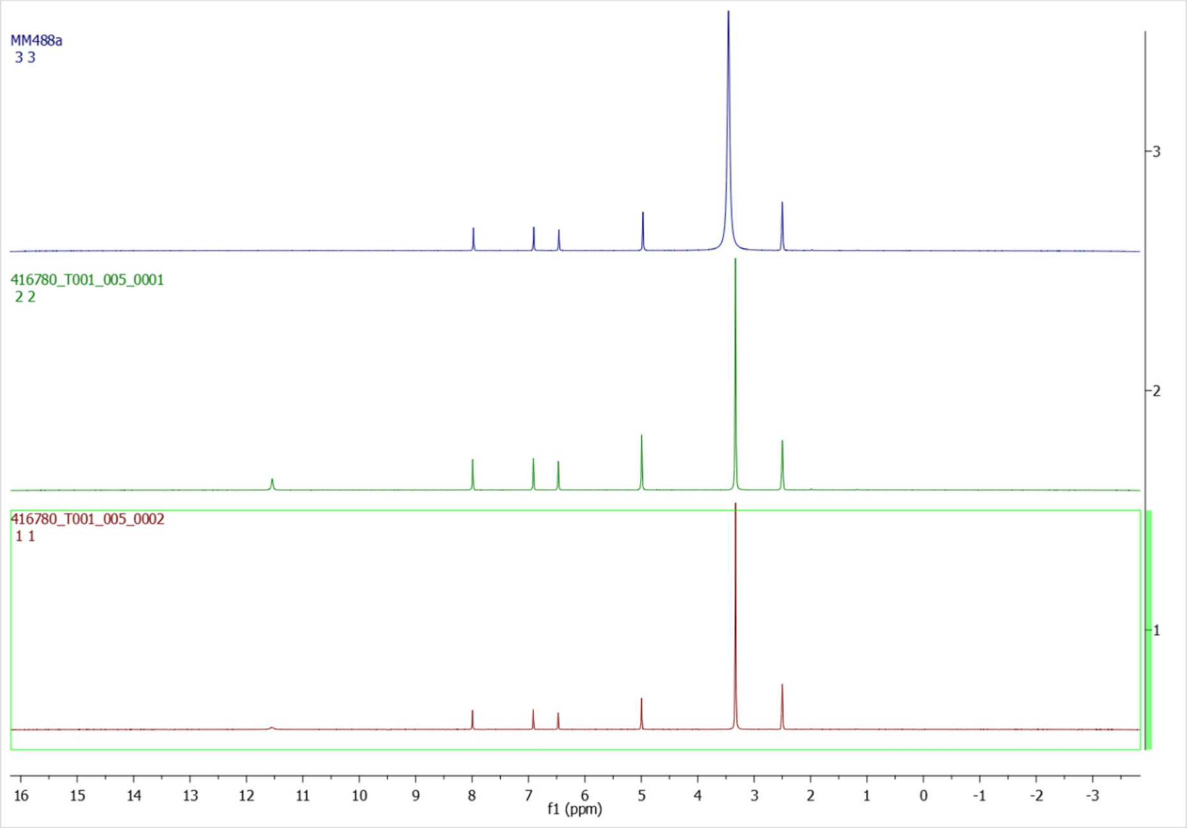

*Figure S20: ¹H-NMR (300 MHz, DMSO-d$_6$) spectra of a freshly synthesised COU-3 (MM488a), the second spectrum shows the same batch kept at room temperature (sample 0001), and the lower spectrum shows the batch kept at 4 °C for 14 months (sample 0002). The -OH group is probably given by a different batch of DMSO.*

### 3 Proteins library

#### 3.1 Generation of substitution libraries

Substitution libraries targeting the cap domain of LinB were prepared via error-prone PCR (epPCR) followed by Gibson Assembly (GA). The wild-type LinB gene was cloned into pET21b expression plasmid. Random substitutions were introduced into the cap domain following the GeneMorph II Random Mutagenesis kit (Agilent Technologies). The PCR conditions were optimised to achieve an approximate yield of 0.7-1.5 mutations per gene (*Table S9*). Primers flanking the cap domain were designed to amplify the target region (approximately 340 bp) with respect to the recognition sites necessary for the GA reaction. PCR reactions were run on the Thermal cycler GenePro (Bioer Technology Co.). PCR products were visualised by 1% agarose gel electrophoresis (Mini-PROTEAN Tetra electrophoresis system, Bio-Rad) and purified using the NucleoSpin Gel and PCR Clean-up kit (Macherey-Nagel). The fragment was assembled into a linearised vector using GA (*Table S10*). The linearisation was performed by digestion with BsrGI and XmaI restriction enzymes, followed by dephosphorylation and gel purification. Twenty microliters of GA reactions contained approximately 50 ng of linearised vector and 100 ng of epPCR product and were incubated at 50 °C for 1 hour (*Table S8*).

The assembled libraries were transformed into NEB 10-beta electrocompetent *E. coli* cells (New England Biolabs) using a BTX ECM 399 Exponential Decay Wave Electroporation System (Harvard Apparatus/BTX) with 2 µL of DNA, followed by recovery in SOC media at 37 °C for 90 min. Transformed cells were plated and used to inoculate overnight cultures (50 mL of LB media supplemented with 100 $\mu g \cdot mL^{-1}$ of ampicillin) for plasmid isolation using the NucleoSpin Plasmid kit (Macherey-Nagel). The quality as well as the approximate mutation frequency were evaluated by Sanger sequencing of random clones (Eurofins Genomics) (*Figure S21* and*Figure* S22).

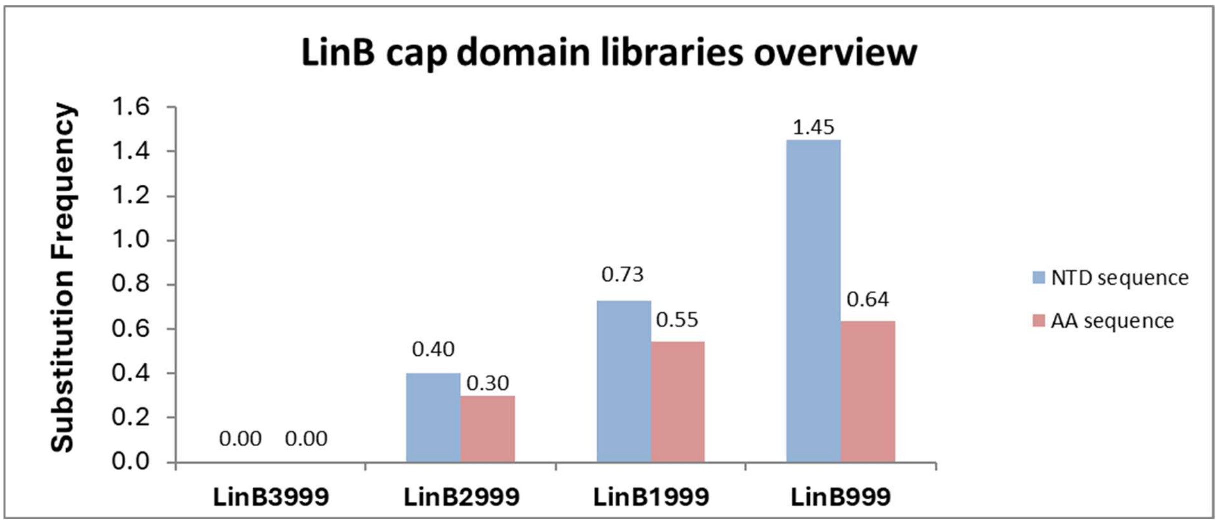

*Figure S21: Mutation frequency estimation libraries overview*

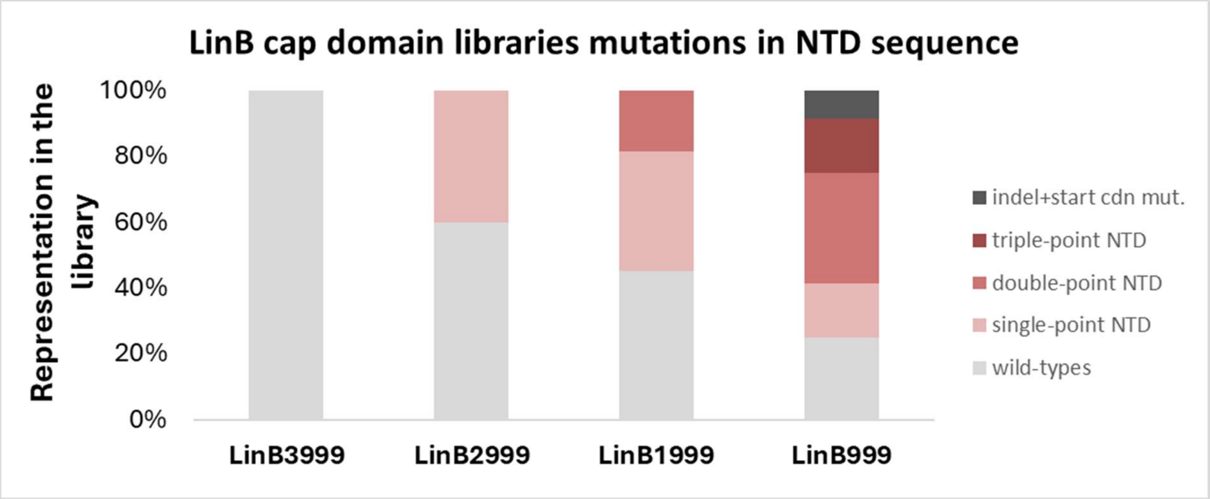

*Figure S22: Percentage representation of variants in the libraries*

*Table S8: Vector pET21b linearization*

| | Reaction mixture | | Conditions |
|---|---|---|---|
| | components | volume added | |
| Plasmid linearisation | CutSmart buffer | 5 µL | 37 °C, 2 hours |
| | DNA template (250 ng/µL) | 4 µL | |
| | BsrgI | 3 µL | |
| | XmaI | 3 µL | |
| | miliQ | 5 µL | |
| | | 20 µL | |

*Table S9: Parameters of the epPCR reactions during libraries generation.*

| | Reaction mixture | | | Conditions |
|---|---|---|---|---|
| | components | concentration/mass | volume added | |
| error-prone PCR Gene Morph II Random Mutagenesis | miliQ | - | 30.1/32.3/34.6/36.8 µL | 98 °C, 2 min |
| | Mutazyme II reaction buffer (1x) | - | 5 µL | 30x (98 °C, 30s; 65 °C, 30 s; 72 °C, 1 min) |
| | dNTPs (200 µM each) | 4 µM each | 1 µL | 72 °C, 10 min |
| | Fw primer (10 µM) | 0.2 µM | 1 µL | |
| | Rv primer (10 µM) | 0.2 µM | 1 µL | |
| | DNA template (457 ng/uL) | 215/160/125/54 ng | 8.9/6.7/4.4/2.2 µL | |
| | Mutazyme II DNA polymerase (2.5 U) | - | 1 µL | |
| | | | 50 µL | |

*Table S10: Gibson assembly reactions during libraries generation.*

| | Reaction mixture | | | Conditions |
|---|---|---|---|---|
| | components | stock concentration | volume added | |
| Gibson assembly | miliQ | - | 1.3/1.6/1.2/0.9 µL | 50 °C, 1 hour |
| | GA master mix | - | 15 µL | |
| | Linearised vector | 57.8 ng/µL | 0.9 µL | |
| | Cap domain after epPCR (LinB3999/2999/1999/999) | 46.6/51.5/44.3/40.3 ng/µL | 2.8/2.5/2.9/3.2 µL | |
| | | | 20 µL | |

### 4 On-chip sorting

#### 4.1 Cells cultivation, protein production, and samples preparation for the sorting

Chemically competent *E. coli* BL21(DE3) cells (New England Biolabs, USA) were transformed with the substitution library LinB999 by heat-shock at 42 °C for 30 s, followed by recovery in SOC media at 37 °C at 190 rpm for 1 h. After cultivation, the cells were transferred to LB medium (10 mL supplemented with 100 μg·mL$^{-1}$ of ampicillin), grown for 16 h at 37 °C and 190 rpm, and protein expression was induced with 0.5 mM IPTG, followed by 18 h of protein production at 20 °C and 120 rpm. Cells were harvested (4,000 g, 30 min, 4 °C), washed twice with PB buffer (20 mM phosphate buffer (pH 8.0)), concentrated, aliquoted, and stored at −20 °C.

#### 4.2 Cells encapsulation and incubation of droplets

A 20 mM stock solution of the COU-3 substrate was prepared in DMSO to a final volume of 1 mL. The working solution (100 μM) was generated by diluting this stock in PB buffer to obtain a final DMSO concentration of 0.5%. A bacterial suspension was prepared by combining 125 μL of OptiPrep (ab286850, Abcam, UK) with 375 μL of PB buffer, followed by filtration through a 0.2 μm PVDF membrane filter. From the filtrate, 396.8 μL was mixed with 3.2 μL of *E. coli* cells expressing the plasmid-encoded LinB mutant library, providing a source of distinct enzyme variants for encapsulation.

For cell encapsulation and primary emulsion formation, fluorinated oil (HFE-7500) supplemented with 1.25% (w/w) RAN008 surfactant was introduced into a 1 mL Hamilton Gastight syringe, while the bacterial matrix and COU-3 working solution were each loaded into separate 250 μL Hamilton Gastight syringes. Fluids were supplied to the microfluidic device at flow rates of 335 μL·h$^{-1}$ for the continuous oil phase and 30 μL·h$^{-1}$ for both the cell-containing matrix and the COU-3 solution. Droplet generation conditions were kept constant, and the resulting primary emulsion was collected in silanised vials over 2 h.

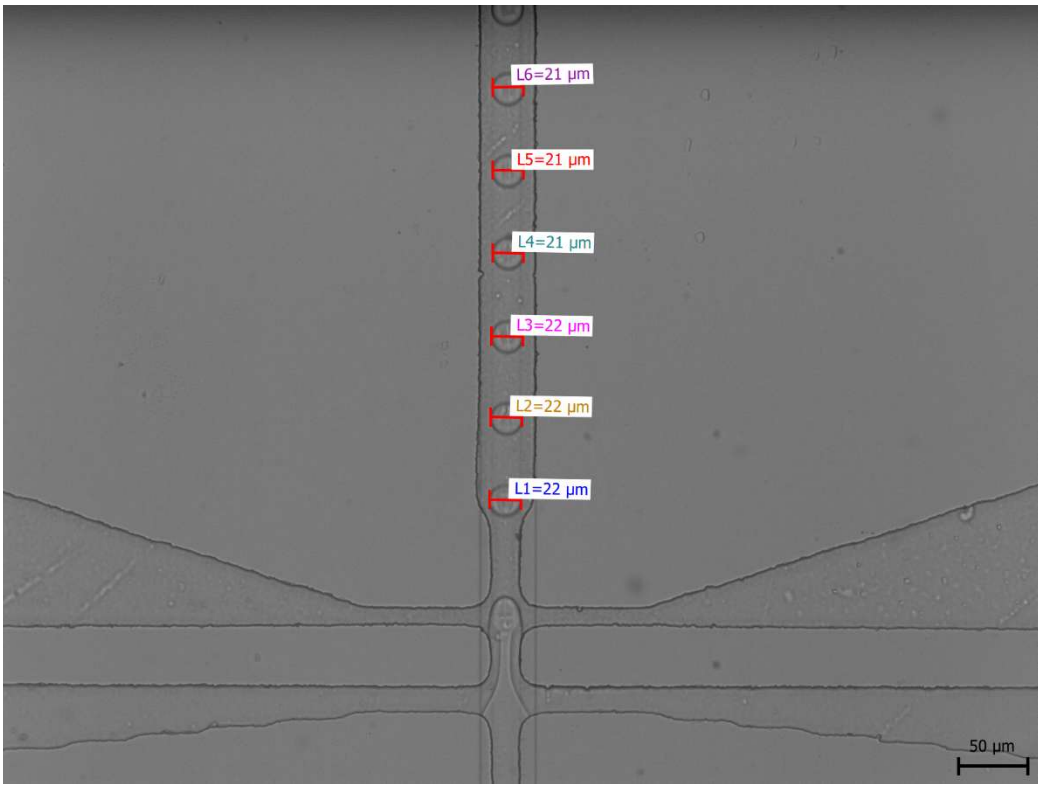

*Figure S23: Generation of droplets (LinB library in E. coli BL21(DE3) in the presence of COU-3 substrate. Image acquired by QuickPHOTO CAMERA 3.2 software.*

##### 4.3 FADS sorting of LinB999 Library

The sorting process represents the culmination of our workflow and requires meticulous optimisation of multiple parameters. Spacing oils were introduced at carefully calibrated flow rates (8 $\mu L \cdot h^{-1}$ for spacing oil 1 and 15 $\mu L \cdot h^{-1}$ for spacing oil 2) to ensure proper separation between droplets during analysis. The pressure for droplet reinjection was set to 65 mBar, balancing droplet integrity and sufficient throughput. Dielectrophoretic pulse parameters were also optimised, with an amplitude of 8 V (8 kV after amplification), a frequency of 30,000 Hz, a burst count of 40 cycles, a delay of 60 µs, and a laser intensity of 100 % of maximum power (1005 mW for the 405 nm laser).

Using the pre-sorting analysis in Cinderella[A] tool, the threshold of 0.6 V was determined to be set for the sorting. Three portions of sorted droplets, each containing 600–1000 droplets, were collected to provide optimal samples for subsequent PCR amplification (*Table S11*).

*Table S11: Properties of the three sorted portions of droplets. Only droplets containing cells with an active enzyme are detectable. The empty droplets or inactive variants are not visible to the PMT detector; therefore, the overall number of droplets that passed through the sorting chip is approximately 10-20× higher than the number detected by the PMT because λ = 0.05.*

| Sorted samples | Sorted into | Sorted number of droplets | From the total detected number | Percentage of sorted droplets* |
|---|---|---|---|---|
| Sample01<br>Gain 1.1 V<br>Threshold 0.6 V<br>Positive oil 15 μL · h$^{-1}$<br>Negative oil 8 μL · h$^{-1}$<br>Droplets: 40 mbar | Master Mix (PCR) | 1000 | 11.254.073 | 0.09 % |
| Sample02<br>Gain 1.1 V<br>Threshold 0.6 V<br>Positive oil: 15 μL · h$^{-1}$<br>Negative oil: 8 μL · h$^{-1}$<br>Droplets: 50 – 67 mbar | Master Mix (PCR) | 1000 | 16.211.221 | 0.006 % |
| Sample03<br>Gain 1.1 V<br>Threshold 0.6 V<br>Positive oil 15 μL · h$^{-1}$<br>Negative oil 8 μL · h$^{-1}$<br>Droplets: 67 mbar | Master Mix (PCR) | 600 | 10.020.909 | 0.006 % |

** Sorted droplets from the population of total number of detected droplets having fluorescence emission*

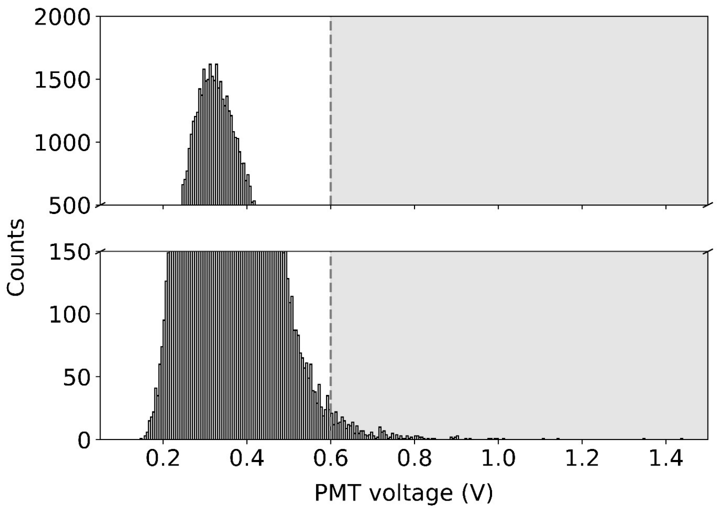

*Figure S24: Histogram of the first portion of sorted droplets.*

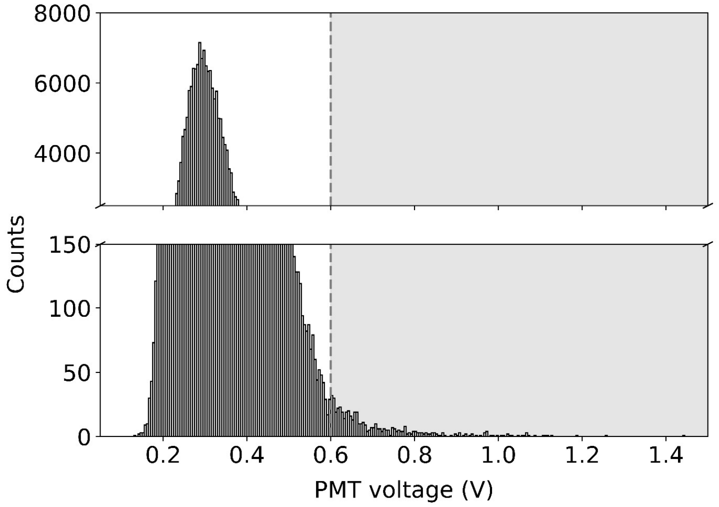

*Figure S25: Histogram of the second portion of droplets.*

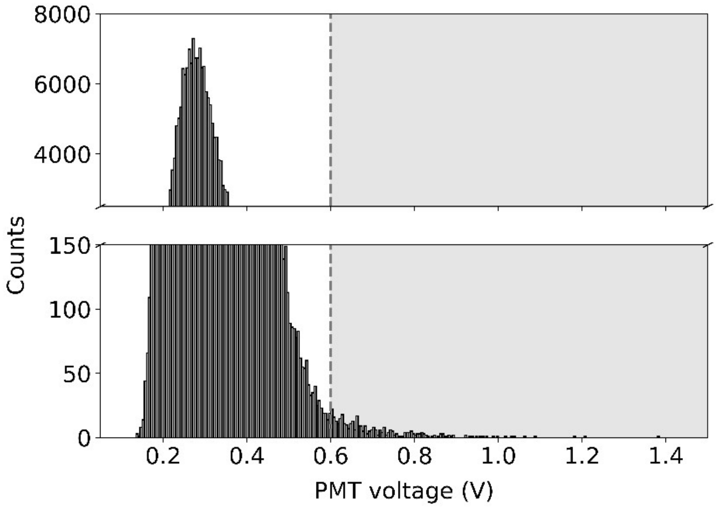

*Figure S26: Histogram of the third portion of sorted droplets.*

### 5 Hits generation:

#### 5.1 DNA recovery of the plasmid DNA from the sorted-out cells

For DNA recovery, PCR was performed with the following primers:
5′-GAAGGAGATATACATATGAGC-3′ and reverse
5′-CGAGTGCGGCCGCAAGCTTTCAG-3′.
During sorting, cells were collected into a PCR master mix composed of milliQ water, primers, and Verifi polymerase buffer (PCR Biosystems). The composition of the PCR reaction is summarised in *Table S12*. Verifi polymerase (PCR Biosystems) was added after the first denaturation step (98 °C; 30 s; 1x), followed by cooling of the whole mixture to 10 °C. After addition of the polymerase, the PCR consisted of one cycle of initial denaturation (95 °C, 1min) followed by thirty-five cycles of 95 °C (denaturation, 15 s), 58 °C (annealing, 15 s) and 72 °C (extension, 40 s) and one cycle of final extension at 72 °C for 8 min (*Table S12*). PCR was followed by agarose electrophoresis; the entire PCR mixture was mixed with 10 µl Gel Purple Loading Dye (6x) (New England Biolabs), and a 1% gel was run at 120 V, 400 mA for 35 minutes. Corresponding bands were cut out and extracted from the gel using NucleoSpin Gel and PCR Clean-up kit (item no. 740609; Macherey-Nagel).

The Gibson assembly (GA) reaction was used to assemble PCR fragments into the pET21b vector. The pET21b vector was linearised with the restriction enzymes NdeI (New England Biolabs) and HindIII (New England Biolabs); reverse linearization of the vector was prevented by the addition of Antarctic phosphatase (New England Biolabs) after cleavage. Approximately 100 ng of DNA fragment isolated from the gel and about 50 ng of linearised vector backbone pET21b were added to the thawed 15 µL in-house prepared Gibson assembly master mix in a total reaction volume of 20 µL. The whole mixture was then incubated at 50 °C for 1 h. After incubation, the GA mixture was cooled, then transformed into in-house prepared *E. coli* DH5α (200 µL of thawed cells per reaction) via heat shock (1 min, 42 °C). The cells were recovered in 600 µL of pre-heated SOC medium followed by 2 h incubation (37 °C, 190 rpm). After incubation, the cells were spread onto LB-agar plates containing 100 µg·mL$^{-1}$ of ampicillin and incubated overnight at 37 °C.

*Table S12: Parameters of the DNA recovery PCR conditions.*

| | Reaction mixture | | | Conditions |
|---|---|---|---|---|
| | components | concentra-tion/mass | volume added | |
| DNA rescue PCR | miliQ | - | 35.4 µL | 98 °C, 2 min |
| | Verifi polymerase buffer (5x) | 1x | 10 µL | 35x (95 °C, 15s; 60 °C, 15 s; 72 °C, 40s) |
| | dNTPs (200 µM each) | 4 µM each | 1 µL | 72 °C, 8 min |
| | Fw primer (10 µM) | 0.2 µM | 2 µL | |
| | Rv primer (10 µM) | 0.2 µM | 2 µL | |
| | DNA template (cells after sorting) | - | 0.1 µL | |
| | Verifi polymerase (2 U) | - | 0.5 µL | |
| | | | 50 µL | |

#### 5.2 Large-scale cultivation and purification of the hits

Recombinant plasmids (pET21b::linB variants and empty vector) were transformed into *E. coli* BL21(DE3) cells via heat shock (42 °C, 30 s). After recovery in SOC media (37 °C, 190 rpm, 1 h), the cells were plated on LB agar supplemented with 100 µg·mL$^{-1}$ of ampicillin and grown overnight at 37 °C. Ten colonies per construct were used to inoculate 20 mL of LB medium (supplemented with 100 µg·mL$^{-1}$ of ampicillin), which was incubated at 37 °C and 180 rpm for 4 h. Each 1 L LB culture was inoculated with 10 mL of preculture and grown at 37 °C, 150 rpm until $OD_{600}$ reached approximately 0.8 (*Table S13*). Protein production was induced with IPTG (50 µM), and the cells were incubated overnight at 20 °C, 120 rpm. The cells were harvested (5,000 g, 40 min, 4 °C), resuspended in 40 mL of harvesting buffer (16 mM $K_2HPO_4$, 3.6 mM $KH_2PO_4$, 5 mM imidazole, pH 7.5), and stored at −70 °C.

Prior to purification, cultures were thawed, and 90 µL of DNase (1 mg·mL$^{-1}$) was added. After sonication (MODEL FB705, Thermo Fisher Scientific, USA) in 3× 2 min cycles with a 50% amplitude (5 s pulse, 5 s pause), clarification of lysates was achieved via centrifugation (19,500 g, 4 °C, 1 h) using an Eppendorf centrifuge (5910Ri G, Eppendorf, Germany) equipped with a Rotor FA-6x50.

The supernatant was filtered through a 0.45 µm Millipore membrane filter (Merck, Germany) prior to purification by metal ion affinity chromatography using a HisTrap Excel column on the ÄKTA Pure system (Cytiva, USA). Proteins were eluted using a linear gradient (0-100%) with a buffer containing 20 mM phosphate, 500 mM NaCl, and 500 mM imidazole at pH 7.5, followed by dialysis against 50 mM phosphate buffer (pH 7.5).

Fractions were collected, concentrated, and further purified via size exclusion chromatography on the same ÄKTA Pure system, utilising a HiLoad 26/600 Superdex 200 pg column (Cytiva, USA) in PB buffer (pH 8.0). Purified proteins were concentrated to a final concentration of approximately 5 mg·mL$^{-1}$ (NanoDrop One) using Centrifugal Filter Units (Amicon R Ultra-15 Ultracel R -10K, Merck Millipore Ltd.) and were flash-frozen in liquid nitrogen and stored at −70 °C or lyophilised.

*Table S13: Cultivation, protein production, and purification overview, including the final yields for individual protein variants.*

| VARIANT | CULTIVATION | | PURIFICATION IMAC | | PURIFICATION SEC | | |
|---|---|---|---|---|---|---|---|
| | $OD_{600}$ before IPTG | $OD_{600}$ after IPTG | concentration | volume | concentration | volume | **final yield** |
| **LinB WT** | 1.22 | 5.20 | 10.5 mg·mL$^{-1}$ | 5 mL | 5.1 mg·mL$^{-1}$ | 7.4 mL | **18.87 mg·L$^{-1}$** |
| **P208S** | 1.00 | 4.70 | 11.7 mg·mL$^{-1}$ | 5 mL | 5.1 mg·mL$^{-1}$ | 9.3 mL | **23.59 mg·L$^{-1}$** |
| **R209L** | 0.84 | 5.04 | 6.7 mg·mL$^{-1}$ | 5 mL | 5.1 mg·mL$^{-1}$ | 4.8 mL | **12.24 mg·L$^{-1}$** |
| **I138N** | 0.76 | 4.64 | 2.0 mg·mL$^{-1}$ | 5 mL | 5.0 mg·mL$^{-1}$ | 0.7 mL | **1.75 mg·L$^{-1}$** |
| **P137S** | 0.86 | 5.80 | 2.3 mg·mL$^{-1}$ | 5 mL | 4.9 mg·mL$^{-1}$ | 2.0 mL | **4.90 mg·L$^{-1}$** |
| **V173F** | 0.96 | 5.60 | 5.5 mg·mL$^{-1}$ | 5 mL | 5.0 mg·mL$^{-1}$ | 7.3 mL | **18.25 mg·L$^{-1}$** |

### 6 Experimental Hits Characterisation

#### 6.1 Thermal stability

Thermal stability of the LinB variants was evaluated by nano differential scanning fluorimetry (nanoDSF) on a Prometheus Panta instrument using nanoDSF-grade standard capillaries. Thermograms were acquired from 20 to 95 °C at a heating rate of 1 °C·min$^{-1}$. Protein stock solutions were passed through a 0.2 µm PVDF filter and diluted to a final concentration of approximately 1 mg·mL$^{-1}$ in 50 mM potassium buffer (pH 7.5). Fluorescence was excited at 280 nm, and emission was recorded simultaneously at 330 and 350 nm with a dual-UV detector. Data were processed using Panta Analysis software, and the melting temperature (Tm) was defined as the inflexion point of the fluorescence intensity ratio (330/350 nm) as a function of temperature, reporting changes in the local environment of aromatic residues.

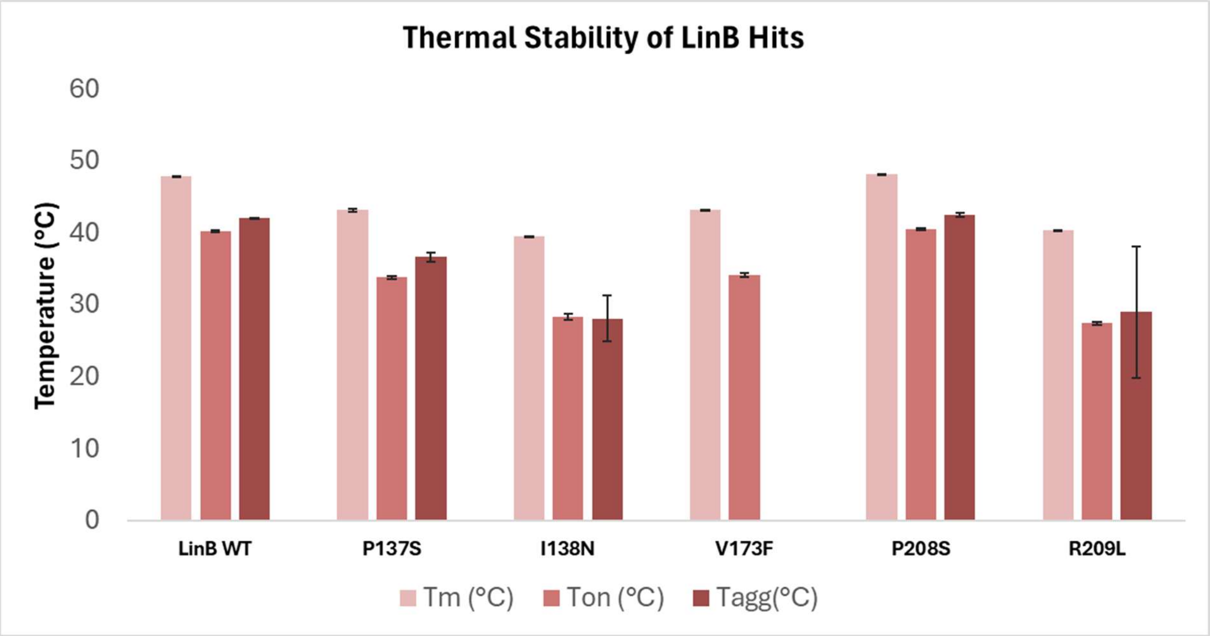

*Figure S27: Comparison of thermal stabilities of LinB variants measured via nanoDSF – melting temperature (Tm), onset temperature of unfolding (Ton), and aggregation temperature (Tagg).*

*Table S14: Means and standard deviation (SD) values of thermal stabilities obtained from nanoDSF.*

| VARIANT | Tm (°C) | | Ton (°C) | | Tagg(°C) | |
|---|---|---|---|---|---|---|
| | Mean | SD | Mean | SD | Mean | SD |
| **LinB WT** | 47.94 | 0.02 | 40.29 | 0.15 | 42.12 | 0.09 |
| **P137S** | 43.24 | 0.17 | 33.89 | 0.26 | 36.74 | 0.65 |
| **I138N** | 39.62 | 0.05 | 28.44 | 0.39 | 28.16 | 3.21 |
| **V173F** | 43.35 | 0.02 | 34.24 | 0.30 | - | - |
| **P208S** | 48.22 | 0.10 | 40.62 | 0.15 | 42.62 | 0.31 |
| **R209L** | 40.50 | 0.04 | 27.51 | 0.15 | 29.11 | 9.15 |

##### 6.2 Steady-state kinetics

A potassium phosphate (PB) buffer (20 mM) was freshly prepared and filtered through a 0.2 µm membrane filter (Macherey-Nagel, catalogue no. 659020047) prior to use.

Enzyme stock solutions were filtered through a 0.2 µm PVDF membrane filter (Cytiva, catalog no. 6777-0402) before measurements. Protein concentrations were determined using a NanoDrop One microvolume UV–Vis spectrophotometer (Thermo Fisher Scientific, USA) and subsequently diluted to the desired final concentrations. Final enzyme concentrations were either 125 nM (wild-type, I138N, P137S) or 1.0 µM (R209L, R209H, R209S, P208S, P208H, V173F).

The COU-3 substrate stock solution (20 mM) was prepared by dissolving the compound in DMSO. Working solutions were obtained by diluting the stock to a final concentration of 25–250 µM in PB buffer and stored shortly in amber vials prior to use.

Kinetic measurements were performed by monitoring fluorescence intensity changes using a CLARIOstar Plus plate reader (BMG Labtech, Germany) equipped with a linear variable filter (LVF) monochromator (configured to excitation/emission: 402–8/474–9 nm; dichroic filter: 437.8 nm) and dual injectors. Injector A (500 µL volume) was primed with the COU-3 substrate (50 µL stroke volume, 750 µL prime volume). Enzyme variants and buffer blanks were dispensed in triplicate (75 µL per well) into an OptiPlate 96-well polystyrene plate (PerkinElmer, catalogue no. 6005290). Following the injection of 75 µL of substrate, the plate was mixed using a double orbital shaking function for 5 seconds at 300 rpm. Fluorescence readings were recorded over 120 intervals (20 flashes per well, 33-second interval time). Instrument parameters, including gain (750), optical focal height (5.2 mm), and reaction temperature (25 °C), were held constant throughout all measurements to ensure consistent data across plates.

##### 6.2.1 Analysis and statistics of the steady-state kinetic data

The steady-state data were fit globally using the KinTek Explorer[9,10] (KinTek, USA), a dynamic kinetic simulation program that allowed multiple data sets to be fit simultaneously to a single model. Data fitting used numerical integration of rate equations from an input model searching a set of kinetic parameters, applying the Bulirsch–Stoer algorithm with an adaptive step size that produces a minimum χ2 value calculated by using nonlinear regression based on the Levenberg-Marquardt method. Residuals were normalised by the sigma value for each data point. The standard error (s.e.) was calculated from the covariance matrix during nonlinear regression. In addition to s.e. values, a more rigorous analysis of the variation in the kinetic parameters was accomplished by confidence contour analysis in FitSpace Explorer[9,10] (KinTek Corporation, USA). In these analyses, the lower and upper limits for each parameter were derived from the confidence contour obtained from setting $\chi^2$ threshold at 0.98. The steady-state model (*Figure S28*) was used to obtain the values of turnover number $k_{cat}$, Michaelis constant $K_M$ and equilibrium dissociation constant for substrate inhibitory complex $K_{SI}$. A conservative estimate of the diffusion-limited substrate binding rate constants, $k_1$ and $k_4$ (100 $\mu M^{-1} \cdot s^{-1}$), was fixed to approximate the rapid equilibrium assumption.

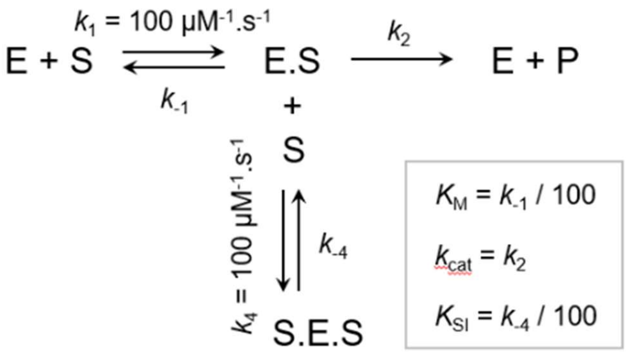

*Figure S28: The steady state model.*

##### 6.2.2 Steady-state kinetic results

*Table S15: Overview of the kinetic results obtained from the steady-state measurements with COU-3.*

| | | ***WT*** | ***I138N*** | ***P137S*** | ***P208S*** | ***R209L*** | ***V173F*** |
|---|---|---|---|---|---|---|---|
| *PARAMETER* | | Value ± s.e. | Value ± s.e. | Value ± s.e. | Value ± s.e. | Value ± s.e. | Value ± s.e. |
| $K_M$ | μM | 174 ± 7 | 35 ± 3 | 111 ± 10 | 69 ± 4 | 71 ± 11 | 86 ± 5 |
| $k_{cat}$ | $s^{-1}$ | 5.0 ± 0.2 | 6.1 ± 0.2 | 2.5 ± 0.2 | 0.115 ± 0.004 | 0.47 ± 0.04 | 0.70 ± 0.03 |
| $k_{cat}/K_M$ | $\mu M^{-1} \cdot s^{-1}$ | 0.029 ± 0.002 | 0.174 ± 0.016 | 0.022 ± 0.003 | 0.0017 ± 0.0001 | 0.007 ± 0.001 | 0.008 ± 0.001 |
| $K_{SI}$ | μM | 115 ± 5 | 87 ± 4 | 273 ± 34 | 395 ± 32 | 150 ± 10 | 139 ± 8 |

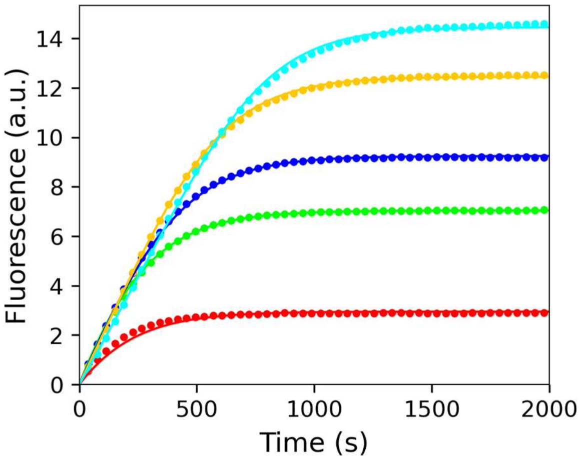

<u>LinB WT</u>

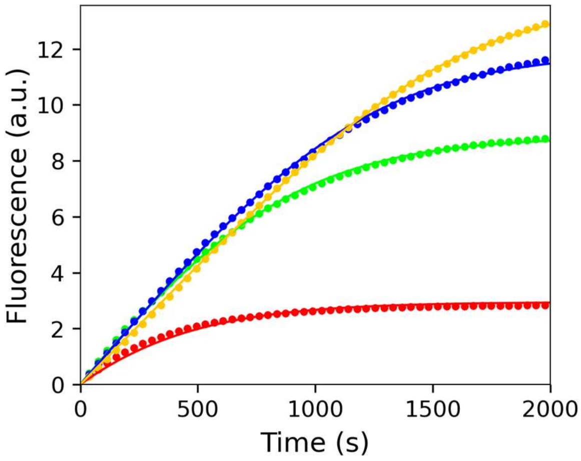

<u>LinB P137S</u>

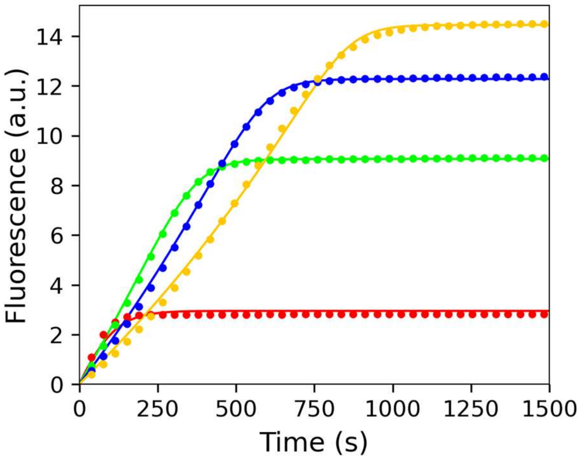

<u>LinB I138N</u>

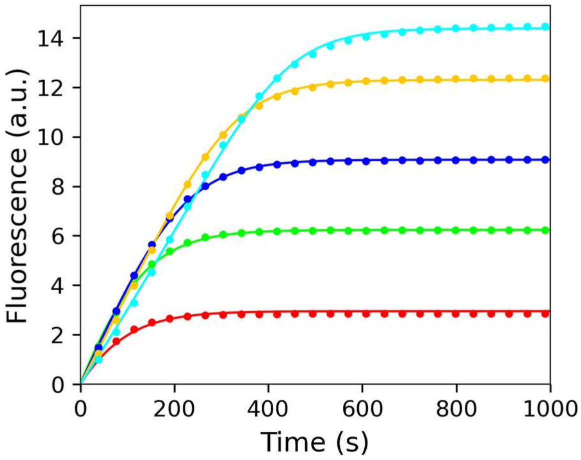

<u>LinB V173F</u>

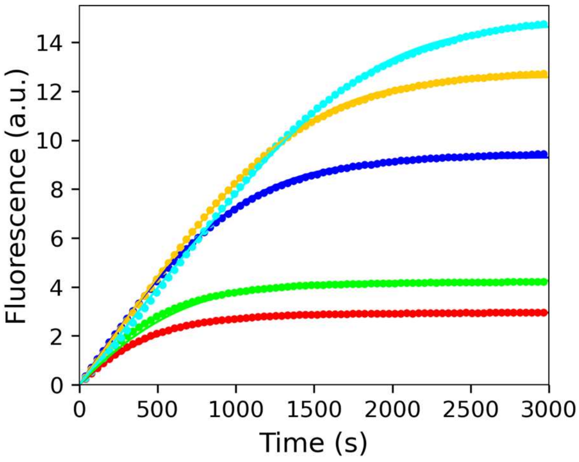

<u>LinB P208S</u>

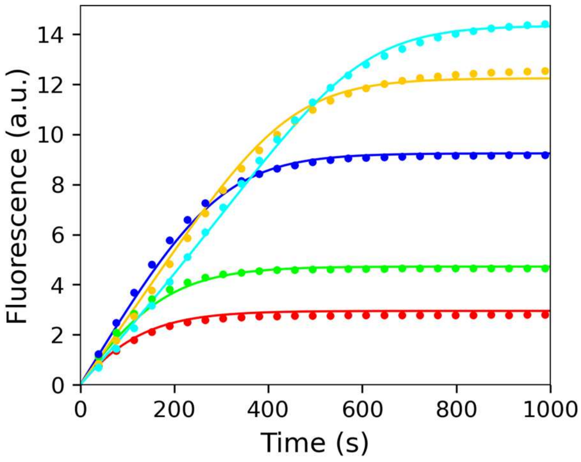

<u>LinB R209L</u>

*Figure S29: Steady kinetics of characterised LinB variants*

#### 6.3 Substrate specificity profiles

The droplets were generated using Mitos Dropix (Dolomite, UK). A custom sequence of droplets (150 nL aqueous phase, 300 nL oil spacing) was generated using negative pressure by a microfluidic pump (Chemyx, USA). The droplets were guided through a polythene tubing (Smith-Medical, UK) to the incubation chamber. Within the incubation chamber, the halogenated substrate was delivered to the droplets via a combination of microdialysis and partitioning between the oil (FC 40) and the aqueous phase. The reaction solution consisted of a weak buffer (1 mM HEPES, 20 mM $Na_2SO_4$, pH 8.0) and a complementary fluorescent indicator 8-hydroxypyrene-1,3,6-trisulfonic acid (50 μM HPTS). The fluorescence signal was obtained using an optical setup with an excitation laser (450 nm), a dichroic mirror with a cut-off at 490 nm to filter the excitation light, and a Si detector. Using a pH-based fluorescence assay, small pH changes were observed, enabling the monitoring of enzymatic activity. Reaction progress was analysed as an endpoint measurement recorded after 10 droplets/sample had passed through the incubation chamber. The reaction time was 4 min. The raw signal of every single measurement was first processed by the in-house LabView-based (National Instruments, USA) software MicroPEX Data Analyzer v4.1.2. The peaks were assigned to the particular sample, and the mean signal was calculated for them. The output XLS file gathering mean signal values for every sample type (calibration, enzyme activity, buffer, blank 3 buffer, and blank enzyme) for the dataset served as an input for the MatLab (Mathworks, USA) script to calculate specific activities using the same principle as for previous measurements[11,12]. The activities were classified as "not determined" whenever the measured product concentration was below the limit of detection (LOD – 3 times the standard deviation of the noise signal). Each substrate had a different calibration curve, so the LOD product concentration was in the range of 10-100 μM).

The matrix containing the activity data of 6 previously identified HLDs which were characterized for substrate specificity[11] at 25°C and the activities obtained for all variants in this study (all measured on MicroPEX) was analysed by PCA in MATLAB (MathWorks, USA) to uncover the relationships among individual HLDs (objects) based on their activities toward the set of halogenated substrates (variables). Two PCA models were constructed to visualise systematic trends in the dataset. The first one was based on the absolute values of specific activities, where the first principal component, which explained the highest variability in the data, ordered the enzymes according to their total activity. The second PCA was performed on the log-transformed data, after adding 1 to each specific activity to avoid taking the logarithm of zero. The resulting values were then divided by the sum of the enzyme's values. The first two principal components, which explained the highest variability in the data, were then plotted to identify trends in substrate specificity.

*Table S16: Results of the substrate specificity screening*

| No. | Name | LinBWT | P137S | I138N | V173F | P208S | R209L |
|---|---|---|---|---|---|---|---|
| **4** | **1-chlorobutane** | 0.019 | 0.019 | 0.011 | 0.017 | 0.008 | 0.013 |
| **6** | **1-chlorohexane** | 0.020 | 0.019 | 0.011 | 0.017 | 0.020 | 0.018 |
| **18** | **1-bromobutane** | 0.060 | 0.056 | 0.023 | 0.074 | 0.078 | 0.048 |
| **20** | **1-bromohexane** | 0.038 | 0.036 | 0.020 | 0.038 | 0.061 | 0.023 |
| **28** | **1-iodopropane** | 0.048 | 0.048 | 0.030 | 0.073 | 0.084 | 0.056 |
| **29** | **1-iodobutane** | 0.042 | 0.040 | 0.020 | 0.066 | 0.077 | 0.037 |
| **31** | **1-iodohexane** | 0.037 | 0.036 | 0.014 | 0.035 | 0.059 | 0.027 |
| **37** | **1,2-dichloroethane** | 0.004 | 0.004 | 0.000 | 0.002 | 0.000 | 0.001 |
| **38** | **1,3-dichloropropane** | 0.033 | 0.035 | 0.021 | 0.041 | 0.024 | 0.018 |
| **40** | **1,5-dichloropentane** | 0.021 | 0.020 | 0.012 | 0.025 | 0.024 | 0.022 |
| **47** | **1,2-dibromoethane** | 0.136 | 0.124 | 0.059 | 0.176 | 0.103 | 0.094 |
| **48** | **1,3-dibromopropane** | 0.109 | 0.099 | 0.048 | 0.116 | 0.153 | 0.071 |
| **52** | **1-bromo-3-chloropropane** | 0.070 | 0.069 | 0.055 | 0.084 | 0.077 | 0.076 |
| **54** | **1,3-diidopropane** | 0.036 | 0.035 | 0.018 | 0.061 | 0.074 | 0.049 |
| **67** | **1,2-dichloropropane** | 0.000 | 0.000 | 0.000 | 0.000 | 0.000 | 0.000 |
| **72** | **1,2-dibromopropane** | 0.062 | 0.058 | 0.024 | 0.051 | 0.007 | 0.031 |
| **80** | **1,2,3-trichloropropane** | 0.000 | 0.000 | 0.000 | 0.000 | 0.000 | 0.000 |
| **111** | **bis(2-chloroethyl)ether** | 0.017 | 0.015 | 0.005 | 0.006 | 0.004 | 0.009 |
| **115** | **chlorocyclohexane** | 0.005 | 0.002 | 0.000 | 0.000 | 0.000 | 0.000 |
| **117** | **bromocyclohexane** | 0.054 | 0.044 | 0.014 | 0.056 | 0.044 | 0.021 |
| **119** | **(1-bromomethyl)cyclohexane** | 0.007 | 0.006 | 0.001 | 0.015 | 0.005 | 0.003 |
| **137** | **1-bromo-2-chloroethane** | 0.072 | 0.082 | 0.055 | 0.111 | 0.080 | 0.074 |
| **138** | **chlorocyclopentane** | 0.021 | 0.020 | 0.009 | 0.018 | 0.016 | 0.015 |
| **141** | **4-bromobutyronitrile** | 0.066 | 0.057 | 0.029 | 0.071 | 0.029 | 0.036 |
| **154** | **1,2,3-tribromopropane** | 0.063 | 0.061 | 0.042 | 0.077 | 0.021 | 0.044 |
| **155** | **1,2-dibromo-3-chloropropane** | 0.041 | 0.032 | 0.021 | 0.055 | 0.003 | 0.014 |
| **209** | **3-chloro-2-methylpropene** | 0.080 | 0.076 | 0.050 | 0.073 | 0.062 | 0.066 |

#### 6.4 HDX-MS

All protein samples were diluted in phosphate buffer (17.2 mM $K_2HPO_4$, 2.8 mM $KH_2PO_4$, pH 8.0). Protein concentrations were measured by nanodrop before MS analysis, then manually diluted to 43 µM and kept at 25 °C in the HDX holder. Using a LEAP HDX robotic station, samples were mixed with $H_2O$/$D_2O$ buffer (pH 8.0, pD/pHread 7.6) to a final concentration of 2 µM and labelled with $D_2O$ for multiple time points (10–7200 s). After quenching with urea/glycine buffer (pH 2.3), samples were digested with Pepsin/Nepenthesin2, trapped/de-salted on a C18 microtrap, and separated on a Luna Omega Polar C18 column (2 °C) using a 6-min gradient. A fully deuterated control was prepared by digesting the protein, eluting in 98% acetonitrile, labelling peptides with $D_2O$ for 24 h, and quenching before MS detection.

LC-MS analysis was performed using an Agilent 1290 UHPLC coupled to a Bruker timsTOF mass spectrometer, with blanks between runs. Mapping samples were acquired in MS/MS mode with trapped ion mobility, while HDX samples were run in MS mode. All measurements were in technical triplicate. Data were searched with Mascot against a LinB protein database (1% FDR, 10 ppm precursor, 0.05 Da fragment tolerance). Duplicate peptides were removed, and MSTools was used for mapping and statistical plots. HDX data were processed with DeutEx (Bruker) and pyHDX for residue-level analysis of deuterium uptake.

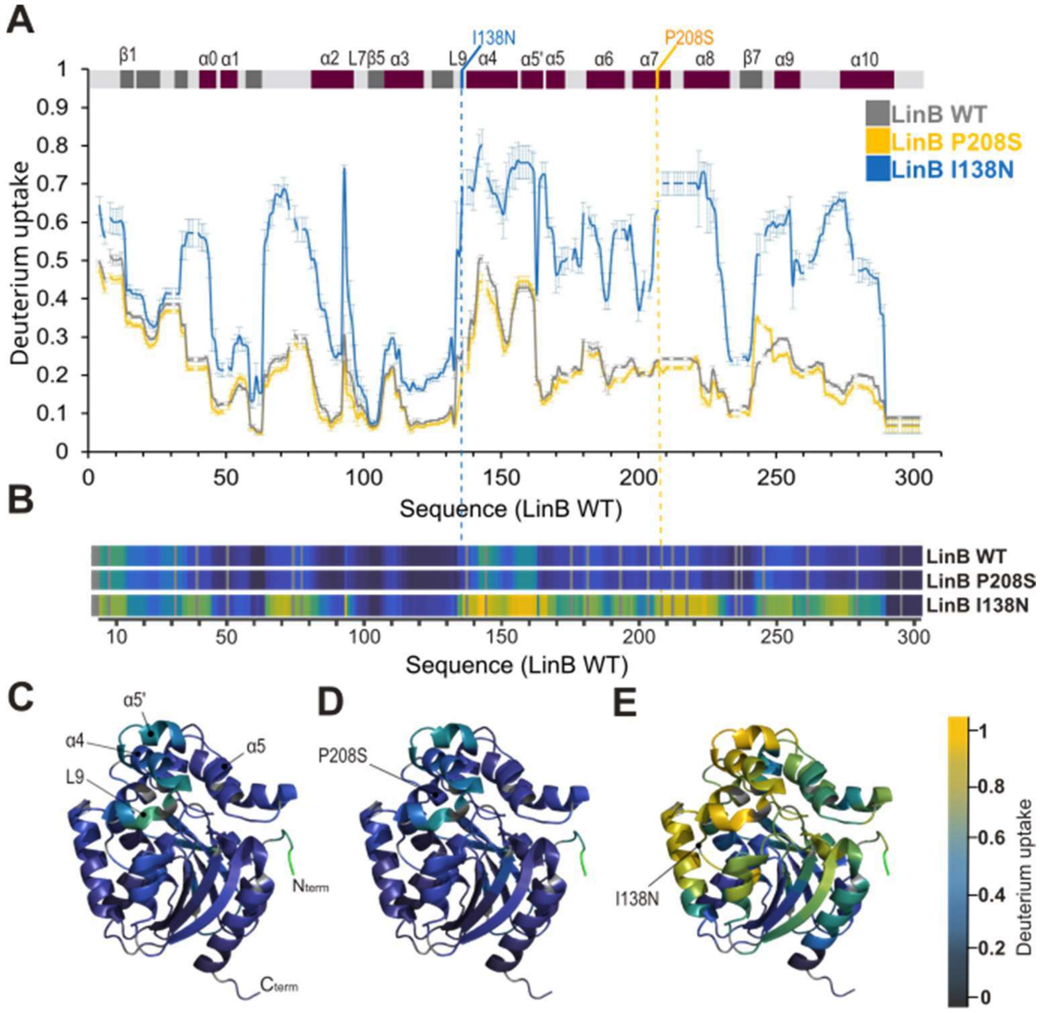

*Figure S30: HDX-MS results. (A) Average deuterium uptake of LinB WT (grey), LinB P208S (yellow), and LinB I138N (blue) after 10 s of $D_2O$ labelling. Mutation sites are indicated by dashed lines on the secondary structure. (B) Heatmap representation of D-uptake, with black indicating the lowest and yellow the highest uptake. (C-E) D-uptake values mapped onto the LinB WT structure (PDB: 1MJ5). LinB WT (C) and LinB P208S (D) display highly similar uptake profiles across the structure, whereas LinB I138N (E) shows markedly higher D-uptake in the cap-domain (residues 130-230), indicating increased flexibility due to the mutation.*

### 7 In Silico Hits Characterization

#### 7.1 Structure modelling of the proteins

The structure of LinB wild-type was obtained from the PDB[13] (ID: 1mj5). Three-dimensional models of the wild-type (for reference values) and the variants were obtained using MODELLER[14] with trivial alignments and following a minimum structural perturbation protocol established before[15]. 5 models were obtained per input sequence, and the one with better energetic assessment[16] (lower DOPE score). These models were used for the exclusive purpose of mapping predicted flexibility values.

#### 7.2 Flexibility Prediction

The sequences of the different LinB variants (LinB wild-type, P137S, I38N, V173F, P208S, P208H, R209H, R209S, and R209L) were used as input to run the inference algorithm of Flexpert Sequential module[17].

Flexpert v.1 (as downloaded from GitHub: https://github.com/KoubaPetr/Flexpert/releases/tag/v1) provides a per-residue flexibility prediction, ranging between 0 and 1, based on its training on molecular dynamics simulations data. Residues 2-296 were considered for this analysis.

##### 7.2.1 Flexibility mapping

In order to make the predicted values comparable among systems (and to make them comparable to other flexibility measures), all values obtained by Flexpert were internally standardised per system. Thus, each value from a LinB variant prediction series was transformed in the following fashion:

$$z = \frac{x - \hat{x}}{s} \qquad \text{Eq. 7.1}$$

where $z$ is the normalised per-residue value, $x$ is the predicted value (Flexpert score), $\hat{x}$ is the average of predicted values and $s$ is their standard deviation. These standardised values were mapped on LinB wild-type structure and represented both in colour (rainbow) and thickness (thicker representation as the value was larger). The comparison of the LinB wild-type flexibility values with those of the other variants, was achieved by calculating, per residue, the log2 ratio of the variant flexibility over the wild type one. This scale was used to represent the intensity and direction of change in colour on a red-white-blue scale, redder indicating positions that were predicted to be more flexible in the variant, bluer indicating positions that were predicted to be more rigid in the variant, and white representing no change. Finally, to facilitate the visualisation of the magnitude of change, the thickness was represented as proportional to the amount of change regardless of its direction (i.e., without considering if the change is towards more or less flexible), and the shorter branch (positive or negative range of values) of the distribution was re-scaled to match the extension of the larger one (i.e. data was transformed to have absolute maximum and minimum values equal). This further transformed data series was used to represent thickness in comparative flexibility mappings.

#### 7.3 Molecular modelling

##### 7.3.1 System preparation for docking and simulations

The three-dimensional structures of the proteins were prepared as described above. The crystallographic water molecules and ions were removed. The hydrogen atoms were predicted using the H++ server[18], calculated in implicit solvent at pH 7.5, 0.1 M salinity, internal dielectric constant of 10 and external of 80.

The three-dimensional structure of COU-3 was built and minimised using Avogadro[19]. The energy minimisation was performed by the Auto Optimisation Tool of Avogadro, using the UFF force field[20] with the steepest descent algorithm, and saved as a PDB file. The structure was submitted to optimisation and calculation of the partial atomic charges using Gaussian 09[21], with the DFT/B3LYP method and the 6-311G(d,p) basis set in vacuum. The antechamber module of AmberTools 16 was used[22] to extract the RESP charges of the ligand from the Gaussian output files and generate the PDB, MOL2, and PREPI files, and parmchk2 to generate the parameter modification file (FRCMOD).

##### 7.3.2 Molecular docking

The input files of the COU-3 ligand and receptors, in PDB format, were converted to the AutoDock Vina-compatible format PDBQT using MGLTools[23]. The non-polar hydrogens of the ligand and protein were preserved. The active site of the haloalkane dehalogenases was selected as the region of interest for the molecular docking performed by AutoDock Vina[24] 1.1.2. This region was represented by a box of 40 × 40 × 40 Å centred in the coordinates of the CG atom of the catalytic residue D108 of LinB. The exhaustiveness parameter was defined as 100, to increase the conformational search. The energy range was increased to 10 kcal·mol$^{-1}$, to obtain a higher number of binding poses, and the maximum number of modes saved was increased to 20. The docking results were visualised using PyMOL[25] 2.3.2.

The identification of pre-reactive conformations for the $S_N2$ reaction, also known as near-attack conformations (NACs), was based on the distances and angles between the nucleophile and the substrate atoms, according to Hur *et al.*[26]: the distance between the nearest carboxyl oxygen atoms of the nucleophile (D108-OD atoms) and the halogen-bound carbon atom must be $d_{O-C} \leq 3.41$ Å, and the angle formed by the oxygen, carbon and halide atoms must be $\alpha_{O-C-X} \geq 157$ °. As reported before[27], we also required at least a weak H-bonding between the reacting chlorine atom and the halide stabilising residues, defined by the distance between the halide and the indole polar hydrogen of W109 or the side chain NH hydrogen of N38 must be $d_{X-H} \leq 3.0$ Å. The reactive binding conformations obtained from docking were combined with the respective protein and saved as separate PDB files.

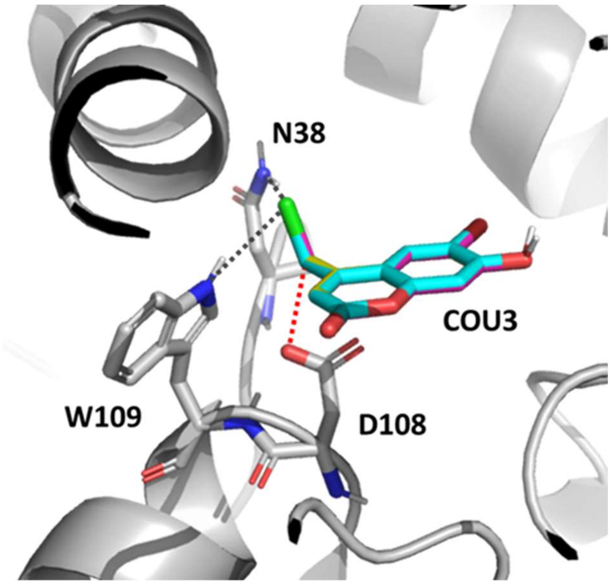

*Figure S31: Docking of COU3, the three LinB variants. The reactive binding conformations of COU3 docked to WT (magenta), I138N (yellow) and P208S (yellow) are superimposed almost perfectly. Only the structure of WT is represented as white cartoons, and the catalytic residues as sticks; the distance between the reacting carbon in the substrate and carboxylic oxygen is represented by the red dashed line, and the hydrogen bonds between the chlorine and the halide-stabilising residues N38 and W109 are represented by the black dotted lines.*

*Table S17: Docking binding affinities ($\Delta G_{bind}$, in kcal/mol) of the productive binding conformations.*

| Variant | Ranking of productive binding mode | $\Delta G_{bind}$ (kcal·mol$^{-1}$) |
|---|---|---|
| I138N | 3 | -6.3 |
| P208S | 3 | -6.2 |
| WT | 2 | -6.3 |

##### *7.3.3* Classical molecular dynamics

Different sets of molecular dynamics (MD) simulations were performed, with the free proteins in water, and with the proteins bound to the COU-3 substrate in reactive binding mode as obtained in the docking studies. The original waters in the crystallographic structure of LinB (PDB entry 1MJ5) were added to all the protein structures prepared as previously described, with and without COU-3. Any water molecule clashing with the protein or ligand atoms was removed. The *tLEaP* program of AmberTools[22] 16 was used to prepare the topology and coordinates files for all systems. For that, force field ff14SB[28] was specified to describe the protein, and the previously compiled PREPI and FRCMOD parameter files for the ligand. $Na^+$ and $Cl^-$ ions were added to neutralise the system and achieve a 0.10 M concentration of NaCl salt, and a truncated octahedron box of TIP3P[29] water molecules with the edges at least 10 Å away from the protein atoms was added.

The MD simulations were carried out with PMEMD.CUDA[30,31] module of AMBER[22] 16. In total, five minimisation steps and twelve steps of equilibration dynamics were performed prior to the production MD. The first four minimization steps, composed of 2,500 cycles of steepest descent followed by 7,500 cycles of conjugate gradient, were performed as follows: (i) in the first one, all the atoms of the protein and ligand were restrained with 500 kcal/mol·Å$^2$ harmonic force constant; (ii) in the following ones, only the backbone atoms of the protein and heavy atoms of the ligand were restrained, respectively, with 500, 125, and 25 kcal/mol·Å$^2$ force constant. A fifth minimisation step, composed of 5,000 cycles of steepest descent and 15,000 cycles of conjugate gradient, was performed without any restraints. The subsequent MD simulations employed periodic boundary conditions, the particle mesh Ewald method for treatment of the long-range interactions beyond the 10 Å cutoff[32], the SHAKE algorithm[33] to constrain the bonds involving the hydrogen atoms, the Berendsen barostat[34] at 1 bar, the Langevin thermostat with collision frequency 1.0 $ps^{-1}$, and a time step of 2 fs. Equilibration dynamics were performed in twelve steps: (i) 20 ps of gradual heating from 0 to 310 K, under constant volume, restraining the protein atoms and ligand with 200 kcal/mol·$Å^2$ harmonic force constant; (ii) ten MDs of 400 ps each, at constant pressure (1 bar) and constant temperature (310 K), with gradually decreasing the restraints on the backbone atoms of the protein and heavy atoms of the ligand with harmonic force constants of 150, 100, 75, 50, 25, 15, 10, 5, 1, and 0.5 kcal/mol·$Å^2$; (iii) 400 ps of unrestrained MD at the same conditions as the previous restrained MDs. The energy and coordinates were saved every 100 ps. The production MDs were run for 500 ns using the same settings employed in the last equilibration step. They were performed in two replicas for the free enzymes and four replicas for the enzymes with the bound substrate.

The replicas of each type were combined into a single trajectory using the *cpptraj*[35] module of AmberTools[22] 16, by stripping water molecules and ions, aligning each snapshot to the respective crystal structures by minimising the root-mean-square deviation (RMSD) of the backbone atoms, excluding the very flexible terminal residues (10 residues at both termini). The combined trajectories were visualised using PyMOL[25] 2.3.2 and VMD[36] 1.9.1, and they were further analysed with *cpptraj* to compute the B-factors by residue, using the respective backbone atoms, and the RMSD of Cα atoms.

The pre-reactive conformations (or near-attack conformations, *NACs*) were analysed in order to find the potentially configurations that may undergo the $S_N2$ reaction. *Cpptraj* was used to calculate distances and angles between the nucleophile and substrate atoms, and an in-house *Python* script evaluated them according to Hur *et al.*[26], as described above. For the P208S variant, we also computed the distances between the hydroxyl oxygen of S208 and each one of the side chain $NH_2$ hydrogen atoms of N38 for all the *NAC* complexes. Since those two hydrogen atoms are equivalent and can be interchanged, we identified the smallest of the two distances ($d_{O\cdots H}$) for all the *NACs* and calculated the respective histogram distribution.

##### 7.4 Access tunnel calculations

CAVER[37] 3.02 was used to calculate and cluster the tunnels in the crystal structures and during the MD simulations of the LinB variants. The tunnels were calculated for every 100 ps-spaced snapshots of the MDs. We specified a probe radius of 0.9 Å, a shell radius of 3 Å, and a shell depth of 4 Å. The starting point for the tunnel calculation was defined by the geometric centre of the carboxylic oxygen atoms of the catalytic D108 residue. The clustering was performed by the average-link hierarchical Murtagh algorithm[38] with a weighting coefficient of 1 and clustering threshold of 3.5 Å. Approximate clustering was allowed only when the total number of tunnels was higher than 20,000, and it was performed using 20 training clusters.

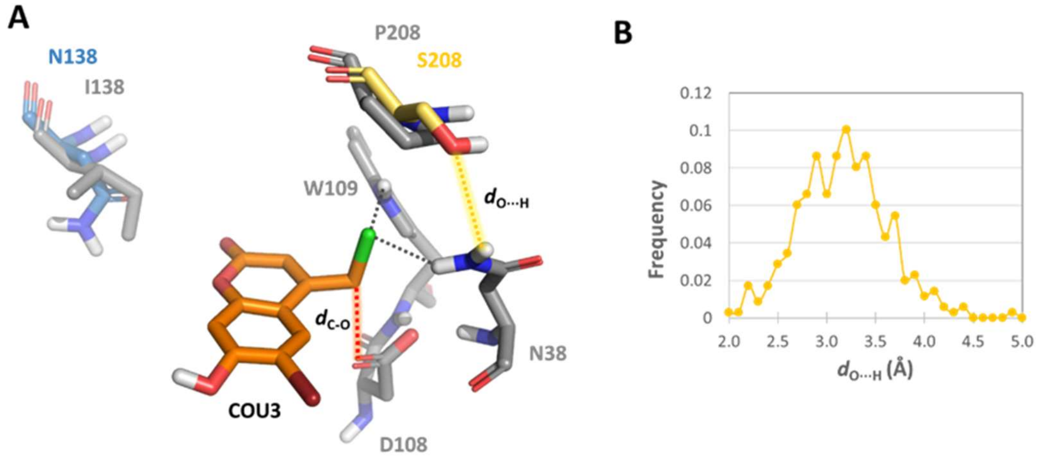

*Figure S32: Structure of the NAC complexes in the three variants. A) Representative NAC complex of COU3 (in orange) in the active site of LinB WT (in grey), displaying the catalytic and the mutated residues, superimposed with the NAC complexes of mutants I138N and P208S. For simplicity, only the mutated residues are represented for the mutant variants (residues N138, in blue, and S208 in yellow); the distance between the reacting carbon in the substrate and the carboxylic oxygen ($d_{C\text{-}O}$) is represented by the red dotted line, and the hydrogen bonds between the chlorine and the halide-stabilizing residues N38 and W109 are represented by the black dotted lines; the distance between the hydroxyl oxygen atom of S208 and the second N38 hydrogen atom ($d_{O\cdots H}$) is represented by the yellow dotted line. B) Histogram distribution of the $d_{O\cdots H}$ distance illustrated in (A) over all the NAC complexes detected in the MD simulations with the P208S variant. The generally small values of the $d_{O\cdots H}$ distance ($d_{O\cdots H} \leq 3.5$ Å in 74% of the NACs) suggests a strong interaction between the serine hydroxyl oxygen with one of the N38 hydrogen atoms. This may lead to a decrease in the stabilization provided by the other N38 hydrogen atom to the transition state, thus explaining the higher activation barrier calculated for of the $S_N2$ reaction in P208S.*

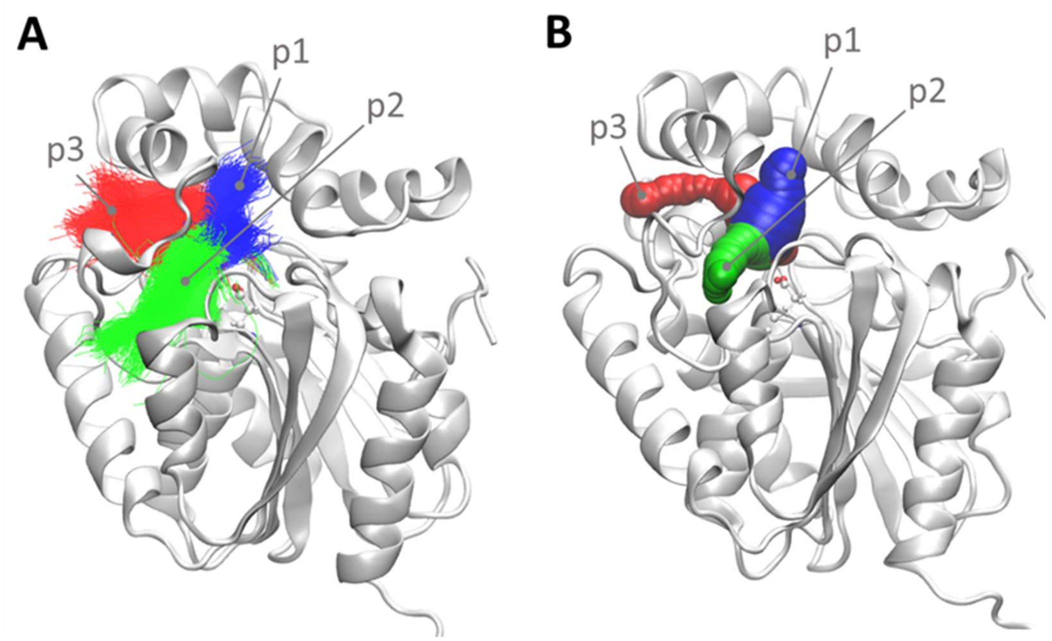

*Figure S33: The three main tunnels found in the MD simulations with the free enzymes (LinB WT is shown here). A) "timeless" tunnels, and B) tunnels in the snapshot at 410 ns. The timeless tunnels show the centreline of all the tunnels detected in all the snapshots of the MD, superimposed and coloured by the respective cluster colour (p1 blue, p2 green, and p3 red). The tunnels are named according to Chovancová et al.*[37,39]

*Table S18: Properties of the main tunnels during the MD simulations with the free LinB variants. **

| Variant | %Tunnels (≥0.9 Å) | %Open Tunnels (≥1.4 Å) | Avg. BRadius (≥0.9 Å) | Avg. BRadius Open (≥1.4 Å) | Max. BRadius | Avg. Length | Priority |
|---|---|---|---|---|---|---|---|
| **Tunnel p1** | | | | | | | |
| **I138N** | 99.5 | 85.6 | 1.54 ± 0.13 | 1.57 ± 0.08 | 1.74 | 11.6 ± 1.6 | 0.76091 |
| **P208S** | 99.5 | 80.5 | 1.51 ± 0.14 | 1.56 ± 0.08 | 1.74 | 12.2 ± 1.4 | 0.73197 |
| **WT** | 98.9 | 77.3 | 1.50 ± 0.16 | 1.57 ± 0.08 | 1.75 | 12.1 ± 1.6 | 0.72893 |
| **Tunnel p2** | | | | | | | |
| **I138N** | 53.8 | 16.0 | 1.26 ± 0.21 | 1.51 ± 0.08 | 1.72 | 17.7 ± 2.4 | 0.30958 |
| **P208S** | 31.4 | 4.5 | 1.14 ± 0.20 | 1.49 ± 0.07 | 1.71 | 21.0 ± 2.7 | 0.15187 |
| **WT** | 58.6 | 20.3 | 1.29 ± 0.21 | 1.52 ± 0.08 | 1.73 | 18.1 ± 2.9 | 0.33917 |
| **Tunnel p3** | | | | | | | |
| **I138N** | 12.8 | 0.3 | 1.02 ± 0.12 | 1.46 ± 0.06 | 1.61 | 19.7 ± 2.3 | 0.05524 |
| **P208S** | 7.0 | 0.02 | 0.97 ± 0.08 | 1.53 ± 0.10 | 1.64 | 19.8 ± 2.6 | 0.02657 |
| **WT** | 16.8 | 0.4 | 1.03 ± 0.13 | 1.47 ± 0.05 | 1.60 | 19.8 ± 2.3 | 0.07273 |

*"%Tunnels" is the ratio of snapshots where the tunnel was detected (with bottleneck radius ≥ 0.9 Å); "%Open Tunnels" is the ratio of snapshots where the tunnel tunnels was open (with bottleneck radius ≥ 1.4 Å); "Avg. BRadius" (in Å) is the average bottleneck radius considering all the tunnels detected; "Avg. BRadius Open" (in Å) is the average bottleneck radius considering only the open tunnels (with bottleneck radius ≥ 1.4 Å); "Max. BRadius" (in Å) is the maximum bottleneck radius observed over the entire trajectory; "Avg. Length" (in Å) is the average length of the tunnel, considering all the tunnels

detected; "Priority" is a parameter defined in CAVER 3.02[36] that combines the average radius, length, curvature, and the frequency with which the specific tunnel were detected during the simulation, and is used to rank the tunnels as their potential likelihood of being used for molecular transport. The average values are reported with the respective standard deviations.

##### 7.4.1 QM/MM adiabatic mapping

To evaluate the energetics of the $S_N2$ reaction of the probes with the different enzymes and estimate the respective energy barrier, $\Delta G^{\ddagger}$, we studied the potential energy surface (PES) along the reaction coordinate using a combined quantum mechanics/molecular mechanics (QM/MM) approach[40,41]. For that, the pre-reactive complexes, previously identified in the MD simulations of the enzymes with bound COU-3 in their active sites, were subjected to QM/MM calculations. The quality of these complexes was assessed by the distance between the reacting atoms ($d_{O-C}$), and only the 100 best complexes with the lowest distances were analyzed by QM/MM adiabatic mapping. The topology of each structure was prepared by *tLEaP* module using the ff14SB force field for the proteins and the PREPI parameters for the ligand. The complexes were minimised in vacuum (*igb=6*). Five rounds of optimization, each one consisting of 500 cycles of steepest descent followed by 500 conjugate gradient cycles, were performed as: (i) one step with all heavy atoms restrained with 500 kcal/mol·Å$^2$ harmonic force constant, and (ii) four steps with decreasing restraints on the protein backbone atoms with 500, 125, 25 and 1 kcal/mol·Å$^2$ force constant. Adiabatic mapping along the reaction coordinate was performed by the sander module of AMBER 14[42]. The QM part of the system contained the probe molecule, the side-chains of the halide-stabilising residues (N38 and W109), and the catalytic aspartate (D108) and had charge -1. The semi-empirical PM6 Hamiltonian was used to treat the QM part of the system[43] and the ff14SB force field to treat the MM part. The QM/MM boundary was treated through explicit link atoms, and the cut-off for the QM/MM charge interactions was set to 999 Å. The backbone was constrained with a force constant of 1.0 kcal/mol·Å$^2$. The reaction coordinate was defined as the distance between the nearest OD atom of D106 and the C-atom of the probe under attack. The tracking along the reaction coordinate was performed in decrements of 0.05 Å, each involving 1,000 minimisation steps of the limited-memory Broyden-Fletcher-Goldfarb-Shanno quasi-Newton algorithm[44]. The energy barrier $\Delta G^{\ddagger}$ was calculated as the difference between the lowest value of the potential energy at the ground state and the energy at the transition state.

##### 7.4.2 Adaptive steered molecular dynamics

Adaptive steered molecular dynamics (ASMD) simulations were performed for each LinB variant with COU-3 located outside of tunnel p1 and tunnel p2. For guidance, the tunnels were calculated on LinB (PDB entry 1MJ5) with the CAVER[37] 3.02 plugin in PyMOL 2.3.2. COU-3 was manually placed outside the proteins, in front of the respective tunnel mouth, using PyMOL, ensuring a distance between the ligand and the protein atoms of at least 5 Å. The crystallographic waters from LinB (PDB entry 1MJ5) were added into each system, removing any overlapping molecules, and the resulting complex was saved as a PDB file. tLEaP was used for every system as described above, for adding ions to achieve a 0.10 M concentration of NaCl, adding a truncated octahedron box of TIP3P water molecules, and generating the topology and coordinates files.

Equilibration was performed with PMEMD.CUDA[30,31], as described above for the classical MDs, except for the final unrestrained equilibration step (in total, 5 minimisations and 11 equilibration dynamics were performed). The unrestrained step was omitted because it can result in a stochastic displacement of the ligand, potentially introducing systematic bias into all subsequent simulation replicas. Excluding this step ensured that the replicas were initiated from equivalent, unbiased configurations.

The binding of COU-3 to the active site of each enzyme, travelling through the different tunnels, was studied with ASMD. In the ASMD method, a constant external force is applied between two selected atoms, and the reaction coordinate is advanced in discrete stages by incrementally decreasing (or increasing) their distance. At each stage, multiple parallel simulations are initiated from the same initial state. Their work profiles are then analysed to compute the Jarzynski average[45,46], and the trajectory with a work value closest to this average provides the starting configuration for the subsequent stage. Here, we used most of the default values recommended in the AMBER tutorial and the ASMD publication[47]. To steer the ASMD, we selected the distance between the reacting carbon in the substrate and the Asp108-Cγ atom of the enzyme (part of the reacting carboxylic group of the nucleophile). The initial distance between the two atoms was measured in the last snapshot from the last equilibration step, and the final distance was defined as 2 Å. The simulations were run with 25 parallel MDs, steered by 1.5 Å stages of distance decrements, with a velocity of 10 Å/ns, and a force of 7.2 N. The rest of the MD parameters were set as described for the unrestricted classical MD simulations. For each ligand-binding simulation, five replicas were performed, and the Jarzynski average of the potential of mean force (PMF) was calculated at each distance point, with the respective standard error, using an in-house Python script.

*Table S19: PMF values at the final distance of 4 Å, obtained by ASMD for tunnels p1 and p2, and respective statistical significance. **

| Variant | $PMF_f$ | $DPMF_f$ (Mut-WT) |
|---|---|---|
| **Tunnel p1** | | |
| **I138N** | 20.64 ± 0.62 | 0.41 |
| **P208S** | 22.49 ± 0.33 | 2.26 |
| **WT** | 20.22 ± 0.35 | - |
| **Tunnel p2** | | |
| **I138N** | 17.48 ± 0.61 | -0.89 |
| **P208S** | 21.17 ± 0.61 | 2.80 |
| **WT** | 18.37 ± 0.50 | - |

| ***P-values*** | **I138N** | **P208S** | **WT** |
|---|---|---|---|
| **I138N** | - | 0.030 | 0.5715 |
| **P208S** | 0.030 | - | 0.0015 |
| **WT** | 0.5715 | 0.0015 | - |
| **I138N** | - | 0.0027 | 0.2919 |
| **P208S** | 0.0027 | - | 0.0075 |
| **WT** | 0.2919 | 0.0075 | - |

*Color-coding:*

Very statistically significant.

Statistically significant.

Not statistically significant.

*"$PMF_f$" (in kcal/mol) is the total relative free energy required for the substrate to travel from the bulk solvent through the respective tunnel, reaching the active site with final distance of 4 Å, as the Jarzynski average and the respective SEM; the *p-values* were obtained from unpaired *t*-tests using the mean, SEM and the number of samples ($N = 5$).
